# Frequent origins of traumatic insemination involve convergent shifts in sperm and genital morphology

**DOI:** 10.1101/2021.02.16.431427

**Authors:** Jeremias N. Brand, Luke J. Harmon, Lukas Schärer

## Abstract

Traumatic insemination is a mating behaviour during which the (sperm) donor uses a traumatic intromittent organ to inject an ejaculate through the epidermis of the (sperm) recipient, thereby frequently circumventing the female genitalia. Traumatic insemination occurs widely across animals, but the frequency of its evolution, the intermediate stages via which it originates, and the morphological changes that such shifts involve remain poorly understood. Based on observations in 145 species of the free-living flatworm genus *Macrostomum*, we identify at least nine independent evolutionary origins of traumatic insemination from reciprocal copulation, but no clear indication of reversals. These origins involve convergent shifts in multivariate morphospace of male and female reproductive traits, suggesting that traumatic insemination has a canalising effect on morphology. Signatures of male-female coevolution across the genus indicate that sexual selection and sexual conflict drive the evolution of traumatic insemination, because it allows donors to bypass postcopulatory control mechanisms of recipients.

## Introduction

The sexes frequently show differences in mating propensity because male fertility (i.e. fertilised egg production) is often limited by the number of matings a male achieves, while female fertility is often limited by the amount of resources a female invests into eggs and offspring [1–3]. The resulting conflict over mating rate has far-reaching consequences, often resulting in “Darwinian sex roles” with choosy females and eager males [4]. Females may benefit from choice by selecting males based on genetic compatibility, genetic quality [5] and/or direct benefits (e.g. nuptial gifts [6]). Indeed, evidence for female choice is widespread and there are many species where females mate multiply, suggesting polyandry may indeed result in such benefits [7]. However, females may also mate multiply as a result of male harassment, and while that could be costly to females, resisting male harassment might be even costlier [7, 8]. Costly harassment is expected to arise frequently, since female choice necessarily goes against the rejected males’ interests [9], potentially leading to sexually antagonistic coevolution between male persistence and female resistance traits [8, 10, 11].

In polyandrous species, sexual selection and sexual conflict continue after copulation through intricate interactions of the female genital tract with the male intromittent organs and the received ejaculate [12–14]. Female genitalia might exert postcopulatory control through differential sperm storage, sperm ejection or sperm digestion, thus applying selective filters on male genital and ejaculate traits. In analogy to the precopulatory conflict, it is then possible for traits in males to arise that attempt to bypass or influence the female-choice and resistance mechanisms, again resulting in sexually antagonistic coevolution [12–14].

Such coevolution can drive the emergence of male traits that inflict considerable harm on females [11, 15, 16]. A striking example that implicates such harm is traumatic insemination, which occurs in some internally fertilising species and involves the infliction of a wound to the female’s integument through which the male then transfers its ejaculate [17]. Since traumatic insemination occurs in both gonochoristic (separate-sexed) and hermaphroditic species [17], we in the following use the more general terms (sperm) donor and (sperm) recipient to refer to the two sexual roles, with no loss of generality [18].

Although traumatic insemination often results in costs to recipients [15, 17, 19–22], it has evolved repeatedly across animals [17]. And while natural selection might play a role in some taxa—especially the endoparasitic Strepsiptera [23, 24]—it likely often evolves due to sexual selection and sexual conflict. Specifically, traumatic insemination can enable donors to enforce copulation and thus minimise the control that the recipient could otherwise exert over mating [15]. And it may also allow the donor to bypass the recipient’s genitalia, by depositing sperm either closer to the site of fertilisation [24, 25] or even directly within the relevant tissue [15, 26], thus likely reducing the recipient’s ability to control the fate of the received ejaculate [12, 17]. In this view, traumatic insemination allows the donor to bypass the influence of the recipient’s sexually antagonistic choice and resistance mechanisms, temporarily gaining an advantage in the coevolutionary chase.

However, since conflicts persist under traumatic insemination, we expect selection to then act on traits that allow the recipient to regain control over mating and/or the fate of the received ejaculate. For example, some species of bed bugs have evolved what is considered a secondary vagina, a structure shown to reduce the costs incurred due to traumatic insemination [19, 27]. But even without the emergence of new organs, recipients could evolve behavioural or physiological responses to avoid traumatic insemination (such as parrying strikes during penis fencing in polyclad flatworms [28]) or to manipulate and control the hypodermically received ejaculate (e.g. similar to sperm digestion in copulating species [29–31]).

Besides bypassing recipient choice and resistance mechanisms, traumatic insemination could also evolve due to sperm competition, since in many internally fertilising species sperm of unrelated donors compete within the female genital tract for fertilisation of the recipient’s eggs [32]. In this context, traumatic insemination might allow donors to avoid sperm competition and prevent competing donors from removing their previously donated sperm, resulting in paternity benefits [17]. Indeed, traumatic insemination seems to affect sperm competition in a family of spiders, where sperm precedence is biased towards the first male in a species with traumatic insemination, while it is biased towards the second male in its non-traumatically mating relatives [17, 23, 33]. In contrast, traumatic insemination is associated with last male precedence in one species of bed bug [26], so its effects on sperm competition might depend on a species’ morphology and ecology.

Traumatic insemination might evolve more frequently in hermaphrodites due to sexual conflict over the mating roles [12, 18, 34–36]. In general, and analogous to the situation outlined for gonochorists [1], a hermaphrodite already carrying enough received sperm to fertilise its own eggs might gain little from additional matings as a recipient, while it could still gain additional fertilisations by acting as a donor [12]. It is thus likely that, on average, individual hermaphrodites show a preference for sperm donation [12, 18, 34–36] and this rationale is supported by several laboratory studies [35, 37, 38]. Traumatic insemination then potentially allows individuals to attempt unilateral enforcement of donation while avoiding receipt. Additionally, hermaphrodites may engage in harmful matings more readily, because any fitness costs an individual incurs as a recipient may be partially compensated by fitness benefits it incurs as a donor [34, 39]. Indeed, 11 out of 23 well-supported independent origins of traumatic insemination occurred in hermaphrodites [17], even though hermaphrodites amount to only ~6% of animals [40]. Hermaphrodites are thus ideal study organisms for investigations of traumatic insemination, since—while it has been studied in some charismatic systems [15, 25, 28, 41–44]—we currently still know little about the frequency and consequences of its evolution [17, 20, 22].

Here we present comparative work on the evolution of traumatic insemination across the genus *Macrostomum*, a species-rich taxon of hermaphroditic free-living flatworms. In *Macrostomum*, traumatic insemination is called hypodermic insemination (HI), since in several species the donor uses a needle-like stylet (Figure 1) to inject sperm through the mating partner’s epidermis and sperm then move through the recipient’s body to the site of fertilisation [45–47]. Injected sperm can often be observed inside the parenchymal tissues of these highly transparent animals [45–48], making it feasible to screen a large number of species for convergent evolution of HI. And while we here present evidence that not all traumatically mating *Macrostomum* species may inject sperm through the external epidermis, we nevertheless use the term HI for consistency with previous literature.

**Figure 1.**
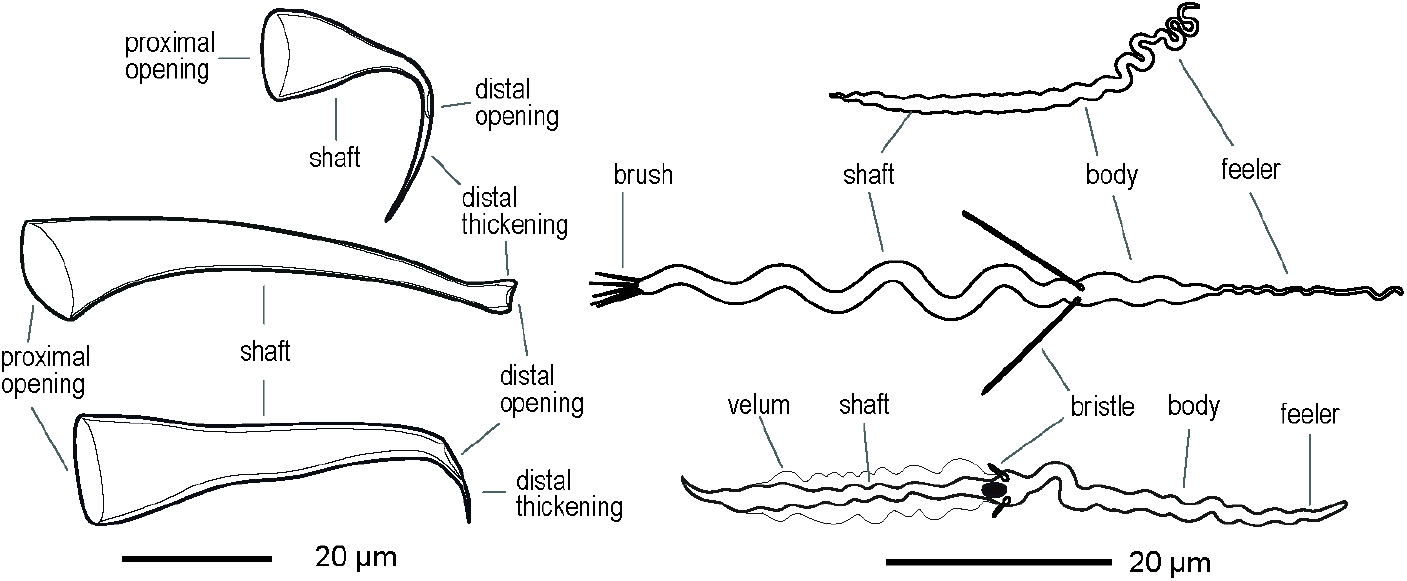
Representative drawings of the morphology of the stylet (male intromittent organ) (left) and the sperm (right) of three *Macrostomum* species. (Top) *M*. sp. 92, a hypodermically mating species from the hypodermic clade, with a typical needle-like stylet and a simple sperm morphology. (Middle) The well-studied model *M. lignano* with the typical morphology for reciprocally mating species, showing a stylet with blunt distal thickenings and a complex sperm with an anterior feeler, two stiff lateral bristles, and a terminal brush. (Bottom) *M*. sp. 9 representing one of the convergent origins of hypodermic insemination in the reciprocal clade, showing a stylet with a highly asymmetric and sharp distal thickening and sperm with reduced sperm bristles, no brush, but a thin velum along the shaft. Note that, given the striking diversity across the *Macrostomum* genus, it is not possible to clearly delimit all the sperm traits originally defined in *M. lignano* in some of the species.

The genus comprises two phylogenetically well-separated clades [49], a “hypodermic clade” thought to exclusively mate through HI and a “reciprocal clade” primarily mating reciprocally (called Clade 1 and 2, respectively, in [45]), with the latter containing a convergent origin of HI in *M. hystrix* [45]. During reciprocal copulation two worms insert their—often relatively blunt—stylet (Figure 1) via their partner’s female genital opening into the female sperm storage organ, the female antrum (further called antrum), so that both can donate and receive sperm in the same mating [50]. Many reciprocally copulating species perform a postcopulatory suck behaviour, where worms place their mouth over their own female genital opening and suck, presumably in an attempt to remove components of the received ejaculate from their antrum [45, 50–53]. This ejaculate removal could target manipulative seminal fluids, since the ejaculate of the model species *M. lignano,* contains substances affecting the mating partner’s propensity to perform the suck behaviour [54, 55]. Alternatively, the suck behaviour could also reduce the number of stored sperm (e.g. to lower the risk of polyspermy), constitute a form of cryptic female choice (e.g. to favour donors of higher quality), and/or represent a resistance trait in sexual conflict over mating roles (i.e. to undo unwanted sperm receipt) [45, 51].

If the suck behaviour is a recipient resistance trait, we might expect the evolution of donor persistence traits, potentially leading to antagonistic coevolution [8]. Indeed, the sperm of reciprocally copulating species generally have a thin anterior feeler and two stiff lateral bristles that could represent such persistence traits (Figure 1), serving to anchor the sperm in the antrum to prevent removal during the suck behaviour [45, 51]. In contrast, sperm of species with HI (i.e. the hypodermic clade and *M. hystrix*) lack these bristles and have a simplified morphology, presumably because they no longer need to resist the suck behaviour [45, 51], which has so far never been observed in species with HI. These sperm may instead be adapted to efficiently move through the partner’s tissues (Figure 1), and one such adaptation could hypothetically also include a reduced sperm size [45]. Moreover, while species with reciprocal copulation have an antrum with a thickened epithelium, those with HI have a simple antrum, presumably because it no longer interacts with the donor’s stylet and sperm, and instead is used for egg-laying only [45]. Based on these findings, the observed adaptations to reciprocal copulation and HI have been described as the reciprocal and hypodermic mating syndrome, respectively, since they each constitute specific combinations of morphological (sperm, stylet and antrum) and behavioural traits [45].

If HI indeed represents a resolution of sexual conflict over mating roles, then we would expect it to evolve frequently. But it is currently unclear whether HI has convergently arisen more than once within the reciprocal clade. It is also unclear if such transitions are reversible or if the emergence of HI alters the coevolutionary dynamics between donor and recipient, so that species cannot readily revert to reciprocal copulation. Here we collate morphological information on 145 *Macrostomum* species to identify additional independent origins of HI and to quantitatively assess convergent changes in both sperm design, and in male and female genital morphology that accompany its evolution, taking advantage of a recent large-scale phylogenomic analysis of the genus [49]. Using ancestral state reconstruction, we further ask whether species can revert to reciprocal copulation once HI has arisen. Moreover, if sexually antagonistic coevolution drives the emergence of HI, we expect signatures of coevolution. We thus test for covariation between male and female genital traits and survey the genus for novel resistance and persistence traits.

## Results

### Species collected and phylogenetics

We used phylogenetic information and operational species assignments that we recently generated by integrating morphological and transcriptome data, supplemented with partial *28S rRNA* sequences and information from the literature [49]. We used a phylogeny including 145 species (C-IQ-TREE, shown in Figure 2), but to assess how sensitive our analyses are to phylogenetic uncertainty, we also performed all analyses on two alternative phylogenies, including only 98 species with full transcriptome information (H-IQ-TREE and H-ExaBayes). Since all results were quantitatively similar and qualitatively identical, we focus on the C-IQ-TREE results, but report the additional analyses in the supplementary files. We collected morphological data on up to eight quantitative traits from 1442 specimens and scored eleven categorical traits on a per-species basis (see Materials and Methods, and SI Morphology for details; see Table S1 for sample sizes, Table S2 for all measurements, and Table S3 for species mean values).

**Figure 2.**
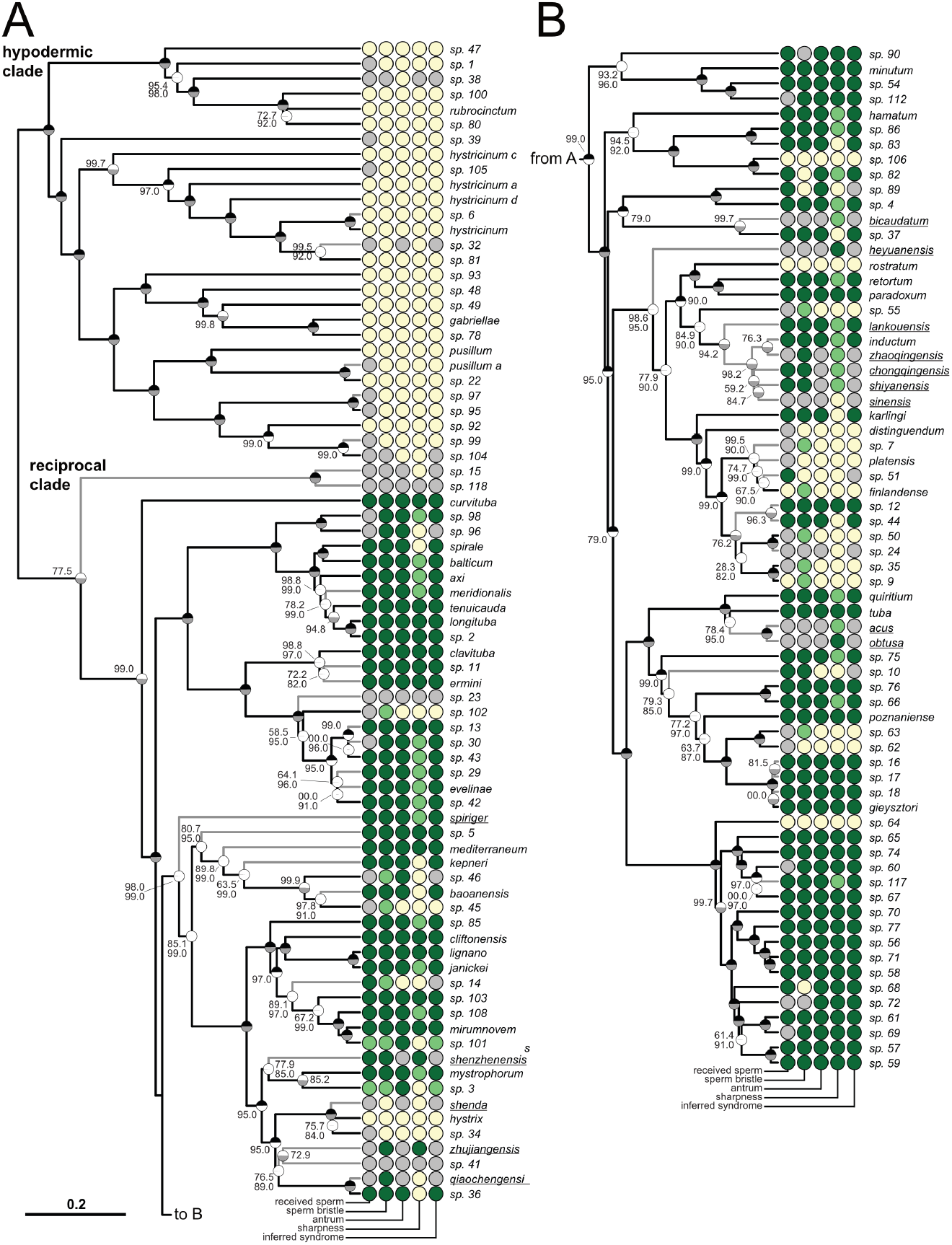
Phylogeny of the genus *Macrostomum*, showing the states of five reproductive traits. The ultrametric phylogeny (C-IQ-TREE) includes all 145 species from [49]. Branch supports are ultrafast bootstraps (top, black if 100) and approximate likelihood ratio tests (bottom, grey if 100). Species without available transcriptomes that were added based on a *28S rRNA* fragment are indicated with grey branches. Underlined species names indicate that the trait scoring is based on information from the literature. Two phylogenetically well-separated clades the “hypodermic clade” thought to exclusively mate through hypodermic insemination (HI) and the “reciprocal clade” primarily mating reciprocally are labelled in A. Columns indicate the states of five reproductive traits from light to dark (i.e. yellow, light green and dark green for trinary states; or yellow and dark green for binary states; grey indicates missing data): received sperm location (hypodermic, both, in antrum), sperm bristle state (absent, reduced, present), antrum state (simple, thickened), sharpness of stylet (sharp, neutral, blunt), inferred mating syndrome (hypodermic, intermediate, reciprocal). The phylogeny is split into two parts (A and B) for visualisation. See also Figure SX combining this figure with drawings of stylet and sperm morphology available from [49].

### Frequent origins of hypodermic insemination

We inferred the number of convergent transitions to HI using ancestral state reconstruction (ASR) of several reproductive traits (Figure 2). Because scoring the received sperm location involves observation of sperm within the recipient’s tissue, it provides the most direct evidence for HI (Table 1 and SI Morphology). However, observation of injected sperm in field-collected specimens can be challenging, especially in species with low investment into sperm production, thus reducing sample size. Since tests of correlated evolution revealed strong associations of hypodermic sperm with absent/reduced sperm bristles and a thin antrum (see the next section), we therefore also performed ASR using the sperm bristle state and antrum state as proxies for HI. Finally, we also performed ASR on the inferred mating syndrome, which represents a synthesis of all available information (Table 2 and Materials and Methods). We performed ASR, with all traits scored as binary and, where appropriate, also as trinary, to test if HI could evolve via an intermediate state (see Materials and Methods).

**Table 1.** Ancestral state reconstructions of reproductive traits, including received sperm location, sperm bristle state, antrum state, and inferred mating syndrome. A range of MK-models (ER: equal rate, SYM: symmetrical rate, ORD-Dollo: ordered model without gains once the trait is in state 0, Dollo: model without gains, ORD: ordered model, ARD: all rates different) were compared based on their AIC weights. For each trait the model with the highest AICc weight (AICcw) is shown in bold type, but we estimated the number of transitions between the states using stochastic character mapping with 1000 posterior samples for all models with an AICc weight >0.15. Given are the average number of transitions and the 2.5% and 97.5% quantiles in brackets. Results are based on the C-IQ-TREE phylogeny. For the quantitatively similar results with the H-IQ-TREE and H-ExaBayes phylogenies see Table S4.

| Reproductive trait | model | dfs | log lik | AICc | ΔAICc | AICcw | N<sub>Species</sub> | N<sub>Changes</sub> | 0 → 1 | 1 → 0 | 1 → 2 | 2 → 1 | 0 → 2 | 2 → 0 |
| --- | --- | --- | --- | --- | --- | --- | --- | --- | --- | --- | --- | --- | --- | --- |
| Rec. sperm location | ER | 1 | -43.3 | 88.6 | 10.1 | 0.004 |  |  |  |  |  |  |  |  |
| 0: hypodermic only | SYM | 3 | -40.9 | 88.1 | 9.6 | 0.005 |  |  |  |  |  |  |  |  |
| 1: hypodermic & in antrum | **ORD-Dollo** | **3** | **-36.1** | **78.5** | **0** | **0.579** | **102** | **21.2 (16, 32)** | - | **8.0 (7, 13)** | **4.0 (0, 11)** | **9.3 (7, 13)** | - | - |
|  | Dollo | 4 | -36.1 | 80.6 | 2.2 | 0.196 | 102 | 10.8 (9, 17) | - | 0.8 (0, 4) | 0.9 (0, 4) | 2.6 (2, 5) | - | 6.5 (4, 8) |
| 2: in antrum only | ORD | 4 | -36.1 | 80.6 | 2.2 | 0.196 | 102 | 23.4 (16, 38) | 0.7 (0, 4) | 8.4 (6, 13) | 4.8 (0, 15) | 9.6 (7, 15) | - | - |
|  | ARD | 6 | -36.1 | 85.1 | 6.6 | 0.021 |  |  |  |  |  |  |  |  |
| Rec. sperm location | ER | 1 | -36.8 | 75.6 | 3.1 | 0.135 |  |  |  |  |  |  |  |  |
| 0: hypodermic | **Dollo** | **1** | **-35.2** | **72.5** | **0** | **0.638** | **102** | **9.3 (9, 11)** | - | **9.3 (9, 11)** |  |  |  |  |
| 1: in antrum only | ARD | 2 | -35.2 | 74.5 | 2.1 | 0.227 | 102 | 9.9 (9, 14) | 0.9 (0, 4) | 9.0 (8, 12) |  |  |  |  |
| Sperm bristle state | ER | 1 | -82.9 | 167.8 | 27.2 | 0 |  |  |  |  |  |  |  |  |
| 0: absent | SYM | 3 | -81.7 | 169.5 | 28.9 | 0 |  |  |  |  |  |  |  |  |
| 1: reduced | **ORD-Dollo** | **3** | **-67.2** | **140.6** | **0** | **0.578** | **131** | **36.3 (29, 48)** | - | **12.2 (11, 17)** | **6.7 (1, 16)** | **17.5 (14, 23)** | - | - |
| 2: present | Dollo | 4 | -67.2 | 142.7 | 2.1 | 0.2 | 131 | 29.3 (22, 41) | - | 7.1 (3, 13) | 5.3 (1, 13) | 12.4 (8, 18) | - | 4.6 (0, 8) |
|  | ORD | 4 | -67.2 | 142.7 | 2.1 | 0.199 | 131 | 37.1 (29, 50) | 0.8 (0, 5) | 13.0 (10, 20) | 6.0 (1, 14) | 17.3 (14, 22) | - | - |
|  | ARD | 6 | -67.2 | 147.1 | 6.5 | 0.023 |  |  |  |  |  |  |  |  |
| Sperm bristle state | ER | 1 | -57.6 | 117.3 | 5.7 | 0.038 |  |  |  |  |  |  |  |  |
| 0: absent or reduced | **Dollo** | **1** | **-54.8** | **111.6** | **0** | **0.635** | **131** | **18.8 (18, 22)** | - | **18.8 (18, 22)** |  |  |  |  |
| 1: present | ARD | 2 | -54.4 | 112.9 | 1.3 | 0.327 | 131 | 20.5 (17, 29) | 2.7 (0, 9) | 17.8 (15, 23) |  |  |  |  |
| Antrum state | ER | 1 | -48.1 | 98.2 | 3.8 | 0.086 |  |  |  |  |  |  |  |  |
| 0: simple | **Dollo** | **1** | **-46.2** | **94.4** | **0** | **0.577** | **127** | **14.7 (14, 17)** | - | **14.7 (14, 17)** |  |  |  |  |
| 1: thickened | ARD | 2 | -45.7 | 95.5 | 1.1 | 0.337 | 127 | 15.2 (13, 21) | 2.1 (0, 6) | 13.1 (11, 16) |  |  |  |  |
| Inf. mating syndrome | ER | 1 | -59.3 | 120.6 | 16.2 | 0 |  |  |  |  |  |  |  |  |
| 0: hypodermic | SYM | 3 | -54.4 | 114.9 | 10.4 | 0.003 |  |  |  |  |  |  |  |  |
| 1: intermediate | **ORD-Dollo** | **3** | **-49.1** | **104.5** | **0** | **0.578** | **119** | **34.1 (27, 47)** | - | **13.7 (12, 18)** | **7.1 (2, 16)** | **13.2 (10, 19)** | - | - |
| 2: reciprocal | Dollo | 4 | -49.1 | 106.6 | 2.1 | 0.199 | 119 | 16.6 (14, 25) | - | 0.7 (0, 5) | 1.1 (0, 6) | 2.8 (2, 6) | - | 11.9 (9, 15) |
|  | ORD | 4 | -49.1 | 106.6 | 2.1 | 0.198 | 119 | 35.4 (27, 50) | 1.0 (0, 5) | 14.3 (11, 20) | 7.2 (2, 16) | 12.9 (9, 18) | - | - |
|  | ARD | 6 | -49.1 | 111 | 6.5 | 0.022 |  |  |  |  |  |  |  |  |
| Inf. mating syndrome | ER | 1 | -49.4 | 100.9 | 14.675 | 0.001 |  |  |  |  |  |  |  |  |
| 0: hypodermic & intermediate | **Dollo** | **1** | **-42.1** | **86.26** | **0** | **0.997** | **119** | **14.7 (14, 17)** | - | **14.7 (14, 17)** |  |  |  |  |
| 1: reciprocal | ARD | 2 | -47.3 | 98.68 | 12.421 | 0.002 |  |  |  |  |  |  |  |  |

**Table 2.** Assignment of the inferred mating syndrome based on different reproductive traits. Species were assigned to an inferred mating syndrome based on the location of received sperm in the body (antrum, in the antrum only; hypodermic, hypodermic only; both, in the antrum and hypodermic; NA, no observation), the sperm bristle state (absent, reduced or present), the antrum state (simple or thickened), and the shape of the distal thickening of the stylet (sharp or blunt). 26 species with either not enough (22 species) or contradictory (four species) information were not assigned to a syndrome. Note, that all 24 species with only hypodermic sperm had the same morphological states, but this was not a condition for their assignment (hence the brackets). Similarly, all 69 species assigned to the reciprocal mating syndrome had a thickened antrum, but this was also not a condition for their assignment. See also Materials and Methods.

| Syndrome | Received sperm location |  |  |  | Morphology |  |  | N |
| --- | --- | --- | --- | --- | --- | --- | --- | --- |
|  | Antrum | Hypodermic | Both | NA | Sperm bristle | Antrum | Stylet |  |
| Hypodermic |  | 24 |  |  | (Reduced/absent) | (Simple) | (Sharp) | 24 |
| Hypodermic |  |  |  | 18 | Reduced/absent | Simple | Sharp | 18 |
| Intermediate |  |  | 2 |  | Reduced | Thickened | Sharp | 2 |
| Reciprocal | 61 |  |  | 6 | Any state | (Thickened) | Blunt | 67 |
| Reciprocal | 8 |  |  |  | Present | (Thickened) | Sharp | 8 |
| Unclear | 7 |  |  | 19 | Other combinations |  |  | 26 |

All reconstructions indicated frequent origins of HI (Table 1 and Figure S1). In all analyses with trinary states, an ordered transition model without gains once traits have been lost (ORD-Dollo) was preferred, and in all analyses with binary states, a model without gains (Dollo) was preferred. However, other models, including some permitting gains, also received at least some support (Table 1). ASR of trinary states inferred frequent transitions to the intermediate state, which were driven by the ordered model’s requirements to transition through it. These transitions were often placed along internal branches of the phylogeny, primarily within the clade containing *M. finlandense* (Figure 2B, middle), which contains several species with reduced or absent states and, nested within them, two species with present states (*M*. sp. 12 and *M*. sp. 44, with received sperm in the antrum, long bristles, and assigned to the reciprocal mating syndrome; Figure S1 A, C, F). To represent this diversity, Figure S2 combines our Figure 2 with drawings of stylet and sperm morphology available from [49]. We estimated a lower bound for the number of transitions by requiring an origin of the derived state to be separated by other such origins via nodes with a >95% posterior probability of having the ancestral state. Applying this rule to traits scored as binary, we find nine transitions to hypodermic received sperm, 17 losses/reductions of sperm bristles, 13 simplifications of the antrum, and 13 transitions to the hypodermic or intermediate mating syndrome (see red stars and numbers in Figure S1). Moreover, these lower-bound estimates were slightly lower for trinary states. Finally, we found qualitatively very similar results on the other two phylogenies included, albeit, since they contain fewer species, showing somewhat lower numbers of transitions (Table S4).

### Correlated evolution

We performed tests of correlated evolution to ask if the numerous convergent changes in received sperm location, sperm bristle state and antrum state are evolutionarily dependent. We found strong support for correlated evolution of received sperm location with both sperm bristle state and antrum state (Figure 3A+B). This supports previous findings that HI is associated with changes in sperm design and antrum simplification [45]. Therefore, when observations of received sperm are missing, both sperm bristle state and antrum state are likely good proxies for the mating syndrome. We expand on the earlier analyses by also providing evidence for the correlated evolution between the sperm bristle state and antrum state (Figure 3C), which was implied in [45], but not formally tested. Across the board, we find substantially stronger support for correlated evolution than [45], with Bayes factors that are ~7-fold larger, reflecting the larger sample sizes and the larger number of transitions. Moreover, these analyses were robust with respect to the phylogeny and the priors used (see SI Correlated evolution).

**Figure 3.**
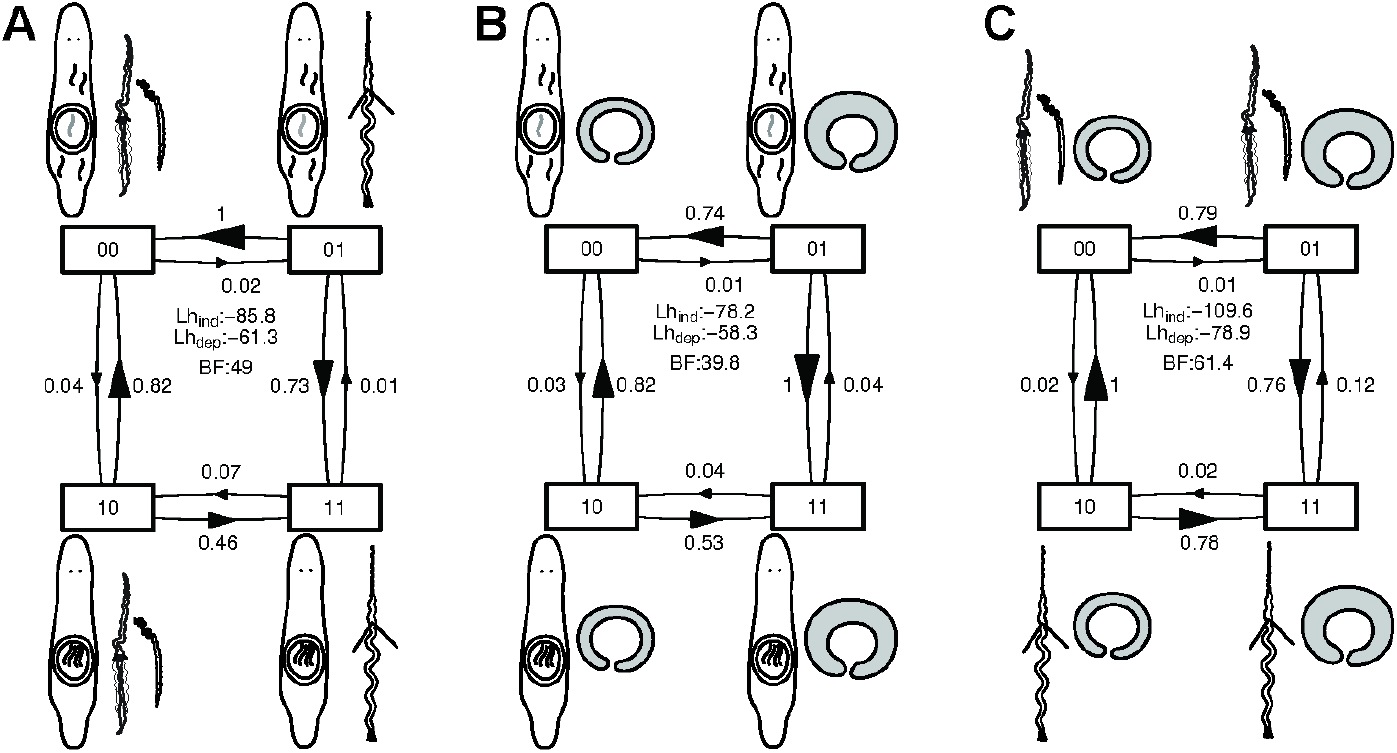
Results of correlated evolution analysis between (A) received sperm location and sperm bristle state, (B) received sperm location and antrum state, and (C) sperm bristle state and antrum state. Shown is the transition matrix for the dependent model from BayesTraits analysis, which was always preferred over the independent model. Transition rates are scaled so that the largest is unity (and arrow sizes are proportional). Also given are the likelihoods of the independent (Lh_ind_) and dependent (Lh_dep_) models, and the resulting Bayes factors (BF). An exponential prior and the C-IQ-TREE phylogeny was used for the results shown here. See SI Correlated evolution for runs with other priors (uniform and reversible-jump hyperprior) and other phylogenies (H-IQ-TREE and H-ExaBayes), which show qualitatively similar results.

### Convergence in morphospace

Next, we used phylogenetically corrected principal component analysis (pPCA) to investigate if these convergent transitions to HI also coincided with changes in a larger set of reproductive traits (see SI Morphology). The first two principal components, PC1 and PC2, captured nearly half of the variation in the reproductive traits (Figure 4A), followed by additional principal components with relatively small contributions (Table S5). Specifically, PC1 captured a change in stylet phenotype, with larger values indicating species with longer, more curved stylets, that are distally more symmetric and less sharp (Figure 4A). Larger values of PC1 also indicated both longer sperm and bristles, and an increased probability for the sperm to carry a brush. Finally, high values of PC1 indicated a thickened antrum with a more pronounced cellular valve, and a more complex internal structure. In comparison, PC2 had a less clear interpretation, with high values indicating larger species with larger proximal and distal stylet openings.

**Figure 4.**
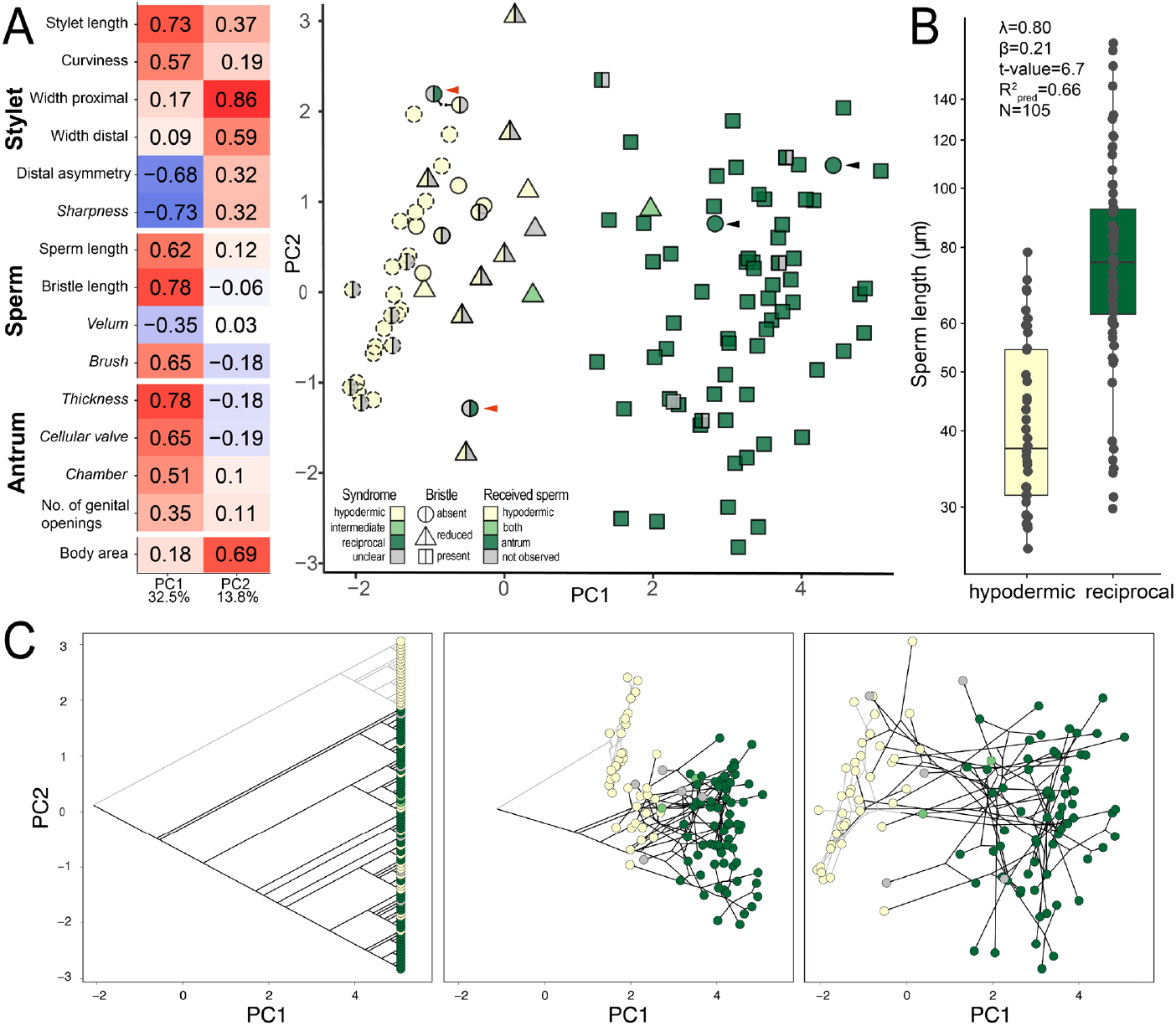
Results of a phylogenetically corrected principal component (pPCA) analysis of the measured quantitative (regular) and categorical (italics) reproductive traits (A,C) and PGLS regression of sperm length dependent on the inferred mating syndrome (B). (A) Left: Loadings of PC1 and PC2, with the percentage of variance explained at the bottom. Right: 2D morphospace defined by PC1 and PC2. As indicated by the legend, the shape represents the sperm bristle state, while the colours represent the inferred mating syndrome (left side) and the received sperm location (right side). All species from the hypodermic clade are outlined with stippled lines. Red arrowheads indicate two species (*Macrostomum* sp. 51 and *M*. sp. 89) that cluster closely with species assigned to the hypodermic mating syndrome, but in which we observed received sperm in the antrum. Black arrowheads indicate two species (*M*. sp. 68 and *M*. sp. 82) assigned to the reciprocal syndrome, which have no discernible sperm bristles (see also Figure S4). (B) Sperm length of species dependent on the inferred mating syndrome. Values are slightly jittered in the x direction, and the y-axis is on a log-scale. Within the panel the main results of PGLS analysis are given, with the slope being significant at p <0.001. Results shown here are based on C-IQ-TREE, while detailed results including analyses with other phylogenies (H-IQ-TREE and H-ExaBayes) are in Table S6. (C) The phylogenetic relationships of all species included in the pPCA analysis is represented in the left panel, and the right panel illustrates how species assigned to the hypodermic mating syndrome cluster in morphospace (as also seen in A). Edges of the hypodermic clade are printed in grey to aid in visualisation. The central panel shows an intermediate state in a phytools [105] phylomorphospace animation converting the left to the right panels, see Figure S3.

Species in the hypodermic clade (stippled outlines) had similar values in PC1 and mainly differed in PC2 (Figure 4A). Interestingly, species from the reciprocal clade (solid outlines) that we had assigned to the hypodermic mating syndrome (left yellow) grouped closely with the species in the hypodermic clade, indicating striking convergence in morphospace concerning stylet, sperm and antrum morphology (Figure 4C and Figure S3). PC1 further separated species based on the received sperm location, with hypodermic received sperm (right yellow) only found in species with low PC1, indicating that low PC1 captures a morphology necessary for HI. Almost all species with reduced (triangles) or absent (circles) sperm bristles grouped closely together in PC1, with the notable exception of *M*. sp. 68 and *M*. sp. 82 (black arrowheads), which cluster together with other species that we assigned to the reciprocal mating syndrome. We observed sperm in the antrum of both species (i.e. in 2 of 7 specimens in *M*. sp. 68 and 16 of 21 specimens in *M*. sp. 82) and the antrum is similar in both, with a long muscular duct that performs a 90° turn towards the anterior before it enters a second chamber that is strongly muscular (Figure S4). Moreover, both species have a similar L-shaped stylet with a blunt tip, which makes it unlikely that they mate through HI.

### Hypodermic insemination and sperm length

In addition to the changes in sperm design mentioned above, we tested whether HI is associated with a change in sperm length using phylogenetic least squares (PGLS) regression. We used received sperm location, sperm bristle state, antrum state, and the inferred mating syndrome as predictors and the log10-transformed sperm length as the response variable. In all cases, the states that indicate the reciprocal mating syndrome were associated with longer sperm, with the largest effect for the antrum state, followed by the inferred mating syndrome (Figure 4B and Figure S5). This is reasonable, since the bristle type falsely classified *M*. sp. 68 and *M*. sp. 82 as hypodermically mating, while the received sperm location and inferred mating syndrome analyses had slightly lower samples sizes. The predictive value of these PGLS models was generally high, indicating that a large proportion of the variation in sperm length is explained by the phylogeny and these mating syndrome indicators of the (Table S6). Note that despite these strong associations, there is considerable overlap in sperm length between the species exhibiting the different states, with some species with the reciprocal mating syndrome having short sperm (Figure 4B, Table S3) and an overall 6.7-fold variation in sperm length across all species (with means ranging from 25.6 to 173.1 μm).

### Male-female coevolution

To investigate coevolution between male and female genital traits, we independently summarised five male and four female genital traits using pPCA. Stylet PC1 was positively loaded with stylet length and the width of the distal opening, and it was negatively loaded with distal asymmetry (Figure 5A; Table S7). Therefore, high values of Stylet PC1 represent a more elongate stylet with a wider and less sharp distal opening. Antrum PC1 was positively loaded with all input variables (Figure 5B), meaning that large values represent more complex female genitalia. A PGLS regression of Stylet PC1 on Antrum PC1 across all species revealed a significant positive relationship (Figure 5C). This relationship closely matches the loadings on PC1 in the pPCA analysis of all reproductive traits (Figure 4A) and could be driven by the simple antra in hypodermically mating species. Therefore, we restricted the analysis to include only species assigned to the reciprocal mating syndrome and could confirm the positive relationship between Stylet PC1 and Antrum PC1 (Figure 5C).

**Figure 5.**
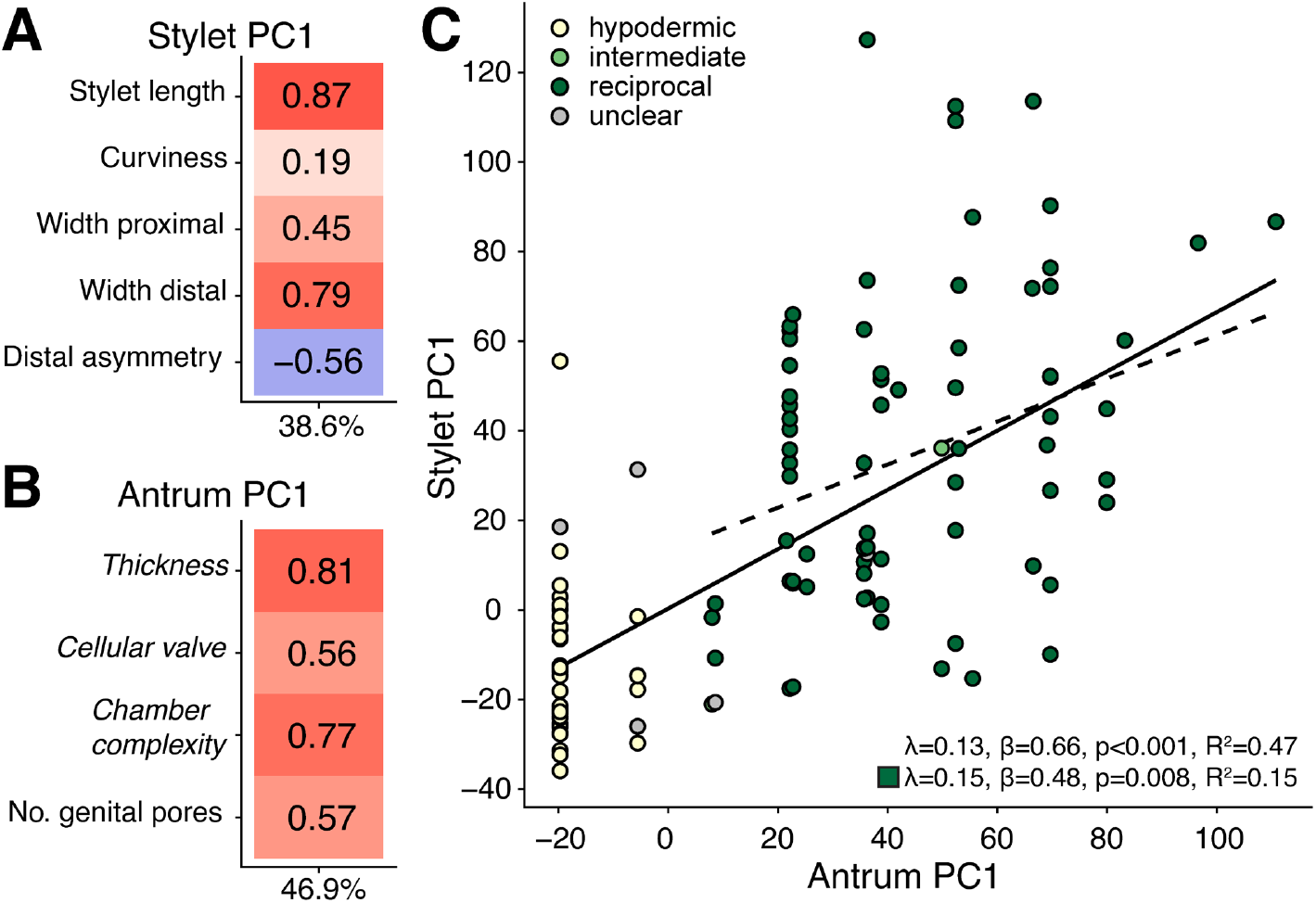
Phylogenetically corrected principal component analyses (pPCA) of stylet and antrum traits, and evidence for male-female coevolution. (A and B) Loadings of Stylet PC1 and Antrum PC1, with the percentage of variance explained at the bottom, for the stylet traits (A) and antrum (B) traits, respectively (categorical reproductive traits are in italics). (C) Results from PGLS regression of Stylet PC1 on Antrum PC1 from (A and B). Regression was performed across all species (solid line, upper statistics) and restricted to species of the reciprocal mating syndrome (dashed line, lower statistics). Dot colour indicates the inferred mating syndrome that the species are assigned to: hypodermic (yellow), intermediate (light green), reciprocal (green), and unclear (grey). Results based on C-IQ-TREE phylogeny, detailed results including analysis with other phylogenies (H-IQ-TREE and H-ExaBayes) are in Table S7.

## Discussion

Across the genus *Macrostomum*, hypodermic insemination (HI) has evolved independently at least 9 times as assessed by the location of received sperm, and at least 13 times based on the more inclusive inferred mating syndrome. According to [17], 12 and 11 origins of traumatic insemination have been found in gonochorists and hermaphrodites, respectively (including the two cases previously documented in *Macrostomum*). This means that, based on an investigation of a single free-living flatworm genus, we here approximately double the number of documented origins of HI among hermaphrodites. Since free-living flatworm diversity remains notoriously understudied [56], and since most of the collected *Macrostomum* species are likely undescribed [49], it seems probable that we have not even documented all convergent origins of HI within *Macrostomum*. Moreover, three additional origins of traumatic insemination occur in the genus’ parent group (Macrostomorpha [57]), suggesting that traumatic insemination may evolve frequently there and potentially also in other groups of flatworms.

Interestingly, we find no clear evidence for reversals back to reciprocal mating once HI has arisen, since the Dollo models were preferred in all our ancestral state reconstructions (although alternative models in some cases also received some support, Table 1). Reciprocal copulation is the ancestral state of the reciprocal clade, but the state of the most recent common ancestor of the genus is less certain (Figure S1), allowing for either a gain or a loss. Similarly, the clade containing *M. finlandense* (Figure 2B, middle) could contain either two independent losses or a single loss with a gain in *M*. sp. 12 and *M*. sp. 44 (Figure S1). We think the former is more likely since shifts to HI might be predominantly unidirectional. Specifically, once copulation is lost, a reversal would presumably require both mating partners to again coordinate reciprocal mating behaviour. Additionally, a subsequent antrum simplification could further hinder reversals, since copulating species have traits that presumably reduce the risk of injury (e.g. thickened antrum epithelia and stylets with blunt distal thickenings). In their absence, occasional reciprocal copulations could result in high fitness costs for both partners. In contrast, HI is presumably often unilateral, thus not requiring both partners to cooperate.

Our detailed observations of received sperm in both the antrum and embedded inside the recipient’s tissues led us to categorise two species, *M*. sp. 3 and *M*. sp. 101, as intermediate between the mating syndromes (light green triangles in Figure 4). These observations suggest that evolutionary transitions to HI occur through initial traumatic injection of sperm during canonical reciprocal copulation, possibly as a result of accidental hypodermic sperm transfer during copulatory wounding (for a more detailed discussion, including drawings and images on where we observed sperm in these two species, see SI Pathways to hypodermic insemination). Once HI has evolved, recipients in some organisms evolve secondary female genitalia to avoid costs of wounding and regain control over the received ejaculate [17]. Why this has not occurred in *Macrostomum* is unclear, but it might imply that costs of HI are generally low (possibly due to the striking regeneration ability of these flatworms [58]) or that the location of insemination is too variable for the evolution of a localised novel organ.

An earlier study [45] classified a needle-like stylet and shorter, simpler sperm as adaptations for HI, and observed an associated simplification of the antrum, presumably because it is only used for egg-laying in hypodermically mating species. Their test of correlated evolution of discrete antrum, stylet and sperm traits supported this hypothesis [45], but included only two independent origins of HI, with one containing a single species (*M. hystrix*). While tests of correlated evolution supposedly correct for phylogenetic dependencies, it was recently outlined that they can support the dependent model of evolution even with only a single (unreplicated) origin of the trait states in question ([59, 60]). Thus, while the previous findings supported correlated evolution, that evidence was not as decisive as these tests may have suggested. By sampling more convergent events, we here remedy this limitation, substantially raising our confidence in a causal link between sperm bristle and antrum state with HI (and this convergence has taxonomic implications, as we outline in more detail in SI Taxonomy). The increased sample size also enabled the pPCA analysis showing that species with HI indeed have similar values of PC1, with such values corresponding tightly to the mating syndromes described by [45] (note our slight adjustment of their definitions due to incomplete behavioural observations, Table 2), suggesting they truly are adaptations to HI. The striking convergent evolution clearly suggests that the origin of HI canalises taxa both morphologically and behaviourally.

Besides HI and its associated traits, another example of convergent evolution in *Macrostomum* is the origin of a second female genital opening, which the phylogeny suggests has evolved at least four times independently within the genus (for a more detailed discussion see SI Female openings). In all species, the novel second opening is associated with a muscular bursa that could possibly allow cryptic female choice by ejecting sperm via muscular contractions. Such contraction occurs during the suck behaviour in *M. hamatum*, a species with only a single opening, where sperm can be observed to be partially pushed out from the antrum even before the worm places its mouth on the female genital opening (P. Singh, pers. comm.).

Frequent convergent evolution of potential resistance traits, like a second female genital opening, or of alternative strategies, like HI, bolsters the interpretation that they resolve sexual conflict over mating rate, mating role or both [12, 18, 28, 34, 45, 51]. HI likely is an alternative strategy in an ongoing evolutionary chase between donor and recipient, with donor persistence traits, such as complex sperm with bristles [45] and manipulative seminal fluids [54, 55], and recipient resistance traits, such as the suck behaviour [50] and complex female genitalia [50, 51], engaged in constant antagonistic coevolution [12, 17, 20, 34]. We find evidence for such male-female genital coevolution, both across all species and within the species assigned to the reciprocal mating syndrome (Figure 5). Our findings agree with other work on hermaphrodites (e.g. [41, 61, 62]) and contribute to a growing body of evidence that male-female coevolution is common in both hermaphrodites and gonochorists [63–66]. Genital coevolution is not only expected due to sexual conflict but also predicted in the context of sexual selection. Under the sexual selection perspective, we expect coevolution due to cryptic female choice, where the recipient will choose based on genital traits of the donor [67]. Donors are therefore selected to closely match their genital morphology to the selection criteria of the recipient. Under both views, the respective selective optima of these traits might differ between species, driving diversification and speciation [68, 69]. Our findings clearly document a dynamic evolutionary history of male-female coevolution driving frequent innovations of sexual traits. Traumatic insemination allows donors to (temporarily) overcome pre- and postcopulatory choice and/or resistance mechanisms of the recipient, and results in striking convergence in morphospace (Figure 4).

One very striking convergent change we observe is that HI leads to a reduction in sperm size, for which we see three possible explanations. First, because HI avoids the recipient’s genitalia, it probably reduces the scope for both cryptic female choice by the recipient (e.g. via the suck behaviour) and sperm displacement/removal by competing donors. These postcopulatory mechanisms can introduce skews in sperm representation [70–73], which can result in lower levels of sperm competition compared to a “fair-raffle” type sperm competition when sperm mix more freely [32, 74, 75]. In this case, HI could increase sperm competition and, if sperm size trades-off with sperm number [74–76], select for smaller sperm [45, 77]. Second, *Macrostomum* sperm is large compared to the antrum, and therefore intimately interacts with its epithelium, often being partially embedded in the cellular valve with the feeler [45, 50, 51, 78], and sperm is also in close contact with rival sperm when recipients mate multiply [79, 80]. Under such conditions of high sperm density, i.e. when sperm displacement is likely (e.g. [81–83]), sperm are predicted to be bigger compared to species in which the sperm storage organ is substantially larger than the sperm [84, 85]. While under HI sperm still intimately interact with the partner’s tissue, the “storage organ” could now include the recipient’s whole body, reducing sperm-sperm interaction and decreasing positive selection on sperm size. Third, if small sperm can move more efficiently through the dense parenchymal tissue of the mating partner then natural selection could favour a decrease in sperm size [45]. Little is known about sperm movement within the recipient’s tissues, but it seems analogous to the undulating movement of the sperm body observed within the antrum [86]. These explanations are not mutually exclusive, and their relative importance might depend on the physiology, morphology, and ecology of each species.

Besides changes in sperm length, we confirm that the evolution of HI involves the convergent reduction and loss of sperm bristles [45, 49] (Figure SX), and document hypodermic received sperm in species with reduced bristles, indicating that HI can precede the complete loss of bristles. The preference for an ordered model in the ASR even suggests that transitions via an intermediate state may be the rule. It is unclear if bristle loss is adaptive or whether it occurs due to relaxed selection and subsequent drift and/or pleiotropy [87, 88]. Sperm bristles might result in costs for the donor, such as a reduced spermatogenesis rate or reduced sperm mobility in the partner’s tissue [45]. Indeed, spermatogenesis of the complex sperm with bristles of *M. lignano* takes longer than the development of the simpler sperm in *M. pusillum* (6 vs. 4 days [89–91]). However, this could also be because *M. lignano* sperm is longer, as sperm length can be associated with a longer sperm development time [92, 93]. Since several hypodermically mating species have reduced bristles, their cost in terms of movement might also be minimal, at least once they are relatively small. We also document species that very likely copulate reciprocally but do not have sperm bristles, suggesting that HI is not the only reason for bristle loss/reduction. From our observations, it appears that sperm is deposited deep inside the complex antrum of these species, so that sperm bristles may no longer be necessary to resist the suck behaviour (note, however, that this behaviour was not seen in mating observations of *M*. sp. 82 and we currently have no mating observations of *M*. sp. 68, P. Singh, pers. comm.).

The sperm of a member of the *M. pusillum* species-complex in the hypodermic clade contains electron-dense bodies [94], similar to the bristle anchor structures identified in the reciprocally mating *M. tuba* and *M. lignano* [86, 95]. If these structures are indeed remnants of sperm bristles, this would support the hypothesis (in agreement with our ASR) that bristles are symplesiomorphic in *Macrostomum*, with bristle loss as the derived condition. Moreover, sperm bristles have not been observed in three species of *Psammomacrostomum* (pers. obs.), the sister taxon of *Macrostomum* [57] (and the outgroup used in our analyses), nor in a presumably closely associated genus (i.e. *Dunwichia* [96]). Sperm bristles thus appear to be a novel trait that is restricted to the genus *Macrostomum*, but detailed investigations of sperm ultrastructure across the Macrostomorpha are needed to evaluate this hypothesis.

Even though sperm morphology and sperm design is exceptionally diverse across animals, little is known about the functional significance of this diversity [97]. Because traumatic insemination originates frequently, it offers an exciting opportunity to elucidate the relative importance of natural and sexual selection for the evolution of sperm morphology (e.g. survival during sperm storage vs. rapid and efficient movement through tissue) and contribute to an integrative view of sperm ecology [98]. To disentangle mechanisms shaping sperm length evolution, we should ideally investigate the sperm morphology of other groups of organisms that have evolved traumatic insemination and make use of natural variation in the location of sperm injection and sperm storage. For example, in bedbugs, the elaboration of the sperm receiving organ varies considerably from just being a slightly thickened epithelium to a complex spermalege [27, 99]. If movement efficiency is a crucial constraint, we might expect a negative correlation between sperm length and tissue transit time. Also of interest are comparative investigations of sperm length in species with traumatic insemination directly into the recipient’s reproductive tract (e.g. the fruit fly *Drosophila parabipectinata* [43] or the spider *Harpactea sadistica* [33]), because here movement through tissue is absent and presumably other factors related to sexual selection dominate.

In summary, our work clearly highlights that the genus *Macrostomum* is a promising taxon for the study of sperm form and function, combining a high morphological diversity with a large number of evolutionary origins, and additionally offering many desirable laboratory animal characteristics and an increasing availability of genetic tools [100–102]. *Macrostomum* will also afford more in-depth investigation of HI and shed light on this intriguing behaviour’s origin and function.

## Materials and Methods

### Phylogenetics

We performed all analyses using three recently generated phylogenies [49]. Two of these are based on 385 protein sequences (94,625 amino acid positions), include 98 species, and were inferred using maximum-likelihood (H-IQ-TREE) or Bayesian methods (H-ExaBayes). The third phylogeny (C-IQ-TREE) was also inferred using maximum-likelihood based on the same protein alignment, but additionally included partial *28S rRNA* sequences, allowing us to include 47 additional species (145 in total) [49].

### Morphological data

We used morphological data primarily from field-collected specimens, from a global sampling effort, for which we previously made available detailed image and video material [49]. We obtained both quantitative (Q) and categorical (C) data from the collected specimens or from the taxonomical descriptions of a few species we did not collect ourselves. Categorical data were determined on a per species basis, while quantitative data were taken per individual. We measured body size (Q) as the total body area and either measured or scored various aspects of the stylet (Q: length, curviness, width of the proximal opening, width of the distal opening, and asymmetry of the distal thickening; C: sharpness of the distal thickening), of the sperm (Q: total length, bristle length; C: sperm bristle state, presence of a brush, and presence of a velum), and of the antrum (Q: number of genital openings; C: antrum thickness, presence and thickness of anterior cellular valve, antrum chamber complexity, and an overall compound measure of antrum complexity). See SI Morphology for details on these measures (and Tables A1-2 and Figures A1-3 therein). Morphometric analyses were performed using ImageJ ([103], version 1.51w) and the plugin ObjectJ (version 1.04r, available at https://sils.fnwi.uva.nl/bcb/objectj/). The pixel length of structures was converted into μm using a stage micrometre. For comparative analysis we transformed body area (log10 of the square-root) and log_10_ transformed all linear measures (stylet length, width of the proximal opening, width of the distal opening, sperm length, and bristle length).

### Inferred mating syndrome

The original definition of the mating syndromes integrated morphological and behavioural traits [45], but because we lacked behavioural data for most species, we adapted these definitions, relying instead on several morphological traits and the observed received sperm location to derive the inferred mating syndrome (Table 2; see also SI Morphology). We assigned species to the hypodermic mating syndrome if we exclusively found hypodermic received sperm, since this represents strong evidence for hypodermic insemination, as opposed to species where we observed both hypodermic sperm and received sperm in the female antrum, which we classified as intermediate (Table 2). Moreover, because hypodermic sperm can be difficult to observe, especially in species with low investment into sperm production, we also assigned species that lacked received sperm observations to the hypodermic mating syndrome based on their morphology alone, namely when they had a simple antrum, a sharp stylet, and absent or reduced sperm bristles (Table 2). And while observing received sperm in the female antrum may not exclude occasional hypodermic insemination, it is a strong indication of the reciprocal mating syndrome, especially when it occurs in a species with a blunt stylet. We, therefore, assigned all species with received sperm in the antrum and a blunt stylet to the reciprocal mating syndrome (Table 2). And since some reciprocally mating species also have a sharp stylet (e.g. *M. spirale*), which could possibly wound the partner internally during mating (pers. obs.), we also assigned these species to the reciprocal mating syndrome, provided that we observed received sperm in the antrum, and that they had sperm with bristles (Table 2). These assignments based on morphology alone are supported by our analysis of correlated evolution, showing a strong association between the received sperm location and both sperm bristle state and antrum type, respectively (see Results). The inferred mating syndrome is therefore a more inclusive classification of hypodermic insemination compared to an assignment based on received sperm location alone.

### Frequent origins of hypodermic insemination

We conducted ancestral state reconstruction (ASR) of the mating syndrome and three proxies (received sperm location, sperm bristle state, and antrum state). First, we used the binary scorings (see SI Morphology) used in the tests for correlated evolution (see below). However, since we predicted that losses/reductions of some traits would transition via an intermediate state, we also performed ASR of the inferred mating syndrome, received sperm location and sperm bristle state scored as trinary states. We conducted ASR using stochastic character mapping [104] with the R package phytools [105]. We determined the appropriate transition matrix for ASR by fitting MK-models with equal rates (ER) of state transitions, with symmetric rates (SYM), with all rates different (ARD), and with a model without the possibility of gains once the trait is lost (Dollo). For traits with trinary states, we additionally fit an ordered model, where transitions are forced though an intermediate state (ORD) and an ordered model with no gains once the trait is lost, but allowing reversions from the intermediate state (ORD-Dollo). We conducted ASR for models with a corrected AIC weight >0.15 (Table 1) and used the Bayesian implementation of stochastic character mapping with a gamma prior throughout (α = 1, β = 1, i.e. a low rate of transitions) and reconstructed 1000 histories (10.000 iterations burn-in followed by 10,000 iterations and retaining every 10^th^ character history). We summarised the number of transitions as the average number of changes as well as the 95% credible interval.

### Correlated evolution

Since we do not have direct observations of received sperm in all species, we first conducted a correlation test between sperm bristle state and received sperm location, and then tested for correlated evolution between both of these variables and the antrum type. We scored all traits as binary and applied Pagel’s correlation test [106] as implemented in BayesTraits3 (available at http://www.evolution.rdg.ac.uk/BayesTraitsV3.0.2/BayesTraitsV3.0.2.html). We ran four independent MCMC chains for 510 million iterations with a burn-in of 10 million iterations and retaining every 1000^th^ iteration. Marginal likelihood was calculated using stepping-stones with 1000 power posteriors estimated with 10,000 iterations each. We assessed convergence using Gelman’s R implemented in the coda R package [107] and upon confirming convergence merged the chains for further analysis. Models were compared with Bayes factors using the marginal likelihoods (i.e. BF=2(logLH_dependent_ – logLH_independent_)). We evaluated the robustness of our results by preforming the analysis with several phylogenies and three different priors (see SI Correlated evolution).

### Convergence in morphospace

We conducted a multivariate analysis to investigate whether the convergent evolution of HI is associated with changes in a variety of reproductive traits (see SI Morphology). We summarised data on stylet, sperm and antrum morphology (including both quantitative and categorical data) using principal component analysis. Since regular principal component analysis assumes independence of observations, an assumption violated by the phylogenetic relationships of species [108], we calculated phylogenetically corrected principal components (pPCAs), using the phyPCA function in phytools with the lambda model. Since we combined data with different scales, we used the correlation matrix for all calculations. When discussing loadings of principal components, we apply an aggressive threshold of ±0.5, since although this results in erosion of power, it keeps false-positive rate within expectations [109].

### Hypodermic insemination and sperm length

To test the influence of HI on sperm length, we performed phylogenetically corrected ordinary least squared regression (PGLS) with the *gls* function in the R package *nlme* (version 3.1). We used *gls* because it allowed us to simultaneously incorporate phylogenetic signal in the residuals and account for variation in the number of measured specimens by using the sample size of the response as weights. We determined the best-fitting evolutionary model for the covariance in the residuals by comparing corrected AIC of PGLS fitted with Brownian motion, lambda or Ornstein-Uhlenbeck models. We assessed if the assumptions of the PGLS were met by checking the distributions of the phylogeny-corrected residuals for normality and profiled the likelihood of the parameter of the correlation structure (i.e. lambda or alpha). Since R-squared values are problematic for PGLS models [110] we calculated R_pred_ [111] to show model fits. As predictors, we used the binary traits included in the test of correlated evolution since they all are strong indicators of HI. Moreover, we also included the inferred mating syndrome as a predictor, but coded it as binary (hypodermic and reciprocal), and excluding the intermediate syndrome due to the low sample size of this group.

## Abbreviations

HI: Hypodermic insemination
PC: principal component
pPCA: phylogenetically corrected principal component analysis

## Acknowledgments

We thank the numerous people that have helped with field work. Especially, we are grateful for the help of, in no particular order, Werner Armonies, Benny Glasgow, Mohamed Charni, Edith Zemp, Bernhard Egger, Peter Ladurner, Gregor Schulte, Floriano Papi, Kazuya Kobayashi, Christopher Laumer, Wim Willems, Tom Artois, Christian Lott, Miriam Weber, Ana-Maria Leal-Zanchet, Kaja Wasik, Mariana Adami, Walter Salzburger, Adrian Indermaur, Bernd Egger, Fabrizia Ronco, Heinz Büscher, Victoria Huwiler, Philipp Kaufmann, Michaela Zwyer, Stefanie von Fumetti, Joe Ryan, Mark Q. Martindale, Marta Chiodin, John Evans, Leigh Simmons, Mauro Tognon, Piero Tognon, Cristiano Tognon, Pragya Singh, Nikolas Vellnow, Christian Felber, Ulf Jondelius, Sarah Atherton, Tim Janicke, Georgina Rivera-Ingraham, Ben Byrne, Yvonne Gilbert, Rod Watson, Jochen Rink, Miquel Vila-Fare, Helena Bilandžija, and Sasho Trajanovski. We thank Peter Fields and Lukas Zimmermann for IT advice. We thank Jürgen Hottinger, Daniel Lüscher and Yasmin Picton for administrative and technical support. We thank Norbert Vischer for help with writing a ObjectJ plugin. We thank Yu Zhang and his collaborators for kindly providing specimens of *M. baoanensis*.

## Data availability

All relevant data is included in the supplementary information of this publication.

## Author contribution

Jeremias N. Brand: Conceptualisation, Data Curation, Formal Analysis, Investigation, Visualisation, Writing – Original Draft Preparation, Writing – Review & Editing. Luke J. Harmon: Methodology, Supervision. Lukas Schärer: Conceptualisation, Investigation, Funding Acquisition, Project Administration, Resources, Supervision, Writing – Review & Editing

## Supp. Figures

**Figure S1.** Ancestral state reconstructions of reproductive traits using the C-IQ-TREE phylogeny. The trait and type of scoring (binary/trinary) is indicated at the bottom of each panel. Stochastic character mapping is summarised with pie charts representing the proportion of stochastic maps with the respective state. Shown is the reconstruction of the best-fitting ordered model without losses. The average number of transitions is given in Table 1, while the red stars and numbers indicate the lower-bound number of transitions that have likely occurred (i.e. separated by nodes with >95% posterior probability of the ancestral state), while acknowledging that the ancestral state of the genus is often unclear (hence the brackets).

**Figure S2.**
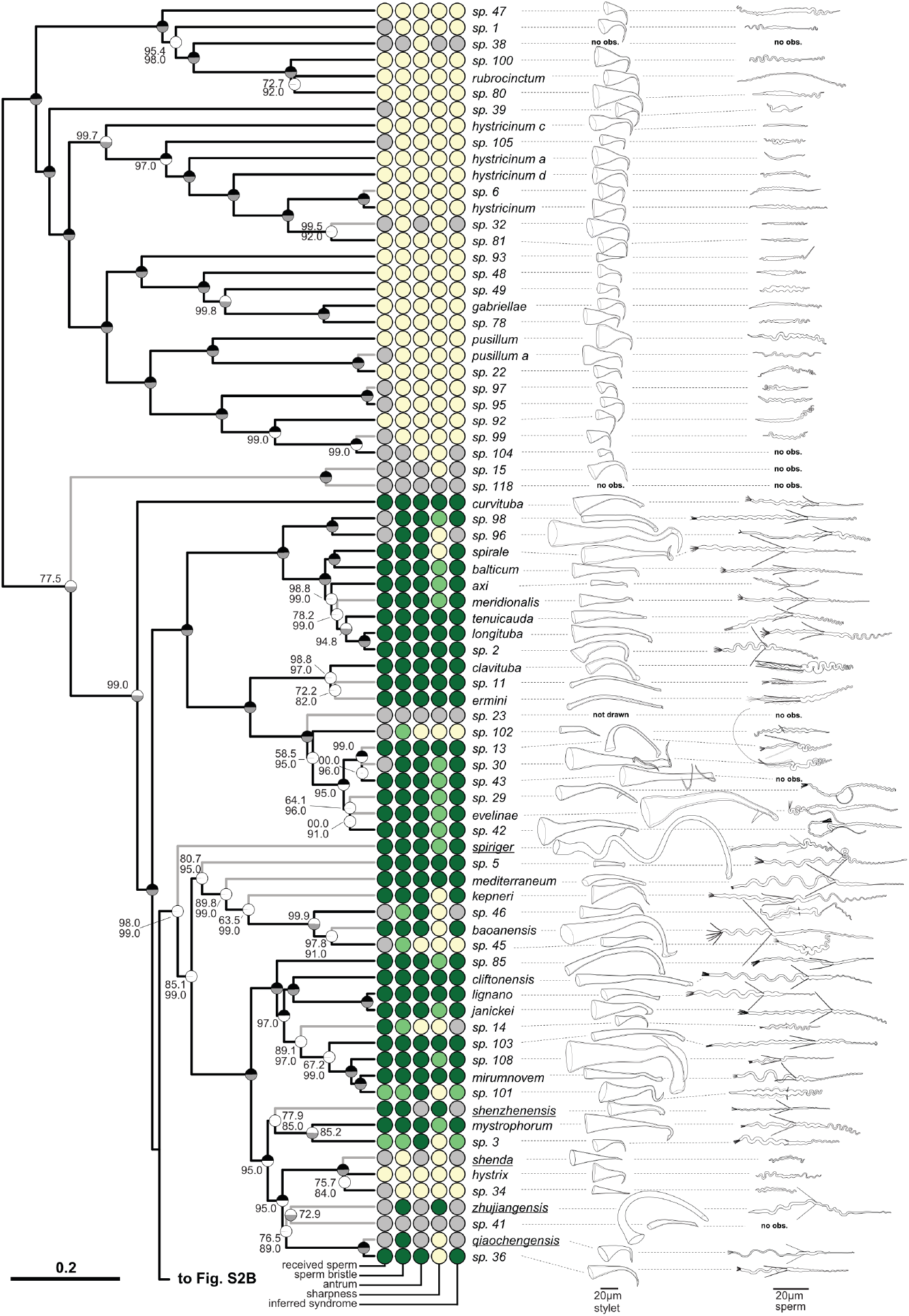
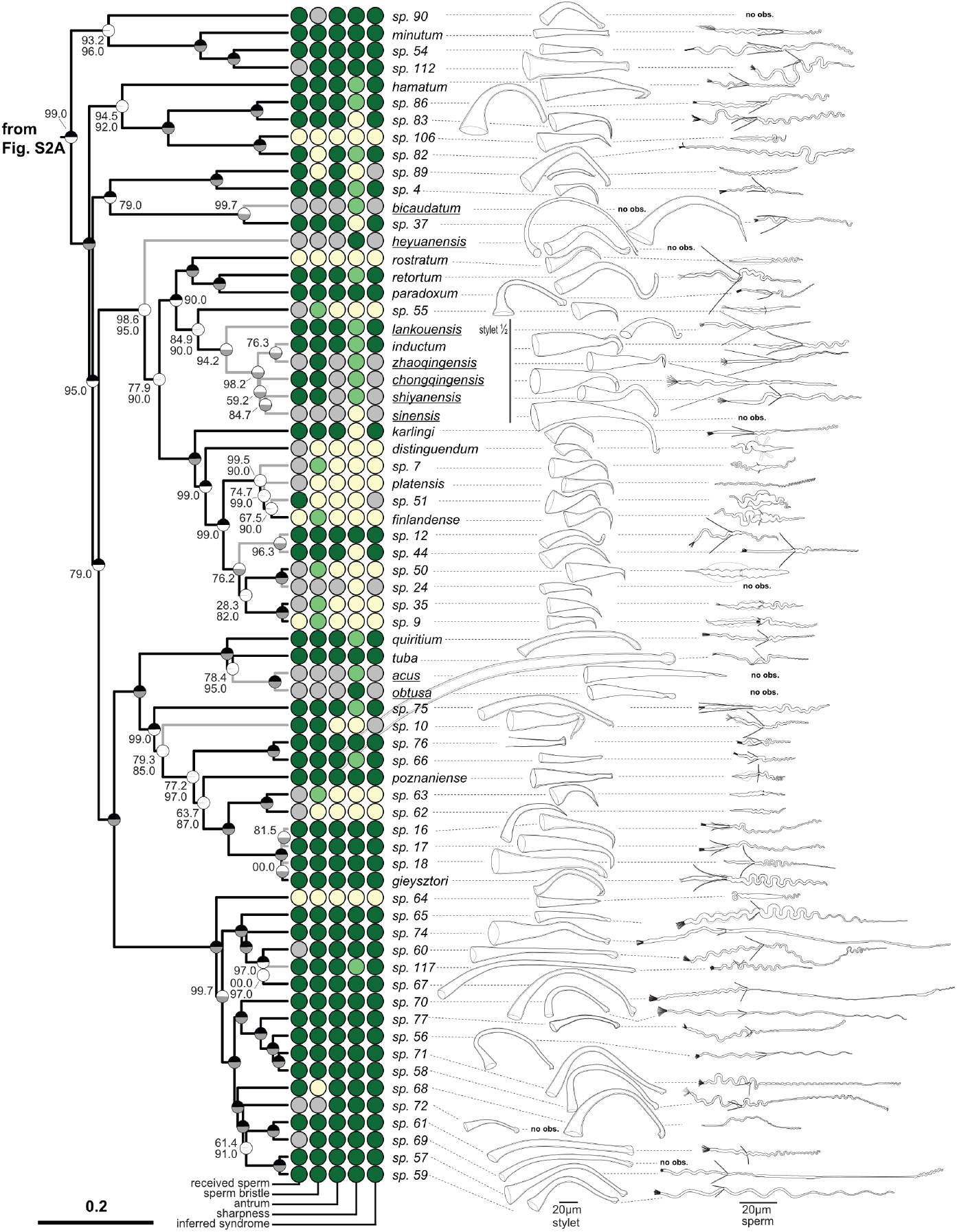
Enhanced version of Figure 2, additionally showing drawings of stylet and sperm morphology available from [49]. The ultrametric phylogeny (C-IQ-TREE) includes all 145 species from [49] (with 77 species depicted in Fig SXA and 68 species in Fig SXB). Branch supports are ultrafast bootstraps (top, black if 100) and approximate likelihood ratio tests (bottom, grey if 100). Species without available transcriptomes that were added based on a *28S rRNA* fragment are indicated with grey branches. Two phylogenetically well-separated clades the “hypodermic clade” thought to exclusively mate through hypodermic insemination (HI) and the “reciprocal clade” primarily mating reciprocally can be seen in A. Columns indicate the states of five reproductive traits from light to dark (i.e. yellow, light green and dark green for trinary states; or yellow and dark green for binary states; grey indicates missing data): received sperm location (hypodermic, both, in antrum), sperm bristle state (absent, reduced, present), antrum state (simple, thickened), sharpness of stylet (sharp, neutral, blunt), inferred mating syndrome (hypodermic, intermediate, reciprocal). Stylet and sperm morphology are drawn based on our live observations, except for species with underlined names, which were redrawn based on the species description *(M. acus, M. obtusa* and *M. sinensis* from Wang 2005; *M. heyuanensis* and *M. bicaudatum* from Sun et al. 2015; *M. chongqingensis* and *M. zhaoqingensis* from Lin et al. 2017a; *M. shiyanensis* and *M. lankouensis* from Lin et al. 2017b; *M. shenzhenensis* and *M. qiaochengensis* from Wang et al. 2017; and *M. spiriger* and *M. shenda* from Xin et al. 2019). The stylet of *M*. sp. 15 is not drawn to scale, the stylets of some species are drawn at half size (stylet ½), and the stylet of *M*. sp. 23 is not drawn since it was incomplete. Unobserved structures are marked as no observation (no obs.).

**Figure S3.** Animation of the phylomorphospace represented by PC1 and PC2 of the species in the C-IQ-TREE phylogeny. The animation initially shows a cladogram that then gradually transforms into the phylomorphospace, which was calculated using the phylomorphospace function in phytools [105].

**Figure S4.**
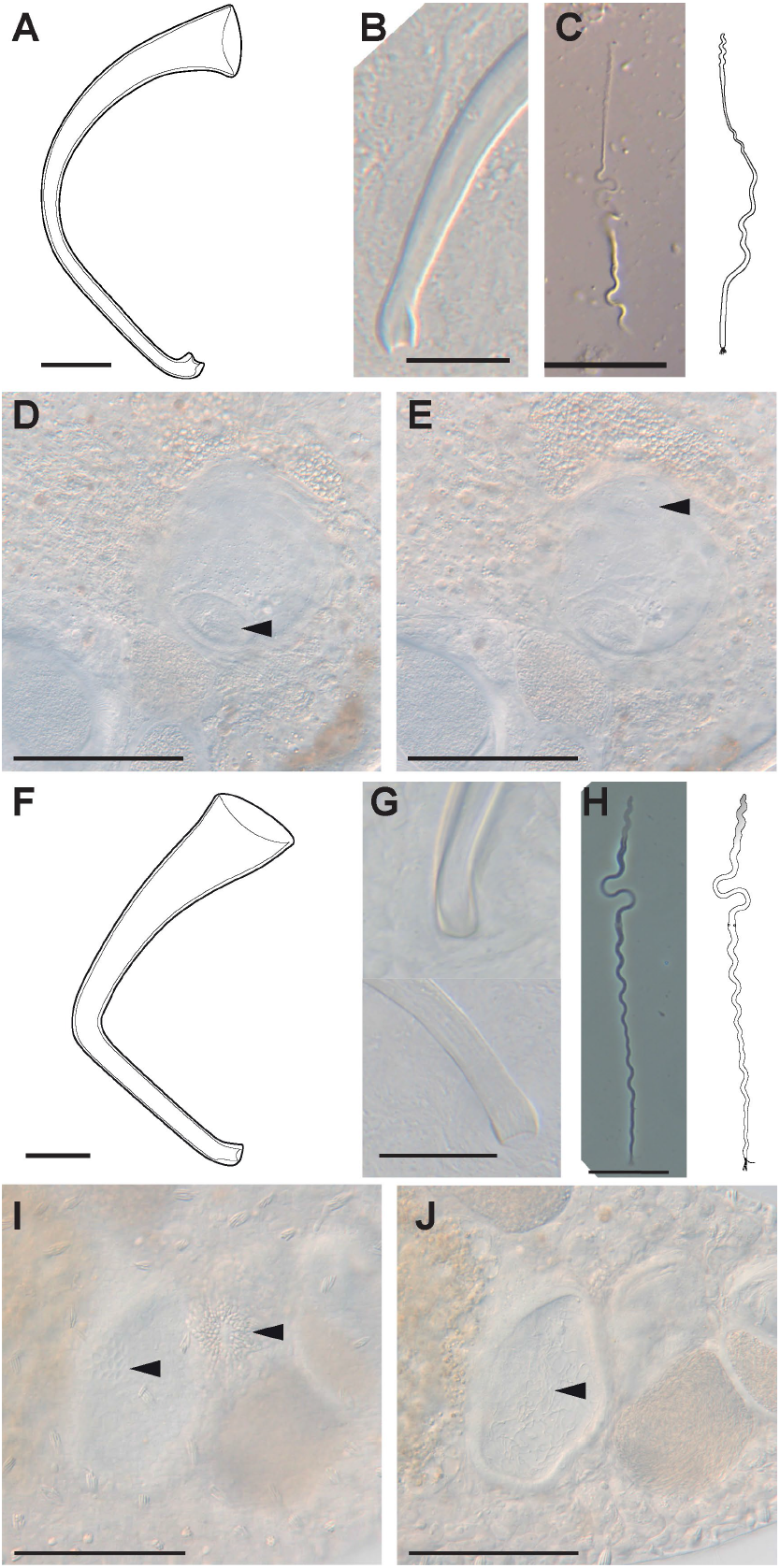
Details on the reproductive morphology of *Macrostomum* sp. 68 and *M*. sp. 82. (A-E) *M*. sp. 68 (A) Stylet drawing showing the blunt distal thickenings; (B) distal stylet tip in a smash preparation (specimen ID MTP LS 2611). (C) Sperm image (MTP LS 2686) and drawing showing what seems to be a long feeler, but no apparent sperm bristles. (D-E). Details of the antrum (MTP LS 2562) indicating the muscular connection between the female genital opening and the antrum (arrowhead in D) and the anterior second chamber containing at least one received sperm (arrowhead in E). (F-J) *M*. sp. 82 (F) Drawing of the stylet showing the slight blunt distal thickenings. (G) Distal stylet tip *in situ* (top, MTP LS 2845) and in a smash preparation (bottom, MTP LS 2846). (H) Sperm image (MTP LS 2877) and drawing indicating the modified anterior part of the sperm (shaded grey) and a less dense area approximately 1/3 along the sperm, which could be a vestigial bristle anchor location (arrowhead). (I-J) Details of the antrum (MTP LS 2848) indicating the anterior genital opening, the bursa pore (I, left arrowhead) next to the posterior genital opening, the gonopore (I, right arrowhead), both connecting into a large chamber containing many received sperm (J, arrowhead). Scale bars represent 100 μm in the antrum images and 20 μm otherwise.

**Figure S5.**
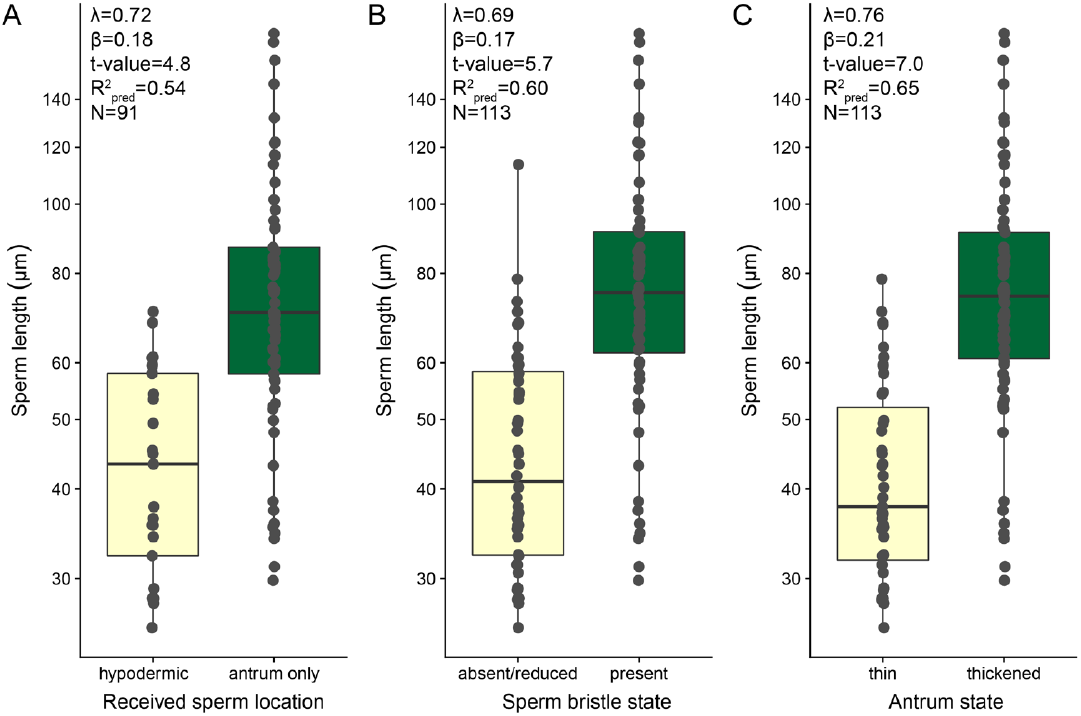
Sperm length of species dependent on (A) received sperm location, (B) sperm bristle state, and (C) antrum state. Values are slightly jittered in the x direction, and the y-axis is on a log-scale. Within each panel the main results of a PGLS analysis are given and in all tests the slopes were significant at p <0.001. Detailed results including analyses with different phylogenies (H-IQ-TREE and H-ExaBayes) are given in Table S6.

## Supp. Tables

**Table S1.** The number of specimens analysed per *Macrostomum* species for all the included quantitative traits.

see the file Tab_S1.xlsx

**Table S2.** Details on all specimens included in this study.

see the file Tab_S2.xlsx

**Table S3.** Mean species values for all morphological variables.

see file Tab_S3.xlsx

**Table S4.** Ancestral state reconstruction using stochastic character mapping.

see file Tab_S4.xlsx

**Table S5.** Scores and loadings from the phylogenetically corrected principal component analysis.

see file Tab_S5.xlsx

**Table S6.** Results of PGLS analysis of states indicating reciprocal copulation versus hypodermic insemination on sperm length. All predictors were binary, with the reference level being the state indicating hypodermic insemination.

see file Tab_S6.xlsx

**Table S7.** Results from PGLS correlating the first principal components of a phylogenetically corrected principal component analysis (pPCA) analysis including five stylet traits with the first principal component of a pPCA analysis including four antrum traits. Analysis was performed across all species and restricted to the reciprocal mating syndrome. Also given are pPCA loadings and results for all three phylogenies.

see file Tab_S7.xlsx

## References

1. Bateman AJ. Intra-sexual selection in *Drosophila*. Heredity. 1948;2:349–68.

2. Arnold SJ. Bateman’s principles and the measurement of sexual selection in plants and animals. Am Nat. 1994;144:S126–49.

3. Janicke T, Häderer IK, Lajeunesse MJ, Anthes N. Darwinian sex roles confirmed across the animal kingdom. Sci Adv. 2016;2:e1500983–e1500983.

4. Parker GA. The sexual cascade and the rise of pre-ejaculatory (Darwinian) sexual selection, sex roles, and sexual conflict. Cold Spring Harb Perspect Biol. 2014;6:a017509–a017509.

5. Puurtinen M, Ketola T, Kotiaho JS. The Good-Genes and Compatible-Genes Benefits of Mate Choice. Am Nat. 2009;174:741–52.

6. Arnqvist G, Nilsson T. The evolution of polyandry: Multiple mating and female fitness in insects. Anim Behav. 2000;60:145–64.

7. Hosken DJ, Stockley P. Benefits of Polyandry: A Life History Perspective. In: Evolutionary Biology. R.J. Macintyre and M.T. Clegg. Boston: Springer; 2003. p. 173–94.

8. Arnqvist G, Rowe L. Sexual conflict. Princeton: Princeton University Press; 2005.

9. Parker GA. Sexual conflict over mating and fertilization: An overview. Philos Trans R Soc B Biol Sci. 2006;361:235–59.

10. Rice WR. Sexually antagonistic male adaptation triggered by experimental arrest of female evolution. Nature. 1996;381:232–4.

11. Arnqvist G, Rowe L. Antagonistic coevolution between the sexes in a group of insects. Nature. 2002;415:787–9.

12. Charnov EL. Simultaneous hermaphroditism and sexual selection. Proc Natl Acad Sci U S A. 1979;76:2480–4.

13. Birkhead TR, Pizzari T. Evolution of sex: Postcopulatory sexual selection. Nat Rev Genet. 2002;3:262–73.

14. Wedell N, Hosken DJ. The evolution of male and female internal reproductive organs in insects. In: The evolution of primary sexual characters in animals. J.L. Leonard and A. Córdoba-Aguilar. Oxford: Oxford University Press; 2010. p. 307–31.

15. Morrow EH, Arnqvist G. Costly traumatic insemination and a female counter-adaptation in bed bugs. Proc R Soc B Biol Sci. 2003;270:2377–81.

16. Morrow EH, Arnqvist G, Pitnick S. Adaptation versus pleiotropy: why do males harm their mates? Behav Ecol. 2003;14:802–6.

17. Lange R, Reinhardt K, Michiels NK, Anthes N. Functions, diversity, and evolution of traumatic mating: function and evolution of traumatic mating. Biol Rev. 2013;88:585–601.

18. Schärer L, Janicke T, Ramm SA. Sexual conflict in hermaphrodites. Cold Spring Harb Perspect Biol. 2015;7:a017673.

19. Reinhardt K, Naylor R, Siva-Jothy MT. Reducing a cost of traumatic insemination: female bedbugs evolve a unique organ. Proc R Soc B Biol Sci. 2003;270:2371–5.

20. Reinhardt K, Anthes N, Lange R. Copulatory wounding and traumatic insemination. Cold Spring Harb Perspect Biol. 2015;7:a017582.

21. Benoit JB, Jajack AJ, Yoder JA. Multiple traumatic insemination events reduce the ability of bed bug females to maintain water balance. J Comp Physiol B. 2012;182:189–98.

22. Tatarnic NJ. Traumatic Insemination and Copulatory Wounding. In: Reference Module in Life Sciences. Elsevier; 2018.

23. Tatarnic NJ, Cassis G, Siva-Jothy MT. Traumatic insemination in terrestrial arthropods. Annu Rev Entomol. 2014;59:245–61.

24. Kathirithamby J, Hrabar M, Delgado JA, Collantes F, Dötterl S, Windsor D, et al. We do not select, nor are we choosy: Reproductive biology of Strepsiptera (Insecta). Biol J Linn Soc. 2015;116:221–38.

25. Peinert M, Wipfler B, Jetschke G, Kleinteich T, Gorb SN, Beutel RG, et al. Traumatic insemination and female counter-adaptation in Strepsiptera (Insecta). Sci Rep. 2016;6:25052.

26. Stutt AD, Siva-Jothy MT. Traumatic insemination and sexual conflict in the bed bug *Cimex lectularius*. Proc Natl Acad Sci. 2001;98:5683–7.

27. Siva-Jothy MT. Trauma, disease and collateral damage: conflict in cimicids. Philos Trans R Soc B Biol Sci. 2006;361:269–75.

28. Michiels NK, Newman LJ. Sex and violence in hermaphrodites. Nature. 1998;391:647–647.

29. Sluys R. Sperm resorption in triclads (Platyhelminthes, Tricladida). Invertebr Reprod Dev. 1989;15:89–95.

30. Koene JM. Tales of two snails: sexual selection and sexual conflict in *Lymnaea stagnalis* and *Helix aspersa*. Integr Comp Biol. 2006;46:419–29.

31. Koene JM, Montagne-Wajer K, Roelofs D, Ter Maat A. The fate of received sperm in the reproductive tract of a hermaphroditic snail and its implications for fertilisation. Evol Ecol. 2009;23:533–43.

32. Parker GA. Sperm competition and the evolution of ejaculates: Towards a theory base. In: Sperm competition and sexual selection. T.R. Birkhead and A.P. Møller. San Diego: Academic Press; 1998. p. 3–54.

33. Řezáč Milan. The spider *Harpactea sadistica:* Co-evolution of traumatic insemination and complex female genital morphology in spiders. Proc R Soc B Biol Sci. 2009;276:2697–701.

34. Michiels NK. Mating conflicts and sperm competition in simultaneous hermaphrodites. In: Sperm Competition and Sexual Selection. T.R. Birkhead and A.P. Møller. San Diego: Academic Press; 1998. p. 219–54.

35. Anthes N, Putz A, Michiels NK. Sex role preferences, gender conflict and sperm trading in simultaneous hermaphrodites: a new framework. Anim Behav. 2006;72:1–12.

36. Anthes N. Mate choice and reproductive conflict in simultaneous hermaphrodites. In: Animal Behaviour: Evolution and Mechanisms. P. Kappeler. Berlin, Heidelberg: Springer Berlin Heidelberg; 2010. p. 329–57.

37. Anthes N, David P, Auld JR, Hoffer JNA, Jarne P, Koene JM, et al. Bateman gradients in hermaphrodites: An extended approach to quantify sexual selection. Am Nat. 2010;176:249–63.

38. Pélissié B, Jarne P, David P. Sexual selection without sexual dimorphism: Bateman gradients in a simultaneous hermaphrodite. Evolution. 2012;66:66–81.

39. Michiels NK, Koene JM. Sexual selection favors harmful mating in hermaphrodites more than in gonochorists. Integr Comp Biol. 2006;46:473–80.

40. Jarne P, Auld JR. Animals mix it up too: The distribution of self-fertilization among hermaphroditic animals. Evolution. 2006;60:1816–24.

41. Koene JM, Schulenburg H. Shooting darts: Co-evolution and counter-adaptation in hermaphroditic snails. BMC Evol Biol. 2005;5:25.

42. Koene JM, Pförtner T, Michiels NK. Piercing the partner’s skin influences sperm uptake in the earthworm *Lumbricus terrestris*. Behav Ecol Sociobiol. 2005;59:243.

43. Kamimura Y. Twin intromittent organs of *Drosophila* for traumatic insemination. Biol Lett. 2007;3:401–4.

44. Tatarnic NJ, Cassis G. Surviving in sympatry: Paragenital divergence and sexual mimicry between a pair of traumatically inseminating plant bugs. Am Nat. 2013;182:542–51.

45. Schärer L, Littlewood DTJ, Waeschenbach A, Yoshida W, Vizoso DB. Mating behavior and the evolution of sperm design. Proc Natl Acad Sci. 2011;108:1490–5.

46. Ramm SA, Vizoso DB, Schärer L. Occurrence, costs and heritability of delayed selfing in a free-living flatworm. J Evol Biol. 2012;25:2559–68.

47. Ramm SA, Schlatter A, Poirier M, Schärer L. Hypodermic self-insemination as a reproductive assurance strategy. Proc R Soc B Biol Sci. 2015;282:20150660.

48. Winkler L, Ramm SA. Experimental evidence for reduced male allocation under selfing in a simultaneously hermaphroditic animal. Biol Lett. 2018;14:20180570.

49. Brand JN, Viktorin G, Wiberg RAW, Beisel C, Schärer L. Large-scale phylogenomics of the genus *Macrostomum* (Platyhelminthes) reveals cryptic diversity and novel sexual traits. bioRxiv. 2021;:2021/437366.

50. Schärer L, Joss G, Sandner P. Mating behaviour of the marine turbellarian *Macrostomum* sp.: these worms suck. Mar Biol. 2004;145:373–80.

51. Vizoso DB, Rieger G, Schärer L. Goings-on inside a worm: Functional hypotheses derived from sexual conflict thinking. Biol J Linn Soc. 2010;99:370–83.

52. Schärer L, Brand JN, Singh P, Zadesenets KS, Stelzer C-P, Viktorin G. A phylogenetically informed search for an alternative *Macrostomum* model species, with notes on taxonomy, mating behavior, karyology, and genome size. J Zool Syst Evol Res. 2020;58:41–65.

53. Singh P, Ballmer DN, Laubscher M, Schärer L. Successful mating and hybridisation in two closely related flatworm species despite significant differences in reproductive morphology and behaviour. Sci Rep. 2020;10:12830.

54. Patlar B, Weber M, Temizyürek T, Ramm SA. Seminal fluid-mediated manipulation of post-mating behavior in a simultaneous hermaphrodite. Curr Biol. 2020;30:143–9.

55. Weber M, Patlar B, Ramm SA. Effects of two seminal fluid transcripts on post-mating behaviour in the simultaneously hermaphroditic flatworm *Macrostomum lignano*. J Evol Biol. 2020;:jeb.13606.

56. Leasi F, Sevigny JL, Laflamme EM, Artois T, Curini-Galletti M, de Jesus Navarrete A, et al. Biodiversity estimates and ecological interpretations of meiofaunal communities are biased by the taxonomic approach. Commun Biol. 2018;1:1–12.

57. Janssen T, Vizoso DB, Schulte G, Littlewood DTJ, Waeschenbach A, Schärer L. The first multi-gene phylogeny of the Macrostomorpha sheds light on the evolution of sexual and asexual reproduction in basal Platyhelminthes. Mol Phylogenet Evol. 2015;92:82–107.

58. Egger B, Ladurner P, Nimeth K, Gschwentner R, Rieger R. The regeneration capacity of the flatworm *Macrostomum lignano—* on repeated regeneration, rejuvenation, and the minimal size needed for regeneration. Dev Genes Evol. 2006;216:565–77.

59. Maddison WP, FitzJohn RG. The unsolved challenge to phylogenetic correlation tests for categorical characters. Syst Biol. 2015;64:127–36.

60. Uyeda JC, Zenil-Ferguson R, Pennell MW. Rethinking phylogenetic comparative methods. Syst Biol. 2018;67:1091–109.

61. Anthes N, Schulenburg H, Michiels NK. Evolutionary links between reproductive morphology, ecology and mating behavior in opisthobranch gastropods. Evolution. 2008;62:900–16.

62. Beese K, Beier K, Baur B. Coevolution of male and female reproductive traits in a simultaneously hermaphroditic land snail. J Evol Biol. 2006;19:410–8.

63. Brennan PLR, Prum RO, McCracken KG, Sorenson MD, Wilson RE, Birkhead TR. Coevolution of Male and Female Genital Morphology in Waterfowl. PLOS ONE. 2007;2:e418.

64. Arnqvist G, Rowe L. Correlated evolution of male and female morphologles in water striders. Evol Int J Org Evol. 2002;56:936–47.

65. McPeek MA, Shen L, Farid H. The correlated evolution of three-dimensional reproductive structures between male and female damselflies. Evolution. 2009;63:73–83.

66. Simmons LW, Fitzpatrick JL. Female genitalia can evolve more rapidly and divergently than male genitalia. Nat Commun. 2019;10:1312.

67. Eberhard W. Female Control: Sexual Selection by Cryptic Female Choice. Princeton: Princeton University Press; 1996.

68. Arnqvist G, Edvardsson M, Friberg U, Nilsson T. Sexual conflict promotes speciation in insects. Proc Natl Acad Sci. 2000;97:10460–4.

69. Ritchie MG. Sexual selection and speciation. Annu Rev Ecol Evol Syst. 2007;38:79–102.

70. Charnov EL. Sperm competition and sex allocation in simultaneous hermaphrodites. Evol Ecol. 1996;10:457–62.

71. Schärer L. Tests of sex allocation theory in simultaneously hermaphroditic animals. Evolution. 2009;63:1377–405.

72. van Velzen E, Schärer L, Pen I. The effect of cryptic female choice on sex allocation in simultaneous hermaphrodites. Proc R Soc B Biol Sci. 2009;276:3123–31.

73. Schärer L, Pen I. Sex allocation and investment into pre- and post-copulatory traits in simultaneous hermaphrodites: The role of polyandry and local sperm competition. Philos Trans R Soc B Biol Sci. 2013;368:20120052–20120052.

74. Parker GA. Why are there so many tiny sperm? Sperm competition and the maintenance of two sexes. J Theor Biol. 1982;:281–94.

75. Parker GA. Sperm competition games: Sperm size and sperm number under adult control. Proc R Soc Lond B Biol Sci. 1993;253:245–54.

76. Parker GA. Selection on non-random fusion of gametes during the evolution of anisogamy. J Theor Biol. 1978;73:1–28.

77. Schärer L, Janicke T. Sex allocation and sexual conflict in simultaneously hermaphroditic animals. Biol Lett. 2009;5:705–8.

78. Ladurner P, Schärer L, Salvenmoser W, Rieger RM. A new model organism among the lower Bilateria and the use of digital microscopy in taxonomy of meiobenthic Platyhelminthes: *Macrostomum lignano*, n. sp. (Rhabditophora, Macrostomorpha). J Zool Syst Evol Res. 2005;43:114–26.

79. Janicke T, Marie-Orleach L, De Mulder K, Berezikov E, Ladurner P, Vizoso DB, et al. Sex allocation adjustment to mating group size in a simultaneous hermaphrodite. Evolution. 2013;67:3233–42.

80. Marie-Orleach L, Janicke T, Vizoso DB, David P, Schärer L. Quantifying episodes of sexual selection: Insights from a transparent worm with fluorescent sperm. Evolution. 2016;70:314–28.

81. Miller GT, Pitnick S. Sperm-female coevolution in *Drosophila*. Science. 2002;298:1230–3.

82. Lüpold S, Manier MK, Berben KS, Smith KJ, Daley BD, Buckley SH, et al. How multivariate ejaculate traits determine competitive fertilization success in *Drosophila melanogaster*. Curr Biol. 2012;22:1667–72.

83. Manier MK, Lüpold S, Belote JM, Starmer WT, Berben KS, Ala-Honkola O, et al. Postcopulatory sexual selection generates speciation phenotypes in *Drosophila*. Curr Biol. 2013;23:1853–62.

84. Parker GA, Immler S, Pitnick S, Birkhead TR. Sperm competition games: Sperm size (mass) and number under raffle and displacement, and the evolution of P2. J Theor Biol. 2010;264:1003–23.

85. Immler S, Pitnick S, Parker GA, Durrant KL, Lüpold S, Calhim S, et al. Resolving variation in the reproductive tradeoff between sperm size and number. Proc Natl Acad Sci. 2011;108:5325–30.

86. Willems M, Leroux F, Claeys M, Boone M, Mouton S, Artois T, et al. Ontogeny of the complex sperm in the macrostomid flatworm *Macrostomum lignano* (Macrostomorpha, Rhabditophora). J Morphol. 2009;270:162–74.

87. Jiang D, Zhang J. Fly wing evolution explained by a neutral model with mutational pleiotropy. Evolution. 2020;74:2158–67.

88. Houle D, Bolstad GH, van der Linde K, Hansen TF. Mutation predicts 40 million years of fly wing evolution. Nature. 2017;548:447–50.

89. Schärer L, Ladurner P, Seifarth C, Salvenmoser W, Zaubzer J. Tracking sperm of a donor in a recipient: an immunocytochemical approach. Anim Biol. 2007;57:121–36.

90. Giannakara A, Schärer L, Ramm SA. Sperm competition-induced plasticity in the speed of spermatogenesis. BMC Evol Biol. 2016; 16.

91. Giannakara A, Ramm SA. Self-fertilization, sex allocation and spermatogenesis kinetics in the hypodermically inseminating flatworm *Macrostomum pusillum*. J Exp Biol. 2017;220:1568–77.

92. Pitnick S, Markow TA, Spicer GS. Delayed male maturity is a cost of producing large sperm in Drosophila. Proc Natl Acad Sci. 1995;92:10614–8.

93. Pitnick S. Investment in testes and the cost of making long sperm in Drosophila. Am Nat. 1996;148:57–80.

94. Rohde K, Faubel A. Spermatogenesis of *Macrostomum pusillum* (Platyhelminthes, Macrostomida). Invertebr Reprod Dev. 1997;32:209–15.

95. Rohde K, Watson N. Ultrastructure of spermatogenesis and sperm of *Macrostomum tuba*. J Submicrosc Cytol Pathol. 1991;23:23–32.

96. Faubel A, Blome D, Cannon LRG. Sandy beach meiofauna of eastern Australia (southern Queensland and New South Wales). I. Introduction and Macrostomida (Platyhelminthes). Invertebr Syst. 1994;8:899–1007.

97. Birkhead TR, Hosken DJ, Pitnick S. Sperm Biology. 2009.

98. Reinhardt K, Dobler R, Abbott J. An ecology of sperm: Sperm diversification by natural selection. Annu Rev Ecol Evol Syst. 2015;46:435–59.

99. Hoogstraal H, Usinger RL. Monograph of Cimicidae (Hemiptera-Heteroptera). J Parasitol. 1967;53:222.

100. Wudarski J, Egger B, Ramm SA, Schärer L, Ladurner P, Zadesenets KS, et al. The free-living flatworm *Macrostomum lignano*. EvoDevo. 2020;11:5.

101. Brand JN, Wiberg RAW, Pjeta R, Bertemes P, Beisel C, Ladurner P, et al. RNA-Seq of three free-living flatworm species suggests rapid evolution of reproduction-related genes. BMC Genomics. 2020;21:462.

102. Wudarski J, Simanov D, Ustyantsev K, de Mulder K, Grelling M, Grudniewska M, et al. Efficient transgenesis and annotated genome sequence of the regenerative flatworm model *Macrostomum lignano*. Nat Commun. 2017;8:2120.

103. Rueden CT, Schindelin J, Hiner MC, DeZonia BE, Walter AE, Arena ET, et al. ImageJ2: ImageJ for the next generation of scientific image data. BMC Bioinformatics. 2017;18:529.

104. Bollback JP. SIMMAP: Stochastic character mapping of discrete traits on phylogenies. BMC Bioinformatics. 2006;7:88.

105. Revell LJ. phytools: An R package for phylogenetic comparative biology (and other things): phytools: R package. Methods Ecol Evol. 2012;3:217–23.

106. Pagel M. Detecting correlated evolution on phylogenies: a general method for the comparative analysis of discrete characters. Proc R Soc Lond B. 1994;255:37–45.

107. Plummer M, Best N, Cowles K, Vines K. CODA: Convergence diagnosis and output analysis for MCMC. R News. 2006;6:7–11.

108. Revell LJ. Size-correction and principal components for interspecific comparative studies. Evolution. 2009;63:3258–68.

109. Peres-Neto PR, Jackson DA, Somers KM. Giving meaningful interpretation to ordination axes: assessing loading significance in principal component analysis. Ecology. 2003;84:2347–63.

110. Ives AR. R^2^s for Correlated Data: Phylogenetic Models, LMMs, and GLMMs. Syst Biol. 2019;68:234–51.

111. Ives A, Li D. rr2: An R package to calculate R^2^s for regression models. J Open Source Softw. 2018;3:1028.

112. Wang A-T. Three new species of the genus *Macrostomum* from China (Platyhelminthes, Macrostomida, Macrostomidae). Acta Zootaxonomica Sin. 2005;30:714–20.

113. Sun T, Zhang L, Wang A-T, Zhang Y. Three new species of freshwater *Macrostomum* (Platyhelminthes, Macrostomida) from southern China. Zootaxa. 2015;4012:120–34.

114. Lin Y, Zhou W, Xiao P, Zheng Y, Lu J, Li J, et al. Two new species of freshwater *Macrostomum* (Rhabditophora: Macrostomorpha) found in China. Zootaxa. 2017;4329:267.

115. Lin Y-T, Feng W-T, Xin F, Zhang L, Zhang Y, Wang A-T. Two new species and the molecular phylogeny of eight species of *Macrostomum* (Platyhelminthes: Macrostomorpha) from southern China. Zootaxa. 2017;4337:423.

116. Wang L, Xin F, Fang C-Y, Zhang Y, Wang A-T. Two new brackish-water species of *Macrostomum* (Platyhelminthes, Macrostomida) from mangrove wetland in southern China. Zootaxa. 2017;4276:107.

117. Xin F, Zhang S-Y, Shi Y-S, Wang L, Zhang Y, Wang A-T. *Macrostomum shenda* and *M. spiriger*, two new brackish-water species of *Macrostomum* (Platyhelminthes: Macrostomorpha) from China. Zootaxa. 2019;4603:105.

