## Supporting information for "Frequent origins of traumatic insemination involve convergent shifts in sperm and genital morphology"

### SI Morphology

#### Stylet

We measured stylet length by placing two segmented lines along its sides (Figure A1) and then taking the average of both. We also used this line to measure the stylet's curviness, by summing all the angles between the neighbouring line segments for each line and again taking the average. Due to this, longer stylets may tend to be curvier simply because they have more line segments and thus more summed angles. However, we decided against standardising by length since this would lead to autocorrelation in our principal component analysis (and since such standardisation is easy to implement for future use). We also measured the width of the proximal and distal stylet openings (Figure A1). As a measure of distal asymmetry, we then took the absolute difference between the size of the two distal thickenings. Finally, we categorised the stylet according to how sharp its distal end is ("stylet sharpness") on a per species basis according to the criteria in Table A1.

Measurements of stylets were performed using *in situ* images and videos of live worms (590 stylets) as well as images of smash preparations (462 stylets). We assessed if the preparation method introduced a measurement bias by comparing stylet length in specimens for which both *in situ* and smash preparations were available (103 specimens across 59 species). We found that *in situ* estimates were slightly shorter (on average by 3.1 µm; median by 3.1%; paired t-test: $t=5.8$, $df=102$, $p< 0.001$). Even though we introduced a slight bias by measuring under both setups, this allowed us to greatly increase the sample size, since stylets can be more challenging to measure *in situ* in some orientations. In cases where both estimates were available, we used the mean between the two. In some specimens, it was not possible to measure the stylet's length, but we still measured the width of the proximal and distal openings (103 cases). Conversely, a few times we were able to measure stylet length but not the width of the distal opening (11 cases), the width of the proximal opening (one case) or both (two cases). Finally, we could not spatially calibrate the images of 7 specimens, but still collected curviness information on these stylets.

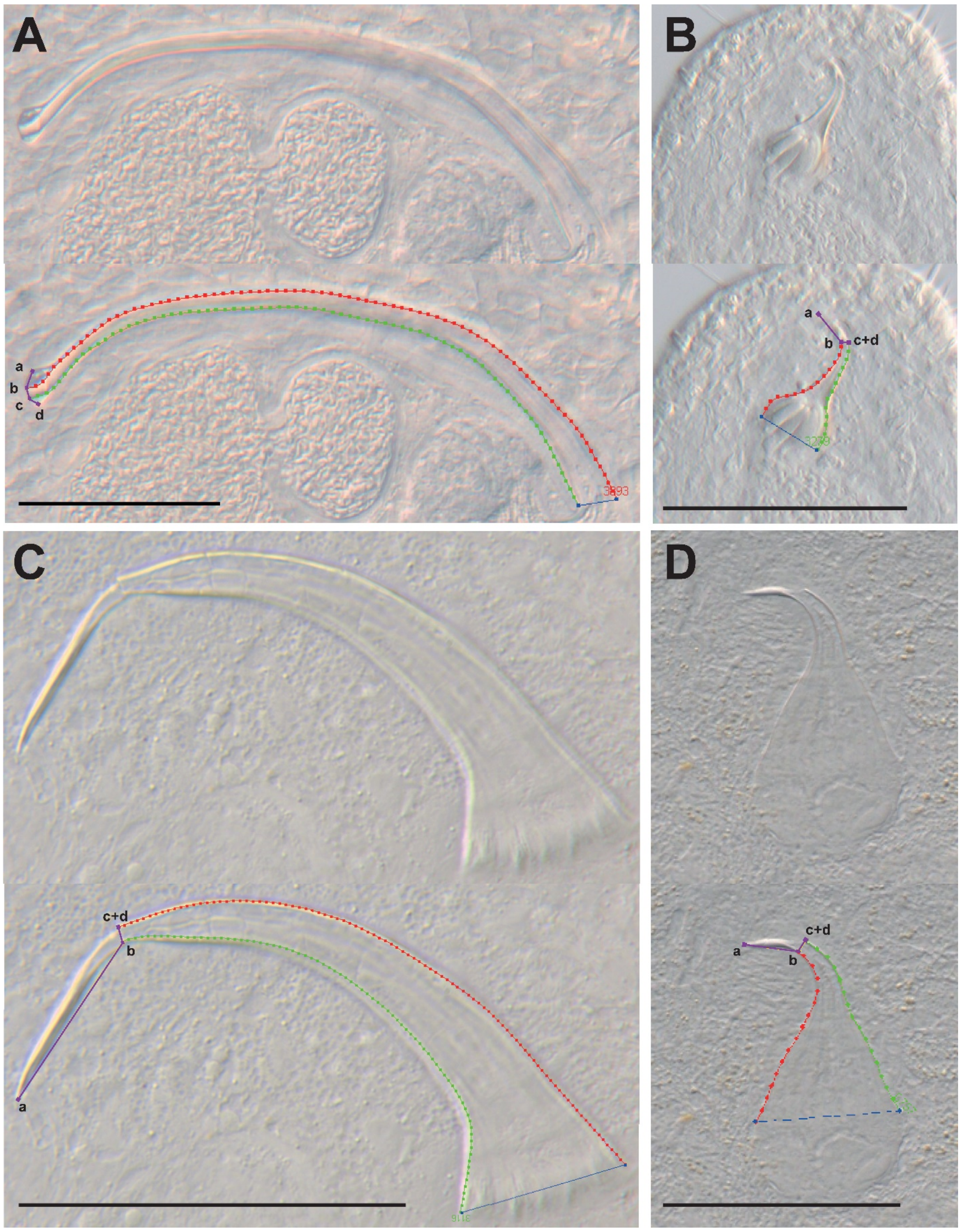

**Figure A1 Details on stylet morphometrics, measured either from images taken *in situ* in live worms (A, B) or from smash preparations (C, D).** In each panel, the top shows the raw image and the bottom the measured lines. The segmented red and green stylet length lines are used to determine stylet length and curviness, and the blue line measures the width of the proximal opening. Each purple segmented line consists of four points (a, b, c, and d) with a to b and c to d measuring the size of the 'red' and 'green' distal thickenings, respectively, and with b to c measuring the width of the distal opening (note that points a and d can be the same as points b and c, respectively, when a side lacks a distal thickening). To determine points b and c, the stylet length line that terminates first (green in panel D) is selected and a line orthogonal to its last line segment drawn through its endpoint (here considered point c). Where this orthogonal line intersects the second stylet length line (red in panel D) is the termination point of the second line and designated point b. In cases where the orthogonal line did not intersect with the other side of the stylet, we instead simply defined the end of the second line as point b. The size of the distal thickenings was then measured as the distance from the endpoints of the orthogonal line (e.g. a relatively symmetrical distal thickening in A and asymmetric thickenings in B-D), which is akin to increasing the line width of the length lines (red and green lines) and placing the point on the structure that is obscured last. (A) *in situ* stylet of *M.* sp. 67 (specimen ID MTP LS 2613). (B) *in situ* stylet of *M.* sp. 48 (MTP LS 2128). (C) smash preparation stylet of *M.* sp. 37 (MTP LS 2082). (D) smash preparation stylet of *M. hystricinum* (MTP LS 2133). Scale bars are 50 µm in A, C and D, and 25 µm in B.

#### Sperm

We measured sperm length of 1765 sperm across 504 specimens by placing a segmented line along the midline, starting from the most distal part of the feeler and terminating just anterior of the brush (if present, Figure A2). Note that this measure of total sperm length is different from that used in a previous study on *M. lignano* where the sperm feeler was excluded [1] but equivalent to the measure used in [2]. This was necessary since, compared to *M. lignano*, the boundary between the feeler and the body of the sperm is less evident in many species. The feeler was excluded in [1] since they determined that repeatability was lower when it was included. However, its inclusion in the current study potentially captures some interesting variation.

We measured sperm bristle length by placing a line along its axis. If possible, we measured both bristles of a sperm, averaged them per sperm, then averaged them per specimen before finally taking the average across all specimens measured per species (1021 bristles measured across 294 specimens). We also categorised sperm on a per species basis according to their bristle type ("sperm bristle state") and whether they carried a velum ("sperm velum") or a brush ("sperm brush") (Table A1). Among the species in which the sperm bristles are visible in the light microscope, these have a fairly continuous distribution. However, very small bristles likely do not serve a function, or at least are too small to anchor sperm in the antrum during the suck behaviour. In some species, the sperm bristles are so small that they do not even protrude outside of the sperm (e.g. *M.* sp. 50). We consider these bristles as reduced and categorised them as such to highlight our functional hypothesis. Since the trait is continuous, an exact cut-off point is hard to define, and we decided that *M. poznaniense* is the species with the smallest sperm bristles (3.2 μm) that we do not consider reduced (Figure A3).

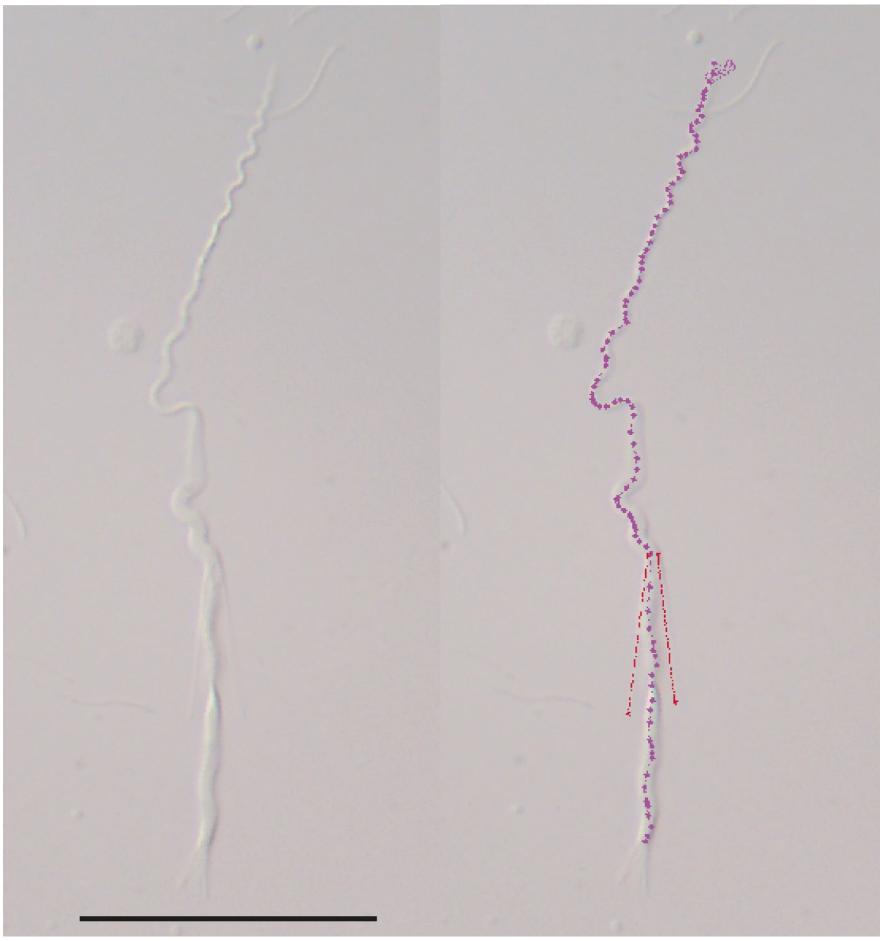

**Figure A2 Details on sperm morphometrics, measured from images taken from smash preparations.** Sperm image of *M. karlingi* (specimen ID MTP LS 3357) on the left, with an example of the segmented sperm length line (purple) and the lines used to measure sperm bristle length (red). Scale bar is 25 µm.

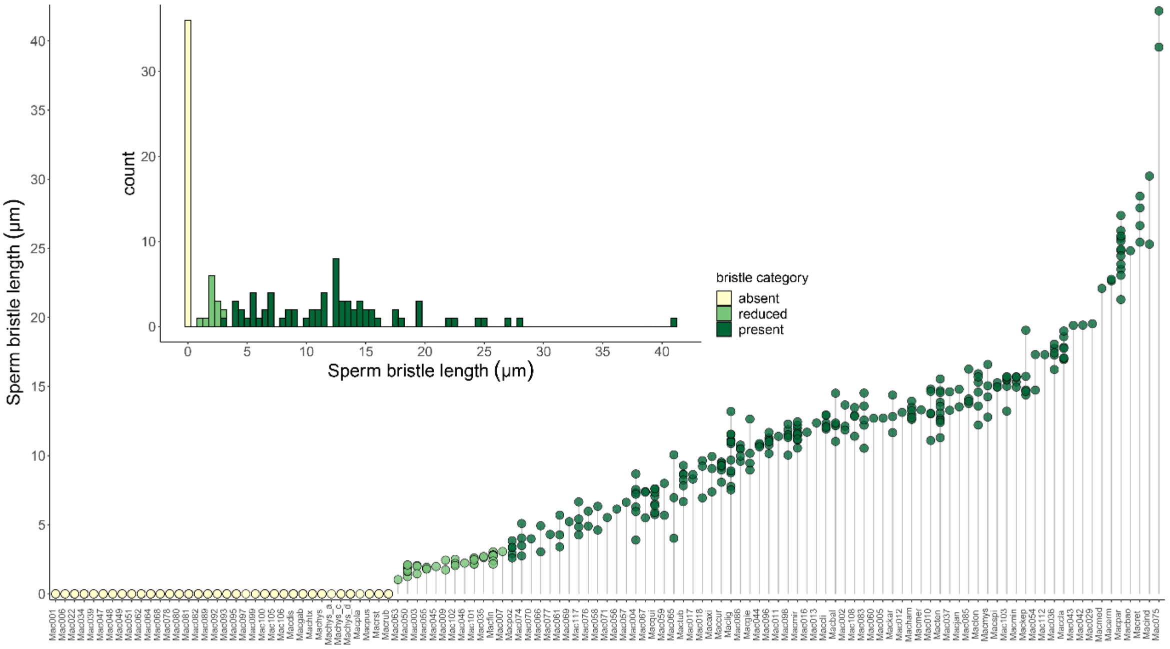

**Figure A3 Distribution of sperm bristle length across 116 *Macrostomum* species.** The large plot shows the individual data points of the species ordered by mean bristle length and the inset is a histogram of mean bristle length. The colours indicate the assigned sperm bristle state.

#### Female antrum

Female antrum morphology was categorised by examining all available material from a species and scoring the species as a whole. We examined the thickness of the overall antrum wall (“antrum thickness”), the presence of a visible anterior thickening of the female antrum (“cellular valve”) [3], the complexity of the chambers in the antrum (“antrum chamber”), and we counted the number of female genital openings (Table A2). For the ancestral state reconstructions, the analysis of correlated evolution, and the PGLS on sperm length, we scored a combined measure of antrum thickness and cellular valve as binary (“antrum state”). Finally, we summed the antrum thickness, cellular valve, and antrum chamber scores to obtain a summary score (“antrum complexity”) (Table A2). When the antrum wall is thickened throughout, it is more difficult to see if there is a cellular valve and potentially there is some conflict between the antrum thickness and cellular valve scorings. However, this would occur only in specimens that already achieve a high score on antrum complexity and might just introduce a slight downward bias for high values.

#### Received sperm location

Whenever possible, we examined specimens for received sperm in the antrum and within the tissue across the entire body. Based on our observations, we then summarised our findings for each species as categories with a binary and a trinary coding. In the binary coding, we grouped all species where we observed sperm within the tissue, irrespective of whether we also observed sperm in the antrum. In the trinary coding, we added a category especially for species with received sperm in both the antrum and the tissue (Table A1).

**Table A1. Categorisation of male reproductive morphology and the received sperm location.**
Given are the state definitions and how they were translated into ordinal coding.

| State | Binary coding | Trinary coding |
|---|---|---|
| ***Stylet sharpness*** | | |
| Rounded/blunt distal thickening | | 0 |
| No clear blunt or sharp distal thickening | | 1 |
| Sharp distal thickening (terminating in edge or point) | | 2 |
| ***Sperm bristle state*** | | |
| Absent | 0 | 0 |
| Reduced (<3.2 um) | 0 | 1 |
| Present (>3.2 um) | 1 | 2 |
| ***Sperm brush*** | | |
| Absent | 0 | |
| Present | 1 | |
| ***Sperm velum*** | | |
| Absent | 0 | |
| Present | 1 | |
| ***Received sperm location*** | | |
| Hypodermic only | 0 | 0 |
| Hypodermic and antrum | 0 | 1 |
| Antrum only | 1 | 2 |

**Table A2. Categorisation female reproductive morphology.**
Given are the state definitions and how they were translated into ordinal coding.

| State | Coding |
|---|---|
| ***Antrum thickness*** | |
| Antrum wall very thin, only visible when egg is present | 0 |
| Thin wall, but structure also visible without egg present | 1 |
| Clear thickening | 2 |
| Strong thickening | 3 |
| ***Cellular valve*** | |
| No clear anterior thickening | 0 |
| Clear anterior thickening | 1 |
| Thickening extends far anterior | 2 |
| ***Antrum chamber*** | |
| One simple chamber | 0 |
| Elaborated chamber and/or vagina | 1 |
| Multiple chambers | 2 |
| ***Genital openings*** | |
| One opening | 1 |
| Two openings | 2 |
| ***Antrum state*** | |
| Antrum thickness <2 AND cellular valve = 0 | 0 |
| Antrum thickness >1 OR cellular valve > 0 | 1 |
| ***Antrum complexity*** | |
| ∑ (antrum thickness, cellular valve, antrum chamber) | |

#### SI Female openings

A second female opening appears to have originated multiple times across the genus *Macrostomum*. It is present in *M. spiriger* Wang & Xin 2019 [4], in *M. gieysztori* Ferguson, 1939 [5] (and its three close relatives, *M.* sp. 16, *M.* sp. 17 and *M.* sp. 18), in *M. paradoxum* (An-der-Lan, 1939) [6] (previously in the genus *Promacrostomum*; see [7]), in *M.* sp. 82, and one additional species, *M. palum* (Sluys, 1986) [8] (previously also *Promacrostomum*; see [7]), that currently lacks phylogenetic placement. Each of these species has one opening that is, and one that is not, associated with cement or shell glands, and we refer to these as the gonopore and bursa pore, respectively. The gonopore is assumed to be homologous to the single female genital opening in all other *Macrostomum,* since it is also surrounded by cement glands [8]. The presumably novel bursa pore is always associated with a small chamber (bursa) that has strong circular musculature [6]. Interestingly, the bursa pore is located anterior to the gonopore in *M. spiriger* [4], in *M. paradoxum* (cement glands not drawn by [6], but confirmed by pers. obs., e.g. MTP LS 3476) and *M.* sp. 82 (pers. obs., e.g. MTP LS 2848), while it is posterior to it in *M. palum* [8] and *M. gieysztori* ([9]; and in its three relatives; pers. obs., e.g. MTP LS 845), further supporting their independent origin. Moreover, in *M. paradoxum* the bursa is described as being connected to the gut via a genito-intestinal duct [6], which could potentially allow sperm digestion. However, we were not able to locate this duct in our *in vivo* observations and may have to resort to histology to confirm its existence.

#### SI Pathways to hypodermic insemination

While 117 of the 145 studied species could be assigned to either the hypodermic or reciprocal mating syndrome, some species were not easily identified as performing either type of mating behaviour (Table 2), which may either have been due to a lack of detailed observations, or more interestingly, due to potentially transitional patterns that deviated from these syndromes. This included two species that we categorised as intermediate because we observed sperm both in the antrum and embedded inside the recipient's tissues (light green triangles in Figure 4). Both species (*M.* sp. 3 and *M.* sp. 101) have a sharp stylet and sperm with reduced bristles, fitting with the hypodermic mating syndrome (Fig. S2). However, we have scored both species as having a thickened antrum because *M.* sp. 3 has a visible antrum wall and a clear cellular valve, while *M.* sp. 101 has a strongly thickened antrum wall with the cellular valve extending far anterior

(Table S3). These antrum states indicate the reciprocal mating syndrome, and both species indeed have intermediate values of PC1. Multiple observations of hypodermic sperm in *M.* sp. 3 show them to be embedded deeply in the anterior wall of the antrum and also more deeply in the tissue lateral to the body axis, extending up to the ovaries, as well as in the tail plate (Figure A4). And while in *M.* sp. 101 we did not observe sperm as deeply in the recipient's tissues, some were fully embedded in the cellular valve and just anterior to it, and thus close to the developing eggs (Figure A5). One explanation for these findings would be that during mating, the stylet of both species pierces the recipient's antrum wall and sperm is traumatically injected into the body internally. Unfortunately, no copulations were observed in mating observations of *M.* sp. 101, but this species has been seen performing the suck behaviour, as expected if ejaculate is, at least partially, deposited in the antrum (P. Singh, pers. comm.). We currently lack mating observations for *M*. sp. 3 and further investigations of the mating behaviour and the antrum histology of both species would be highly desirable.

The observations in *M*. sp. 3 and *M*. sp. 101 suggest a possible route to hypodermic insemination via the initial evolution of a traumatic stylet in reciprocally mating species. A sharp stylet could provide anchorage during copulation, as potentially occurs in *M. spirale* (pers. obs.) and *M. hamatum* (P. Singh, pers. comm.), but it may also serve to stimulate the partner during copulation, or aid in the destruction or removal of rival sperm already present in the partner's antrum. Finally, internal wounding by the stylet may help to embed sperm in the antrum wall and/or cellular valve to prevent their removal, either by rival mating partners or by the recipient during the suck behaviour. However, the fact that we observe many *Macrostomum* species with stylets with blunt distal thickenings suggests that such internal wounding may not always be advantageous for the donor and that these structures may have evolved to avoid harm to the recipient [10]. Irrespective of the initial selective advantage that internal wounding may confer, it could then evolve further to complete internal traumatic insemination, and eventually complete avoidance of the female genitalia and hypodermic insemination via the epidermis.

Accidental sperm transfer due to copulatory wounding has generally been suggested as a possible route from copulation to traumatic insemination (e.g. in traumatically mating bedbugs) [11, 12]. In bedbugs, the attachment during mating is a two-step process, indicating that traumatic insemination was evolutionarily preceded by traumatic penetration for attachment [11]. Similar transitions have also been proposed for *Drosophila* species in the melanogaster group, where extragenital wounding structures are typically used for anchorage during copulation. In traumatically inseminating *Drosophila,* these structures are modified and pierce the mating partner's integument to inject sperm into the genital tract [13, 14]. In this light our findings are remarkable, because previous examples only compared species with and without traumatic insemination, whereas we potentially observe species "in the act" of transitioning to traumatic insemination. These intermediate species should be excellent targets for future studies of the costs and benefits of traumatic insemination, especially because the two candidates represent independent evolutionary transitions.

Three additional species (*M*. sp. 14, *M*. sp. 51 and *M*. sp. 89) were also difficult to classify because, although their morphology indicates hypodermic insemination, we clearly observed received sperm in the antrum. *M*. sp. 51 and *M*. sp. 89 grouped with the hypodermically mating species in PC1 (red arrowheads in Figure 4), while we did not include *M*. sp. 14 in this analysis due to missing data for sperm bristle length. We found sperm within the antrum in only 1 of 4 specimens in *M*. sp. 51 and 1 of 12 specimens in *M*. sp. 89, and it is thus possible that sperm is hypodermically injected and later enters the antrum when an egg passes through the cellular valve into the antrum before egg laying. But these species could also represent an intermediate state between the mating syndromes. We observed sperm in the antrum in 3 of 5 specimens in *M*. sp. 14 and it therefore seems less likely that sperm entering during the transition of the egg into the antrum is the cause of its presence here as well. Instead, sperm is probably deposited in the antrum by the mating partner during copulation. However, based on its general morphology, we predict that closer investigations of this species will reveal hypodermic received sperm in a similar location as found in the other intermediate species (*M*. sp. 3 and *M*. sp. 101).

Finally, we were not able to assign *M*. sp. 10 to a mating syndrome, because—although we found received sperm in the antrum and its sperm carry long bristles—it also has a sharp stylet and a simple antrum. From our previous findings, we would expect this species to have a thickened antrum due to its interaction with sperm and the mating partner's stylet. This discrepancy could possibly be attributed to misclassification of the antrum morphology, since *M*. sp. 10 has very pronounced shell glands, making it difficult to see the anterior part of the antrum, possibly obscuring a thickening or cellular valve (see specimen IDs MTP LS 788 and MTP LS 801 for a possibly thin cellular valve).

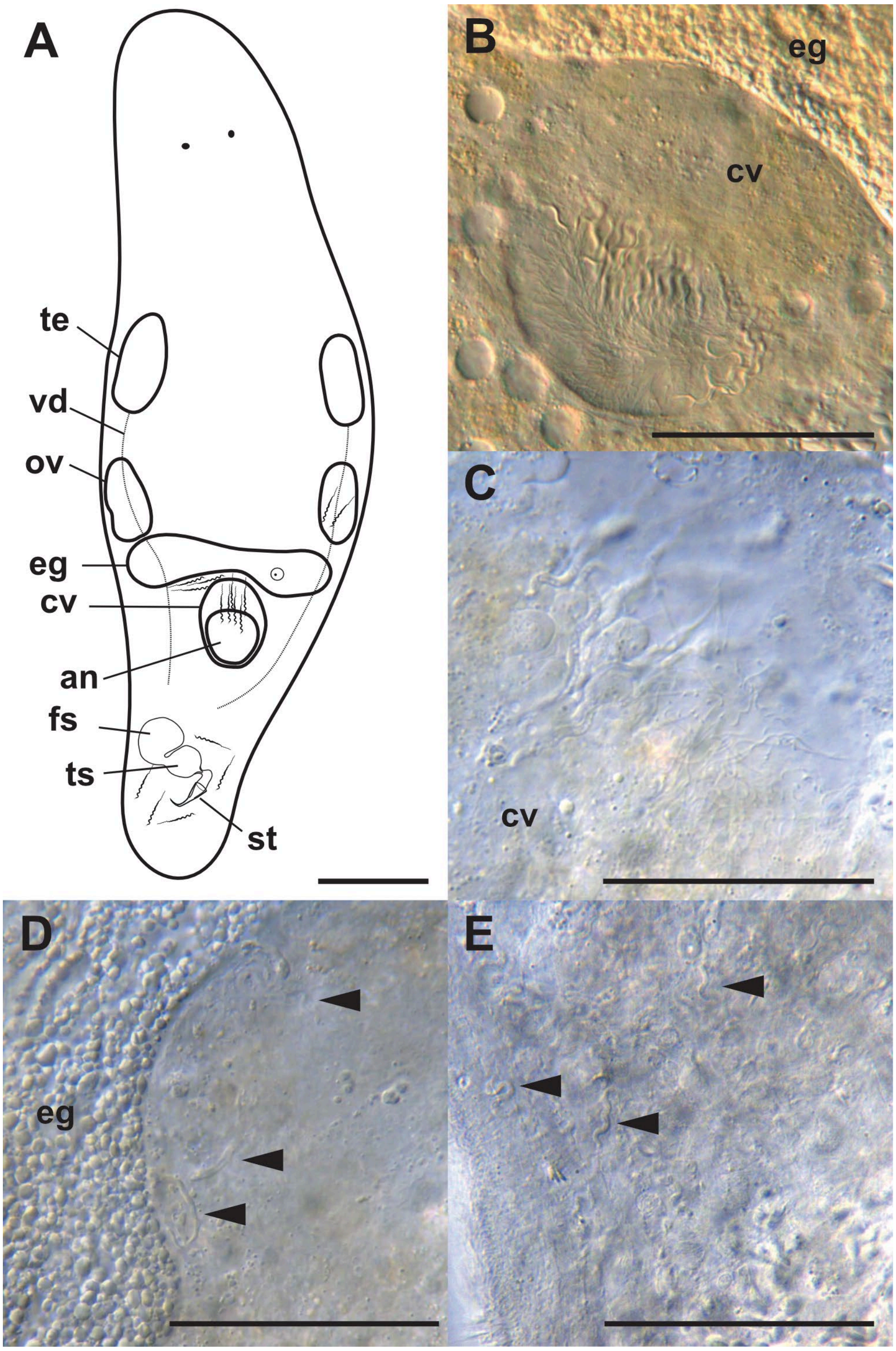

**Figure A4 Detailed observations of received sperm location in *Macrostomum* sp. 3.** (A) Drawing of the sexual organs (te: testis, vd: vas deferens, ov: ovary, eg: developing egg, cv: cellular valve, an: antrum, fs: false seminal vesicle, ts: true seminal vesicle, st: stylet), with sperm drawn at locations where they were observed. (B-E) show sperm *in situ*. (B) sperm in the antrum (specimen ID MTP LS 3286) embedded in the cellular valve (anterior); (C) sperm in the cellular valve with loose tissue (MTP LS 3314); (D) sperm close to developing egg (MTP LS 3314); (E) sperm close to the ovary (MTP LS 3317). Scale bars represent 100µm in (A) and 50 µm otherwise.

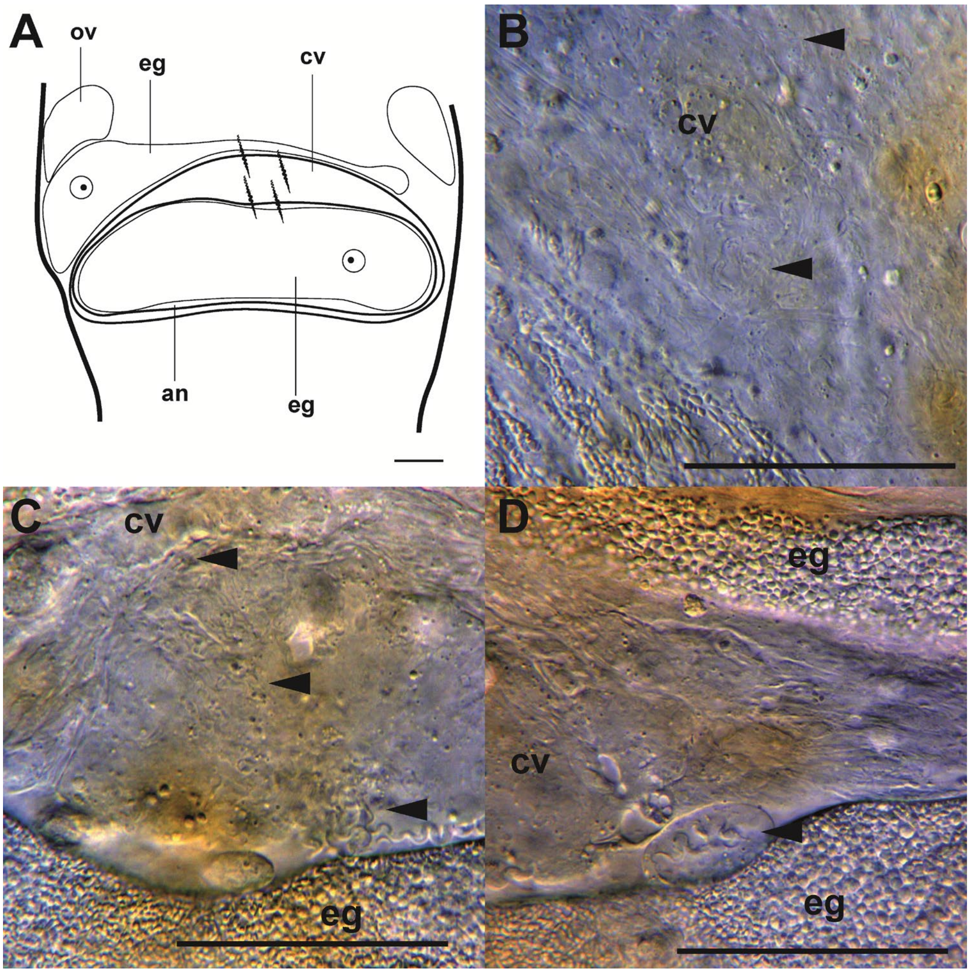

**Figure A5. Detailed observations of received sperm location in *Macrostomum* sp. 101.** (A) Drawing of antrum region (ov: ovary, eg: developing egg, cv: cellular valve, an: antrum), with sperm drawn at locations where they were observed. (B-D) show sperm *in situ* with anterior on the top. (B) sperm (arrow) embedded in cellular valve (specimen ID MTP LS 3206); (C) sperm (arrow) in cellular valve and antrum (MTP LS 3127); (D) same specimen as C with dislodged sperm (arrow) surrounded by what appears to be cellular valve derived tissue. Scale bars represent 100µm in (A) and 50µm otherwise.

### SI Correlated evolution

We performed tests for correlated evolution on three trait combinations with three different phylogenetic trees and three different priors for each test. To evaluate sensitivity to the prior, we ran all analyses with a uniform prior (U 0 100), an exponential prior (exp 10) and a reversible-jump hyperprior with a gamma distribution between 0 and 1 for both the rate prior and the hyperprior (rjhp gamma 0 1 0 1). To assess the influence of the phylogeny we conducted these tests on three different phylogenies (C-IQ-TREE, H-IQ-TREE, and H-ExaBayes). This setup resulted in 54 model runs consisting of 216 MCMC chains (three trait pairs x three priors x three topologies x four chains for the independent and the dependent model). The number of species included was largest for the correlation between sperm bristle state and antrum type (C-IQ-TREE: 124; H-IQ-TREE & H-ExaBayes: 95), and similar for received sperm location and antrum type (C-IQ-TREE: 100; H-IQ-TREE & H-ExaBayes: 77) and received sperm location and sperm bristle state (C-IQ-TREE: 101; H-IQ-TREE & H-ExaBayes: 76).

Here we first show the mean transition rates inferred (Figures A6-A8) and then give the posterior distributions for all transition rates and the likelihood for each combination of parameters (Figures A9-A17). The analyses were moderately sensitive to the chosen priors, with the largest Bayes factors for the uniform prior, slightly lower values for the exponential prior, and substantially lower values for the reversible-jump hyperprior in all tests. The reversible-jump hyperprior in some cases resulted in posterior distributions of the likelihood with multiple peaks, especially in runs using only the species with a transcriptome (H-IQ-TREE and H-ExaBayes). These models also showed two peaks in the distributions in the rates of transitions that were very rare (e.g. the transition away from hypodermic received sperm, which likely did not occur in these datasets) and were thus difficult to estimate and more sensitive to the prior. This phenomenon can also be seen in these rates' posteriors using the uniform and the exponential priors since the posteriors are very similar to the priors. Reversible-jump hyperpriors resulted in more well-behaved posteriors when running on the C-IQ-TREE phylogeny, likely because more data was available to influence the posteriors and because of the presence of species with thickened antrum state in the finlandense clade, which allows for all rates to be estimated more easily. In the main text, we present the results from C-IQ-TREE runs with the exponential prior, because these runs had the highest marginal likelihoods of the dependent model in all analyses.

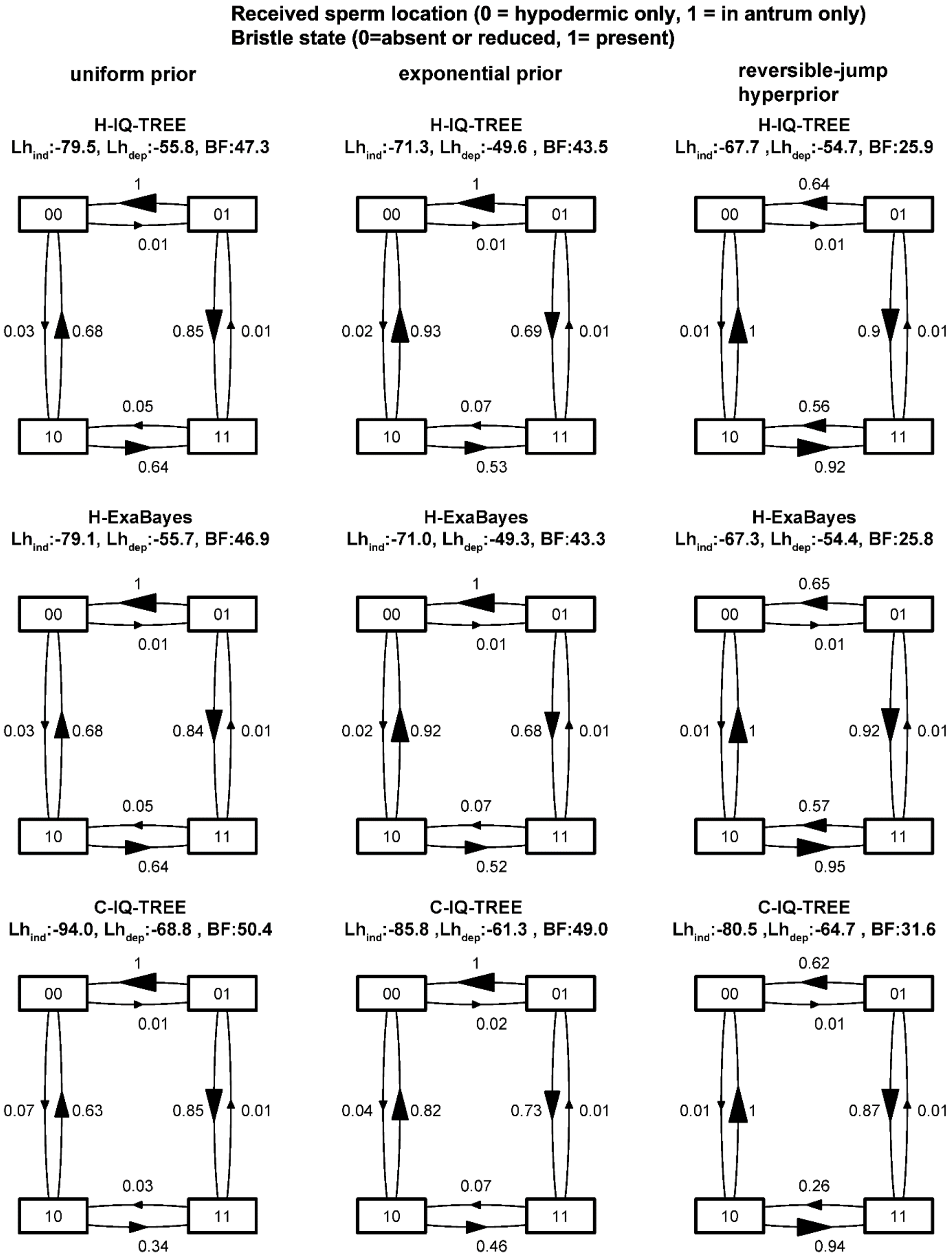

**Figure A6. Visualisation of the transition matrices for received sperm location and sperm bristle state.** Transition rates are scaled so that the largest is unity. Rows show the different phylogenies and columns show different priors (and the bottom centre is what is shown in Figure 4A).

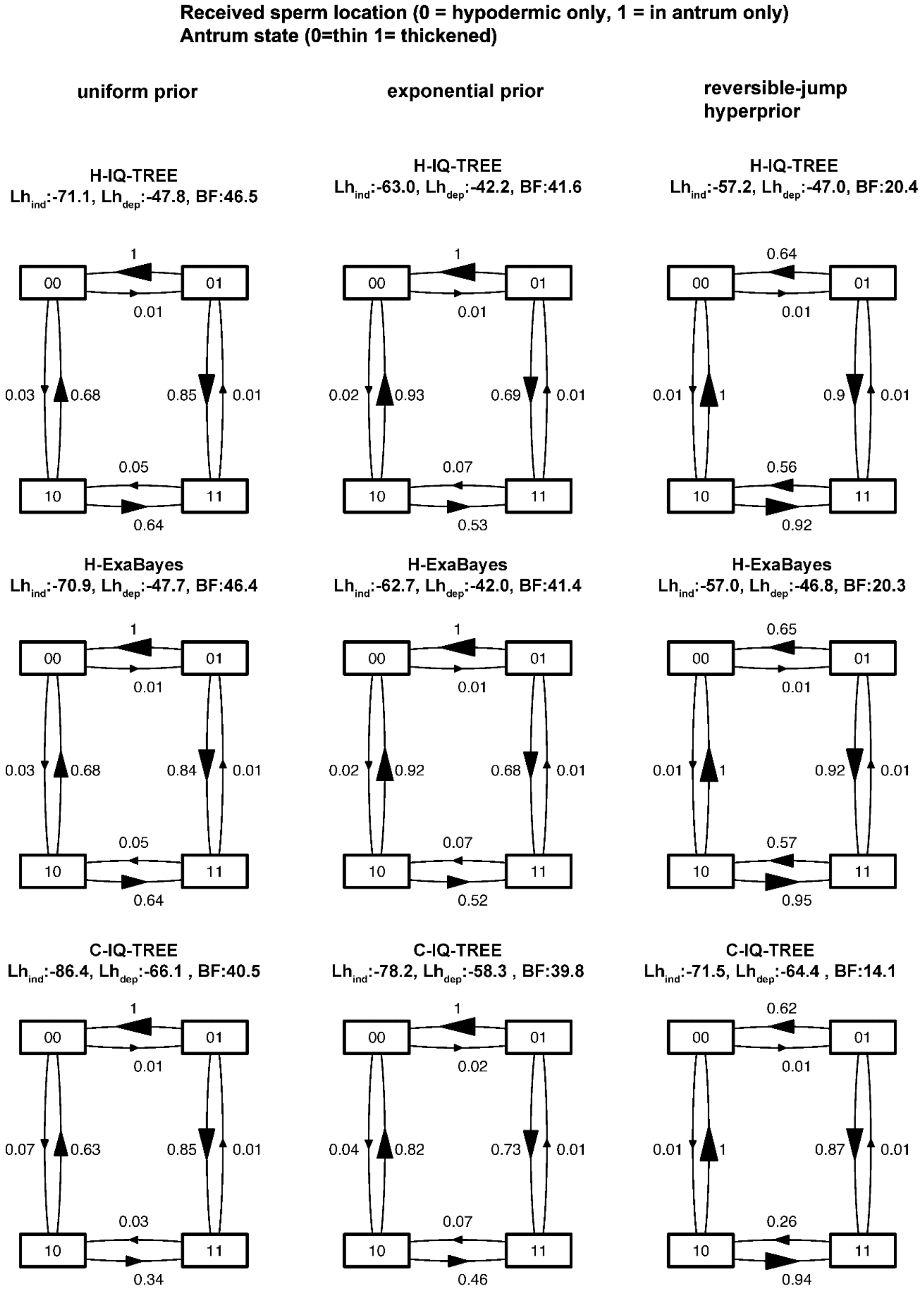

**Figure A7. Visualisation of the transition matrices for received sperm location and antrum state.** Transition rates are scaled so that the largest is unity. Rows show the different phylogenies and columns show different priors (and the bottom centre is what is shown in Figure 4B).

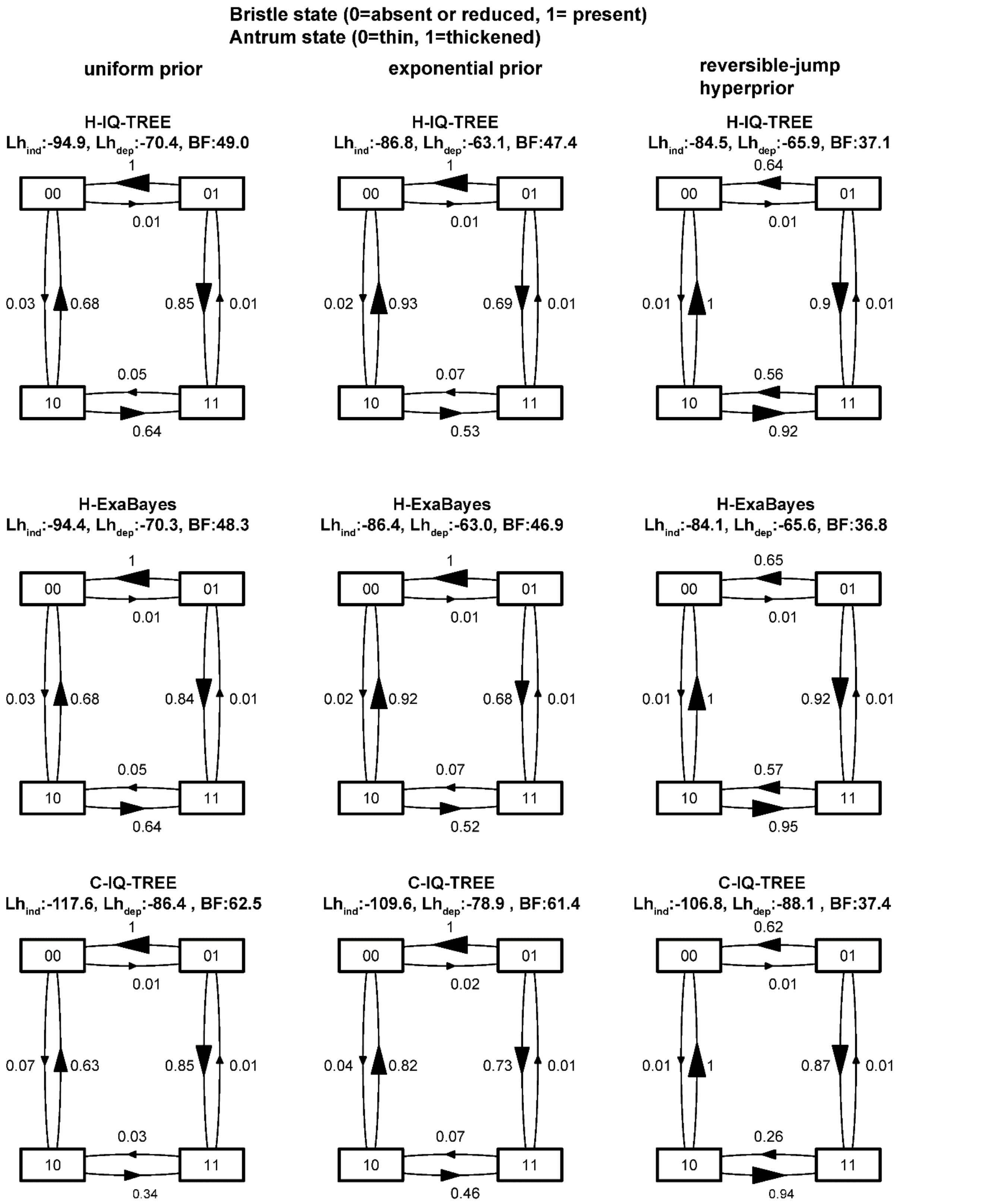

**Figure A8. Visualisation of the transition matrices for sperm bristle state and antrum state.** Transition rates are scaled so that the largest is unity. Rows show the different phylogenies and columns show different priors (and the bottom centre is what is shown in Figure 4C).

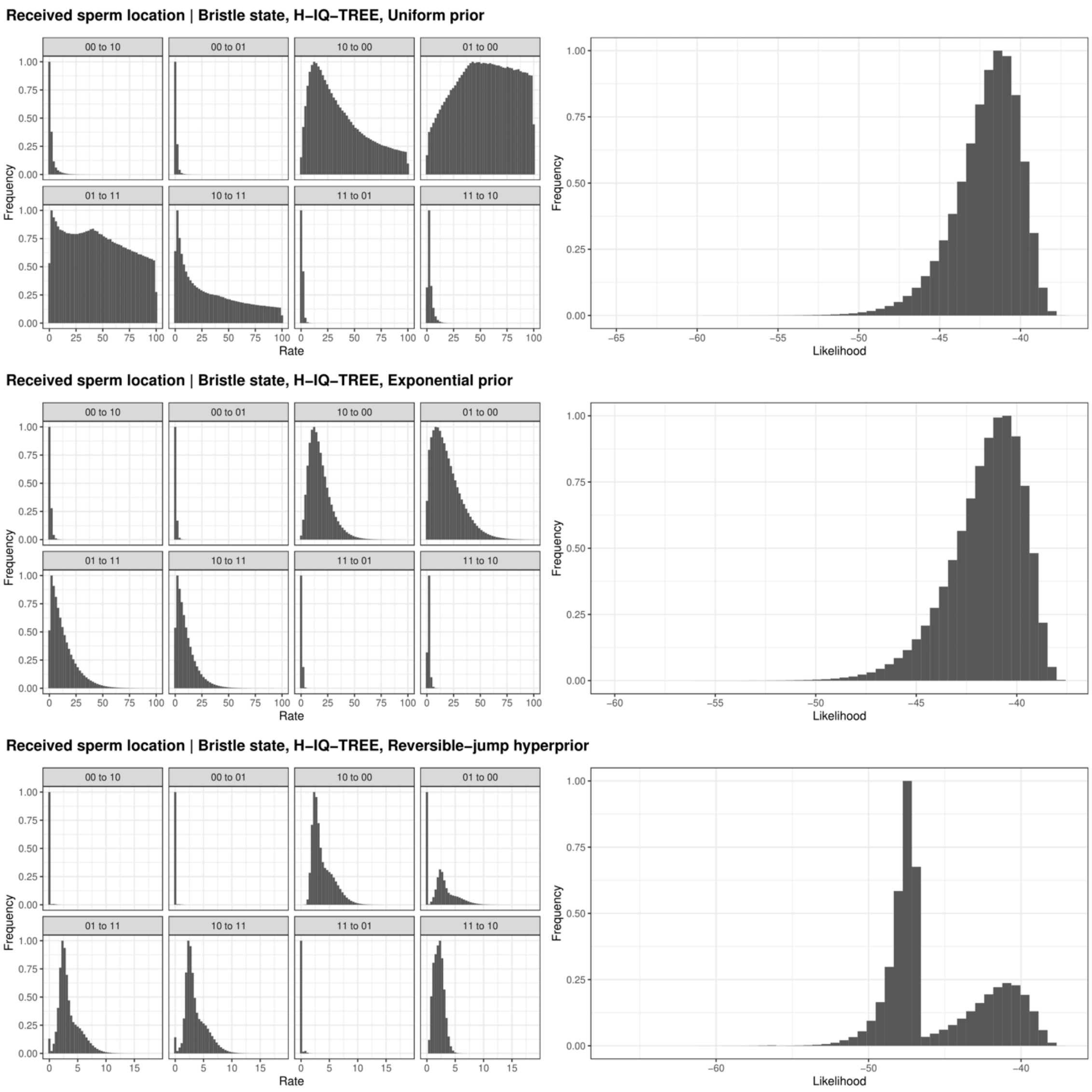

**Figure A9. Posterior distributions of the rate parameters (left) and likelihoods (right) for BayesTraits runs with the H-IQ-TREE phylogeny, and received sperm location and sperm bristle state.** Histograms show values from $2x10^6$ samples from four converged chains and are scaled so that the largest bin has a height of unity. Rate panels are arranged so that rates that would be equal in the independent model are arranged vertically (e.g. 00 to 10 would be equal to 01 to 11).

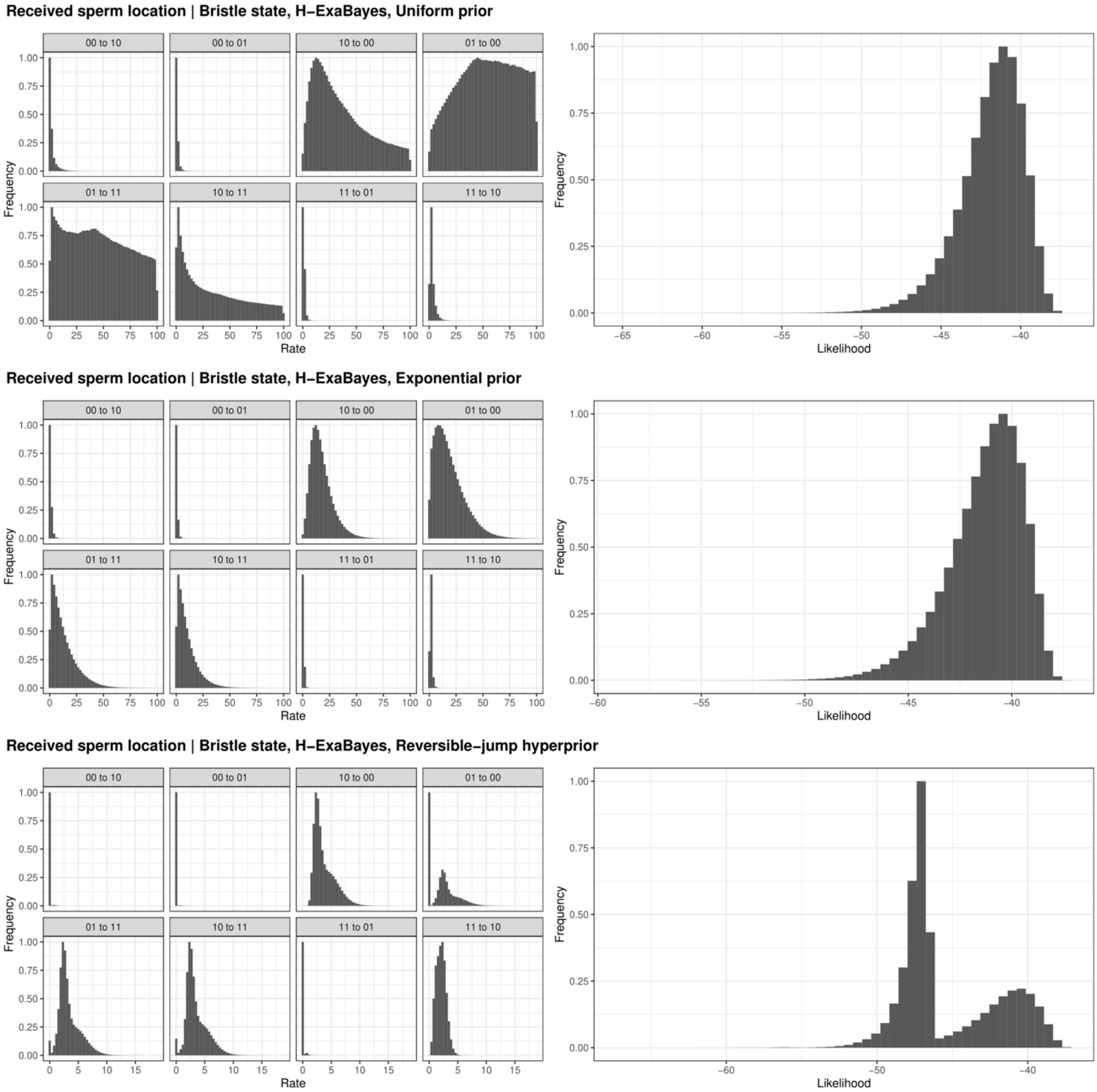

**Figure A10. Posterior distributions of the rate parameters (left) and likelihoods (right) for BayesTraits runs with the H-ExaBayes phylogeny, and received sperm location and sperm bristle state.** Layout identical to the previous figure.

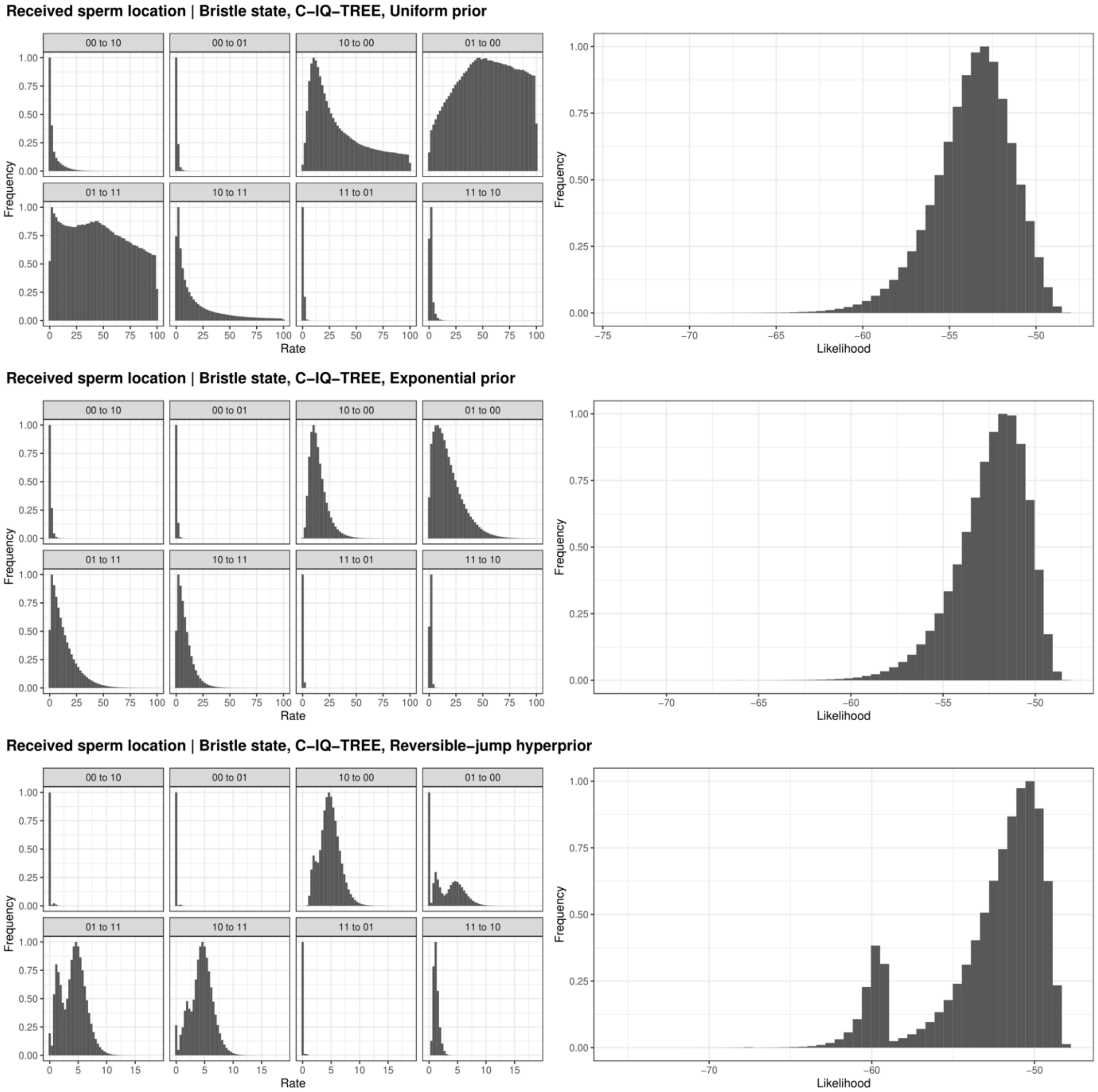

**Figure A11. Posterior distributions of the rate parameters (left) and likelihoods (right) for BayesTraits runs with the C-IQ-TREE phylogeny, and received sperm location and sperm bristle state.** Layout identical to the previous figure.

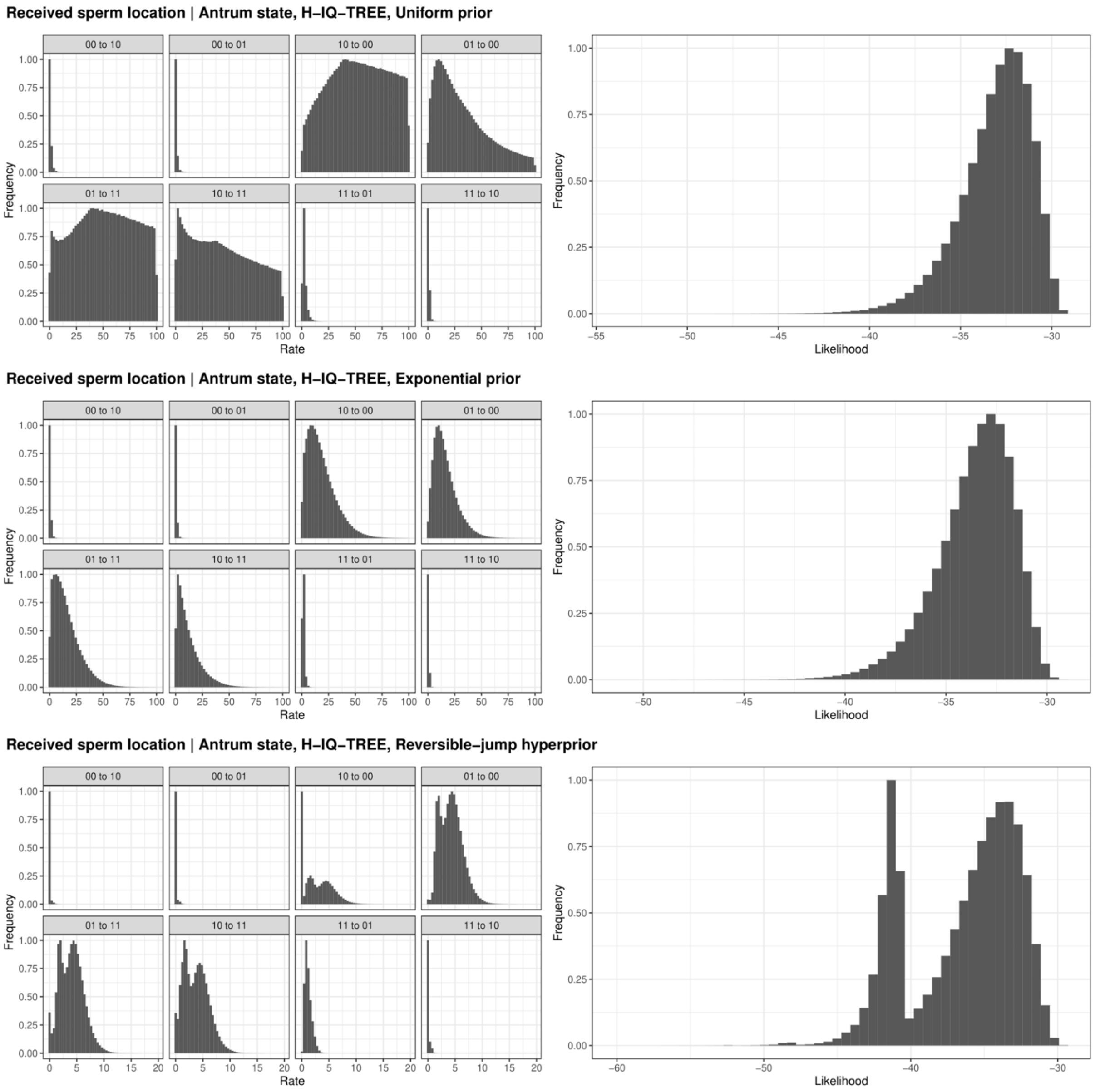

**Figure A12. Posterior distributions of the rate parameters (left) and likelihoods (right) for BayesTraits runs with the H-IQ-TREE phylogeny, and received sperm location and antrum state.** Layout identical to the previous figure.

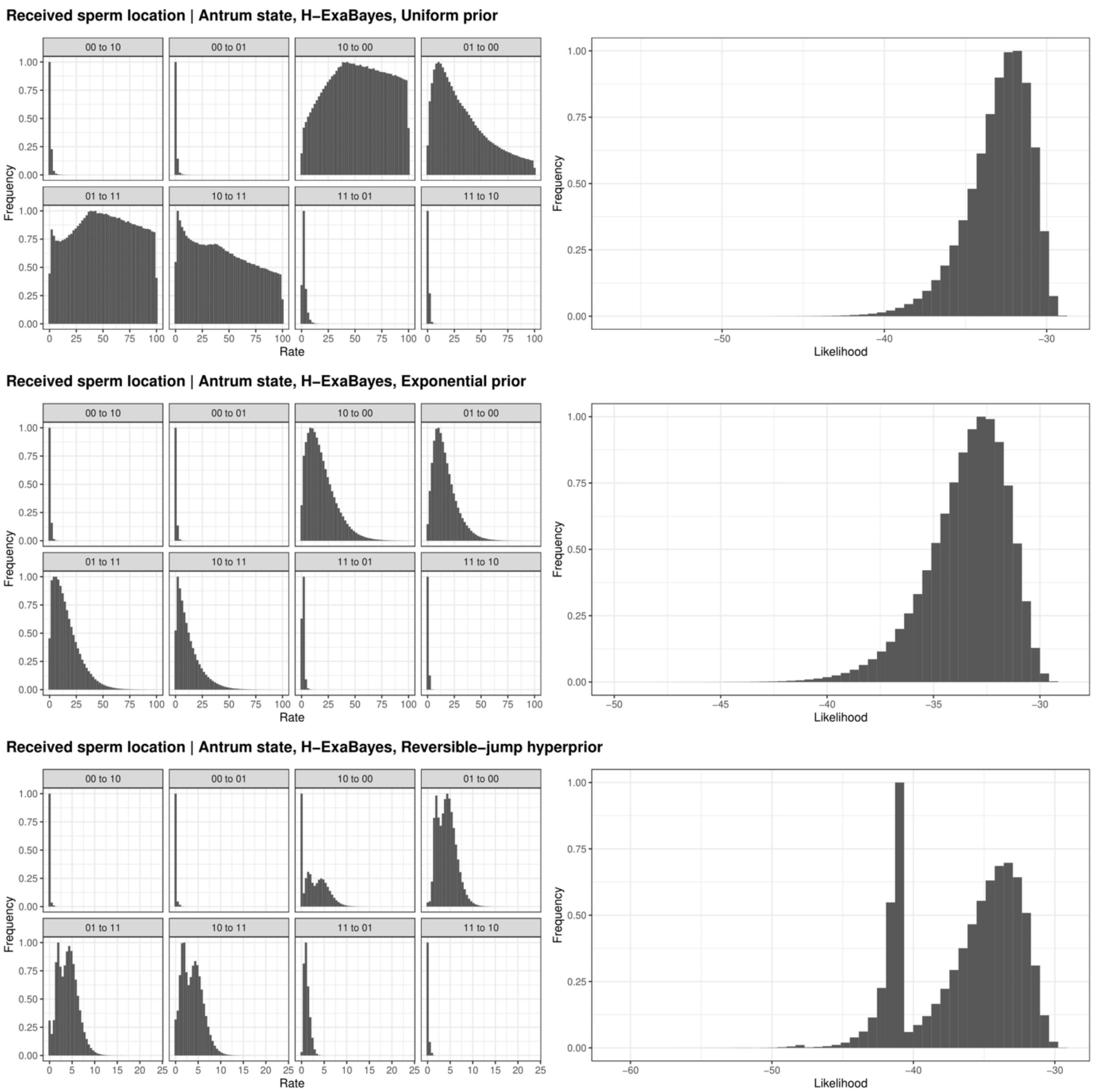

**Figure A13. Posterior distributions of the rate parameters (left) and likelihoods (right) for BayesTraits runs with the H-ExaBayes phylogeny, and received sperm location and antrum state.** Layout identical to the previous figure.

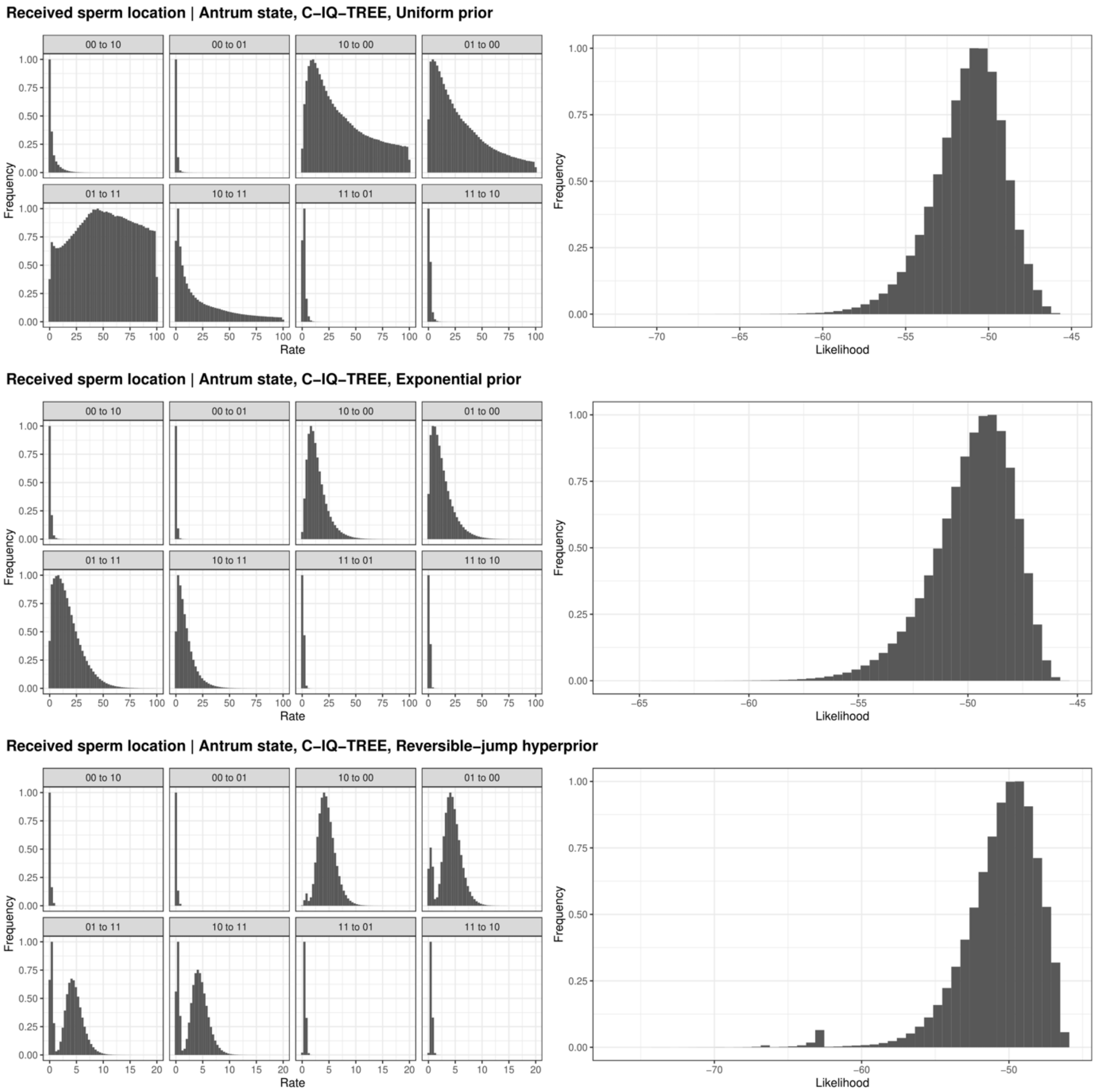

**Figure A14.1 Posterior distributions of the rate parameters (left) and likelihoods (right) for BayesTraits runs with the C-IQ-TREE phylogeny, and received sperm location and antrum state.** Layout identical to previous figure.

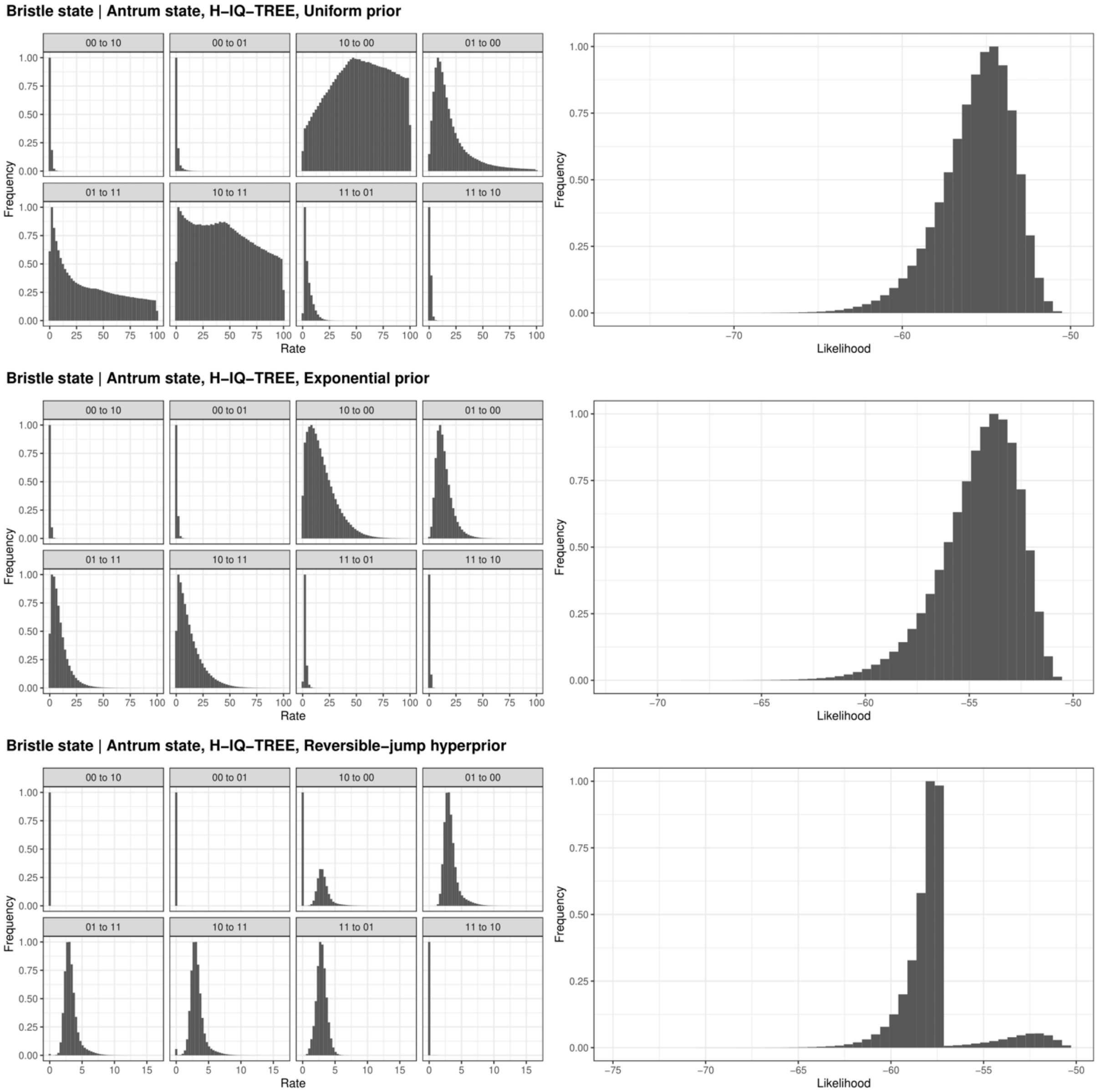

**Figure A15. Posterior distributions of the rate parameters (left) and likelihoods (right) for BayesTraits runs with the H-IQ-TREE phylogeny, and sperm bristle state and antrum state.** Layout identical to previous figure.

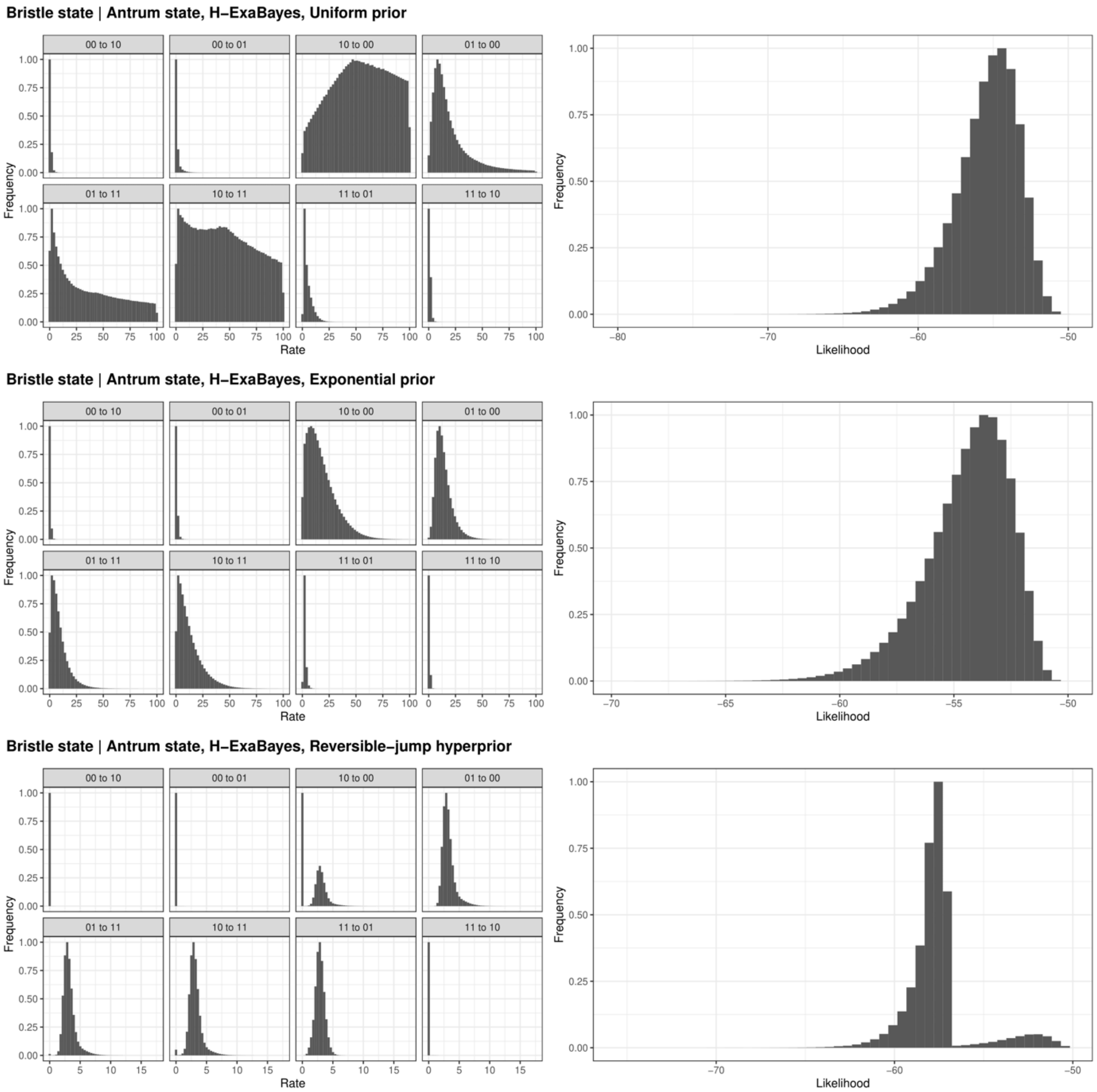

**Figure A16. Posterior distributions of the rate parameters (left) and likelihoods (right) for BayesTraits runs with the H-ExaBayes phylogeny, and sperm bristle state and antrum state.** Layout identical to previous figure.

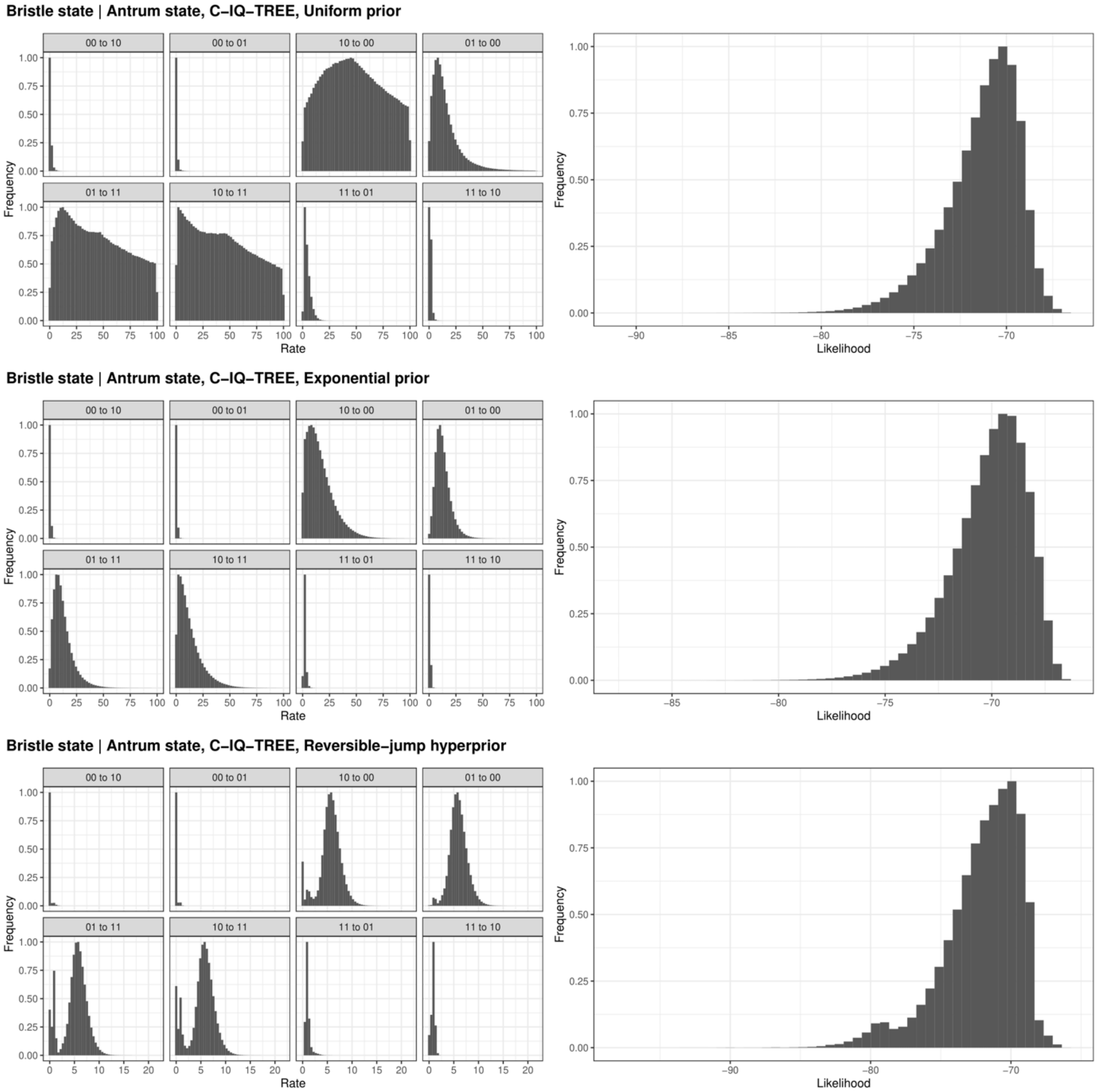

**Figure A17. Posterior distributions of the rate parameters (left) and likelihoods (right) for BayesTraits runs with the C-IQ-TREE phylogeny, and sperm bristle state and antrum state.** Layout identical to previous figure.

### SI Taxonomy

The analyses we present here have implications for the taxonomy and nomenclature of the Macrostomidae [15], and they support taxonomic and nomenclatural revisions. We outline these changes in the following, using the "*Genus species* Author, Year" format to refer to genera and binomials, and additionally including all the citations for the relevant works.

#### *Archimacrostomum* and *Inframacrostomum* should be dropped

As part of his monograph, Ferguson [16] established two new genera based on the absence of parts of the sexual system. On the one hand, the genus *Archimacrostomum* Ferguson, 1954 is characterised by an incomplete male sexual system, specifically a lack of the vesicula granulorum, a prominent structure that usually connects the muscular true seminal vesicle to the proximal stylet opening, and which contains complex muscles and serves as the entry point of the necks of the prostate gland cells into the stylet lumen. On the other hand, the genus *Inframacrostomum* Ferguson, 1954 is characterised by an incomplete female sexual system, specifically the lack of a discrete female antrum (which Ferguson refers to as the "female genital atrium"). Contemporary workers, namely Ax and Papi, strongly opposed the erection of these two genera [17, 18], primarily because "The absence of a specific character as a single trait is [in my view] completely unsuitable as a basis for the erection of independent genera."(our translation from [13]). These critical assessments were later also supported by Schärer et al. [10] and Schockaert [19].

Both Ax and Papi clarified that species assigned to *Archimacrostomum* do not lack the vesicula granulorum, but that it can sometimes simply be accommodated inside a wide proximal stylet base [17, 18]. Concerning the absence of a discrete female antrum in *Inframacrostomum*, both authors point out that this structure can be more or less prominent and thus cannot serve as grounds for a generic placement. These early considerations are underlined by the striking levels of convergent evolution revealed by Schärer et al. [10] and the current study, which clearly show that the antrum has been simplified multiple times independently and hence cannot serve as synapomorphy for these genera. The clear recommendations by Ax and Papi have not been followed uniformly, however, and both *Archimacrostomum* and *Inframacrostomum* still appear in both the literature (e.g. [20]) and in online databases (e.g. http://www.marinespecies.org/aphia.php?p=taxdetails&id=142052), although more often the former genus than the latter.

We here now formally place all the species that Ferguson moved to these genera back to their original combinations in the genus *Macrostomum*. From *Archimacrostomum* we thus reinstate *Macrostomum beaufortense* Ferguson, 1937 (originally named *Macrostomum beaufortensis* Ferguson, 1937) [21], *Macrostomum hustedi* Jones, 1944 [22], and *Macrostomum brasiliense* Marcus, 1952 (originally named *Macrostomum appendiculatum* forma *brasiliensis* Marcus, 1952) [23]. From *Inframacrostomum* we reinstate *Macrostomum rubrocinctum* Ax, 1951 [24]. Moreover, we also reassign a subsequently named species, namely from *Archimacrostomum,* we establish *Macrostomum sublitorale* (Faubel & Warwick, 2005) [20], and reinstate two additional species that were also transferred to *Archimacrostomum* by these authors, namely *M. pusillum* Ax, 1951 [24] and *M. peteraxi* Mack-Fira, 1971 [25].

### References

1. Janicke T, Schärer L. Sperm competition affects sex allocation but not sperm morphology in a flatworm. Behavioral Ecology and Sociobiology. 2010;64:1367–75.

2. Schärer L, Brand JN, Singh P, Zadesenets KS, Stelzer C-P, Viktorin G. A phylogenetically informed search for an alternative *Macrostomum* model species, with notes on taxonomy, mating behavior, karyology, and genome size. Journal of Zoological Systematics and Evolutionary Research. 2020;58:41–65.

3. Ladurner P, Schärer L, Salvenmoser W, Rieger RM. A new model organism among the lower Bilateria and the use of digital microscopy in taxonomy of meiobenthic Platyhelminthes: *Macrostomum lignano*, n. sp. (Rhabditophora, Macrostomorpha). Journal of Zoological Systematics and Evolutionary Research. 2005;43:114–26.

4. Xin F, Zhang S-Y, Shi Y-S, Wang L, Zhang Y, Wang A-T. *Macrostomum shenda* and *M. spiriger*, two new brackish-water species of *Macrostomum* (Platyhelminthes: Macrostomorpha) from China. Zootaxa. 2019;4603:105.

5. Ferguson FF. A monograph of the genus *Macrostomum* O. Schmidt 1848. Part IV. Zoologischer Anzeiger. 1939;128:188–205.

6. An-der-Lan H. Zur rhabdocoelen Turbellarienfauna des Ochridasees (Balkan). Sitzungsberichte d mathem-naturw Kl, Abt I, 148. 1939;5:196–254.

7. Brand JN, Viktorin G, Wiberg RAW, Beisel C, Schärer L. Large-scale phylogenomics of the genus *Macrostomum* (Platyhelminthes) reveals cryptic diversity and novel sexual traits. bioRxiv. 2021;2021/437366.

8. Sluys R. First representative of the order Macrostomida in Australia (Platyhelminthes, Macrostomidae). Records of the South Australian Museum. 1986;19:399–404.

9. Papi F. Sulle affinità morfologiche nella fam. Macrostomidae (Turbellaria). Bolletino di zoologia. 1950;17:461–8.

10. Schärer L, Littlewood DTJ, Waeschenbach A, Yoshida W, Vizoso DB. Mating behavior and the evolution of sperm design. Proceedings of the National Academy of Sciences. 2011;108:1490–5.

11. Lange R, Reinhardt K, Michiels NK, Anthes N. Functions, diversity, and evolution of traumatic mating: Function and evolution of traumatic mating. Biological Reviews. 2013;88:585–601.

12. Reinhardt K, Anthes N, Lange R. Copulatory wounding and traumatic insemination. Cold Spring Harb Perspect Biol. 2015;7:a017582.

13. Kamimura Y. Twin intromittent organs of *Drosophila* for traumatic insemination. Biology Letters. 2007;3:401–4.

14. Kamimura Y. Copulation anatomy of *Drosophila melanogaster* (Diptera: Drosophilidae): wound-making organs and their possible roles. Zoomorphology. 2010;129:163–74.

15. Tyler S, Schilling S, Hooge M, Bush L F. Turbellarian taxonomic database. 2006-2021.

16. Ferguson FF. Monograph of the Macrostomine worms of turbellaria. Transactions of the American Microscopical Society. 1954;73:137.

17. Ax P. Zur Systematik, Ökologie und Tiergeographie der Turbellarienfauna in den ponto-kaspischen Brackwassermeeren. Zool Jb Syst. 1959;87:43–187.

18. Papi F. Specie nuove o poco note del gen. *Macrostomum* (Turbellaria: Macrostomida) rinvenute in Italia. Monit Zool Ital. 1959;66:1–19.

19. Schockaert ER. Marine Macrostomorpha (Platyhelminthes, Rhabditophora) from the Algarve (Southern Portugal). Zootaxa. 2014;3872:577.

20. Faubel A, Warwick RM. The marine flora and fauna of the Isles of Scilly: Free-living Plathelminthes ('Turbellaria'). Journal of Natural History. 2005;39:1–45.

21. Ferguson FF. The morphology and taxonomy of *Macrostomum beaufortensis* n. sp. Zoologischer Anzeiger. 1937;120:230–5.

22. Jones AW. *Macrostomum hustedi* n. sp.; A morphological and cytological study of a rhabdocoel Turbellarian. Journal of Morphology. 1944;75:347–58.

23. Marcus E. Turbellaria Brasileiros (10). Bol Fac Fil Ciênc Letr Univ São Paulo Zool. 1952;17:5–187.

24. Ax P. Die Turbellarien des Eulitorals der Kieler Bucht. Zoologische Jahrbücher: Abteilung für Systematik, Ökologie und Geographie der Tiere. 1951;80:277–378.

25. Mack-Fira V. Deux turbellariés nouveaux de la mer noire. Rev Roum Biol Zool. 1971;16:233–40.
