## Supplementary figures and images for "Frequent origins of traumatic insemination involve convergent shifts in sperm and genital morphology"

### Figure S1

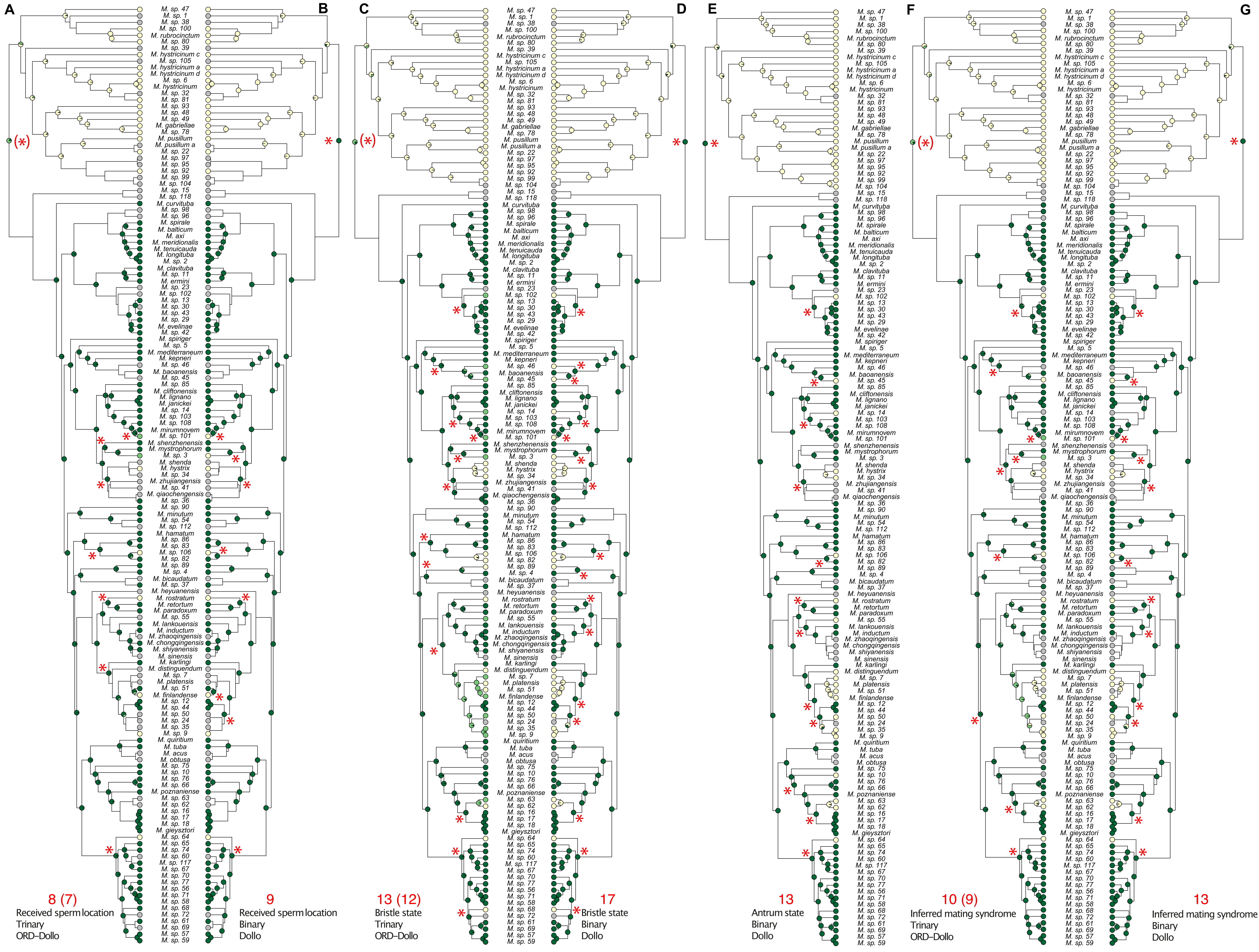

### Figure S2A

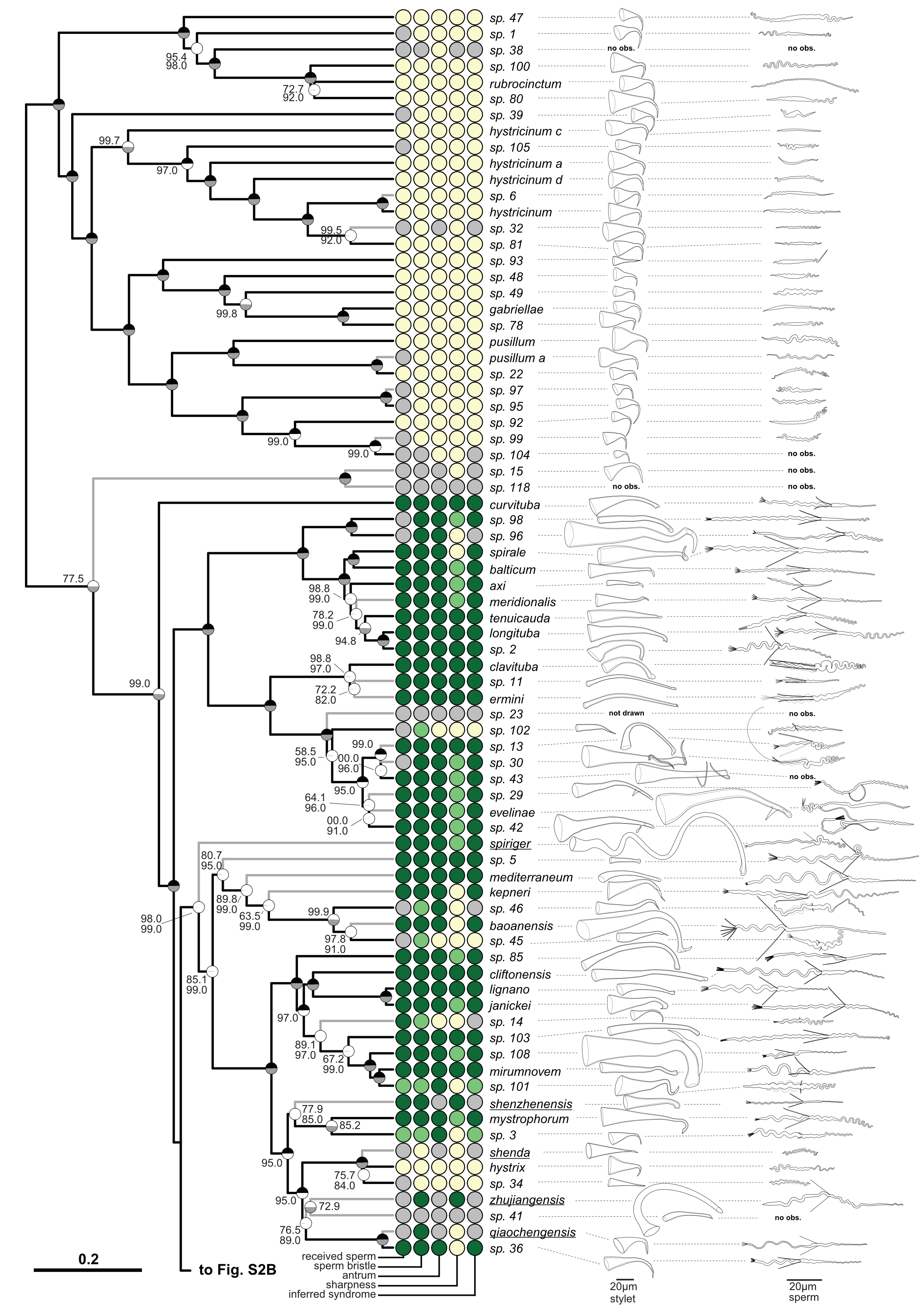

### Figure S2B

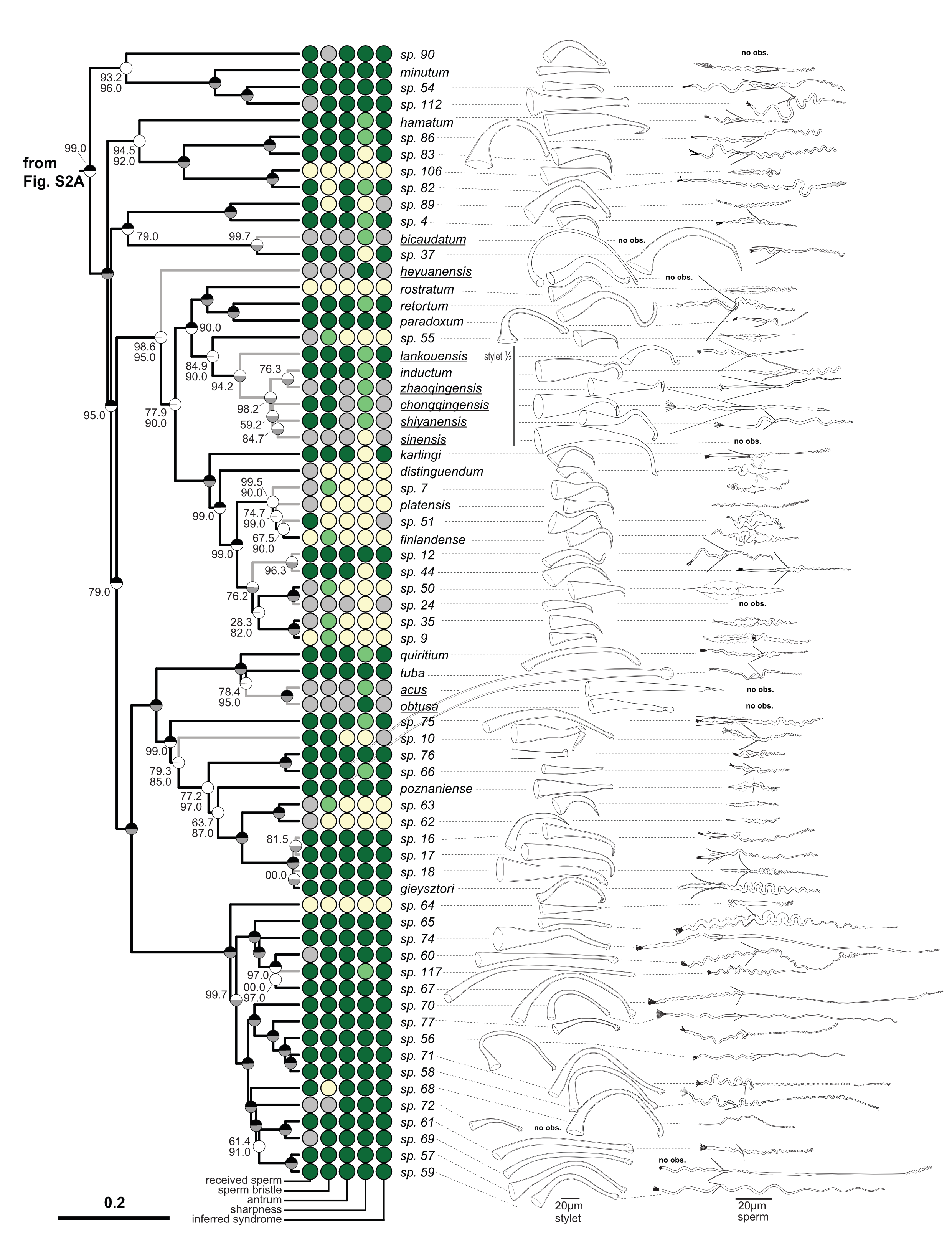

### Figure S3

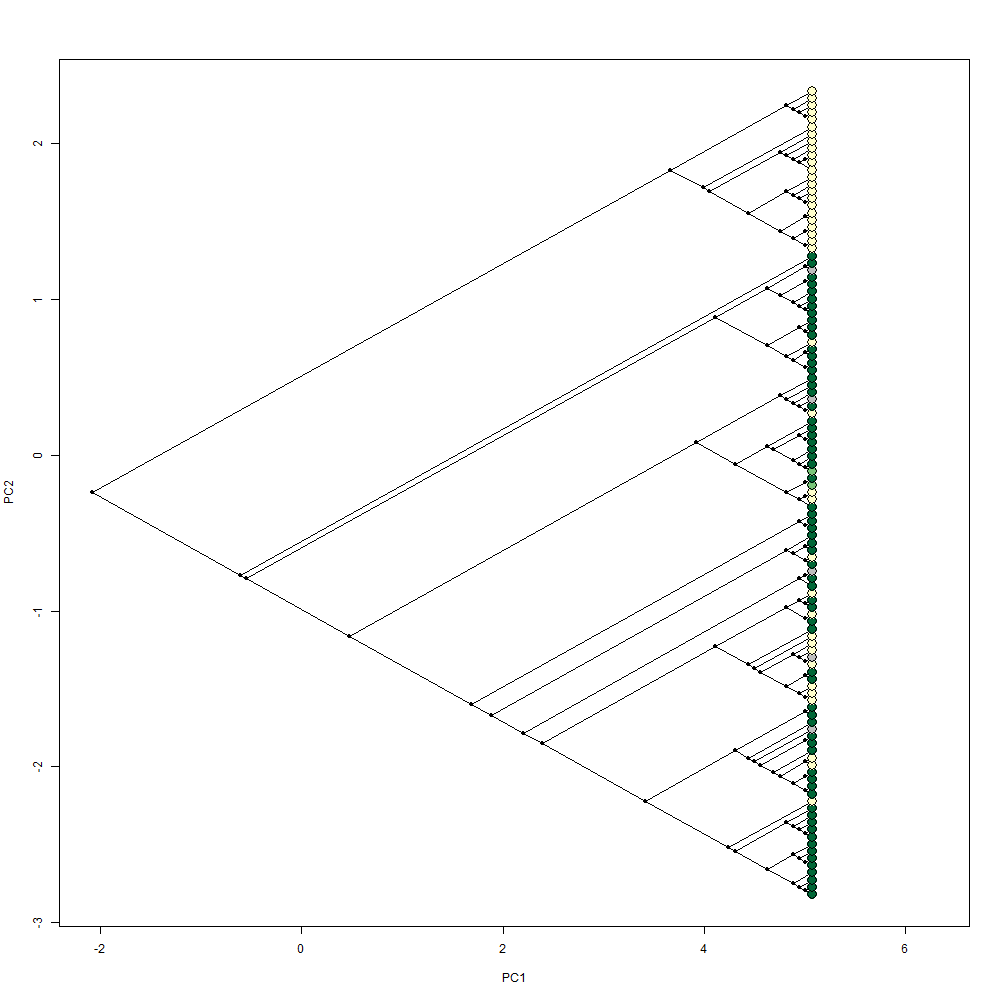

### Figure S4

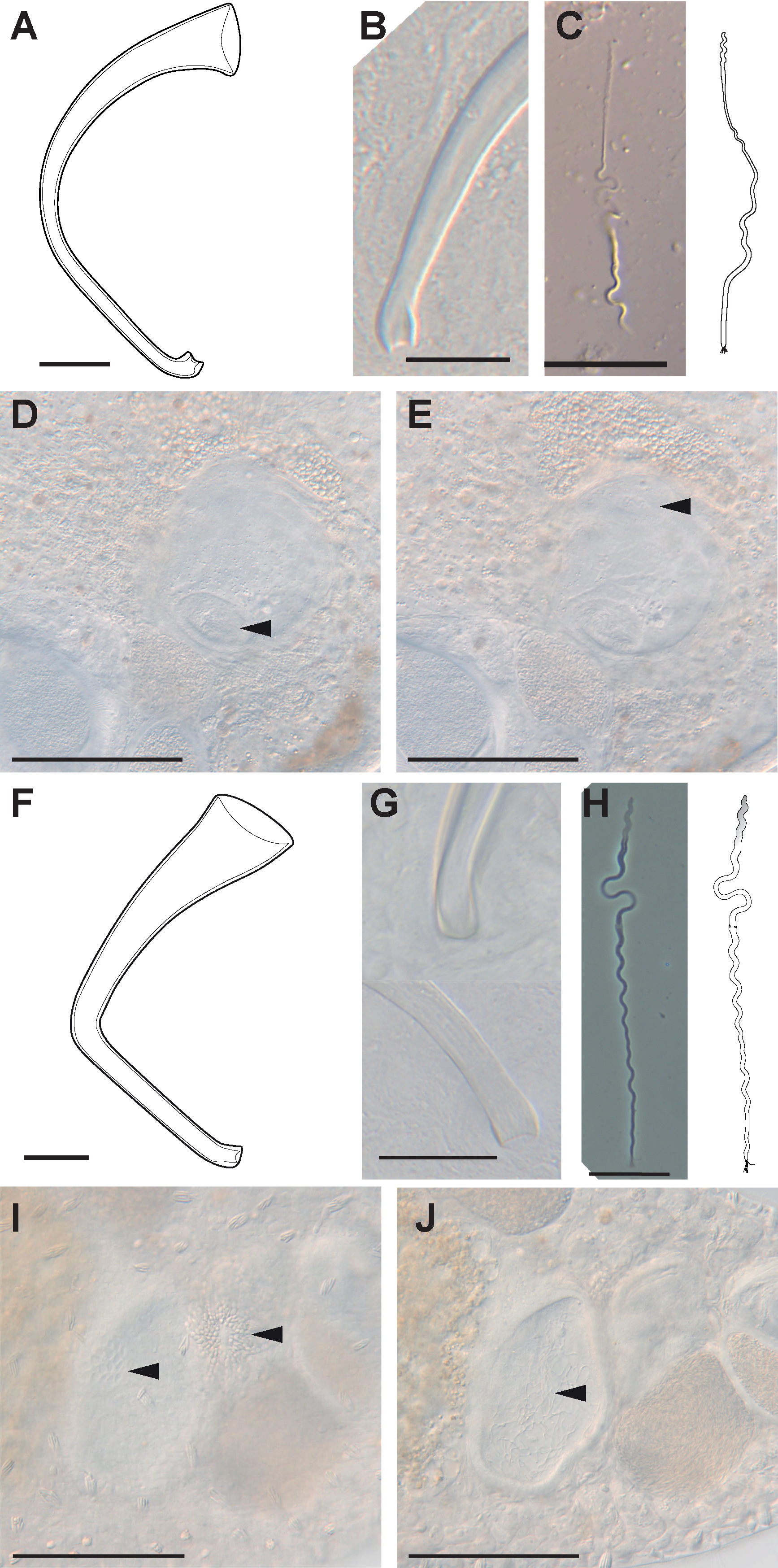

### Figure S5

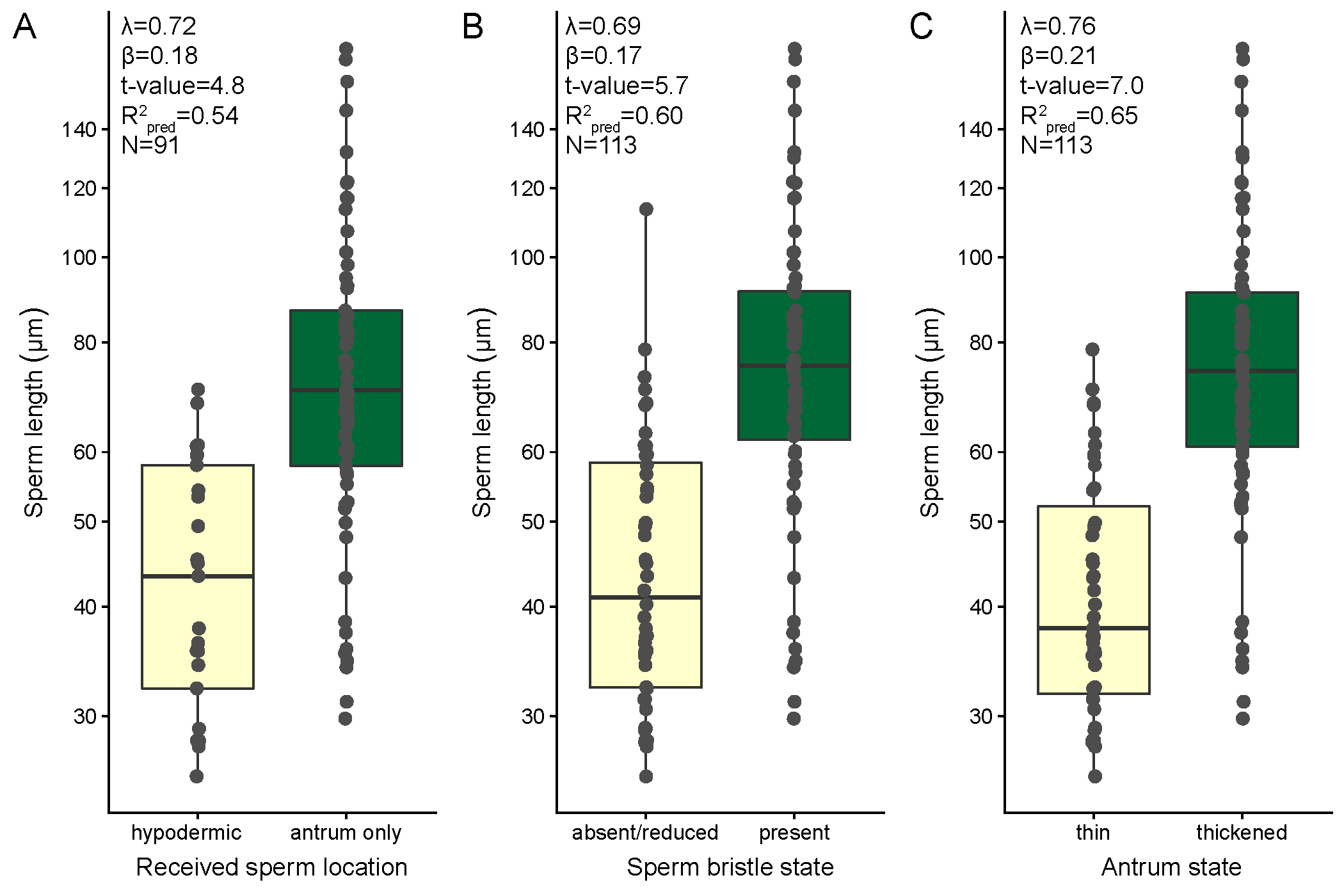
